# Allergic inflammation triggers dyslipidemia via IgG signalling

**DOI:** 10.1101/2023.08.04.551996

**Authors:** Nieves Fernández-Gallego, Raquel Castillo-González, Lucía Moreno-Serna, Antonio J. García-Cívico, Elisa Sánchez-Martínez, Celia López-Sanz, Ana Luiza Fontes, Lígia L. Pimentel, Ana Gradillas, David Obeso, René Neuhaus, Marta Ramírez-Huesca, Ignacio Ruiz-Fernández, Emilio Nuñez-Borque, Yolanda R. Carrasco, Borja Ibáñez, Pilar Martín, Carlos Blanco, Coral Barbas, Domingo Barber, Luis M. Rodríguez-Alcalá, Alma Villaseñor, Vanesa Esteban, Francisco Sánchez-Madrid, Rodrigo Jiménez-Saiz

## Abstract

**Background:** Allergic diseases begin early in life and are often chronic, thus creating an inflammatory environment that may precede or exacerbate other pathologies. In this regard, allergy has been associated to metabolic disorders and with a higher risk of cardiovascular disease, but the underlying mechanisms remain incompletely understood.

**Methods:** We used a murine model of allergy and atherosclerosis, different diets and sensitization methods, and cell-depleting strategies to ascertain the contribution of acute and late phase inflammation to dyslipidemia. Untargeted lipidomic analyses were applied to define the lipid fingerprint of allergic inflammation at different phases of allergic pathology. Expression of genes related to lipid metabolism was assessed in liver and adipose tissue at different times post-allergen challenge. Also, changes in serum triglycerides (TG) were evaluated in a group of 59 patients ≥14 days after the onset of an allergic reaction.

**Results:** We found that allergic inflammation induces a unique lipid signature that is characterized by increased serum TG and changes in the expression of genes related to lipid metabolism in liver and adipose tissue. Alterations in blood TGs following an allergic reaction are independent of T-cell-driven late phase inflammation. On the contrary, the IgG-mediated alternative pathway of anaphylaxis is sufficient to induce a TG increase and a unique lipid profile. Lastly, we demonstrated an increase in serum TG in 59 patients after undergoing an allergic reaction.

**Conclusion:** Overall, this study reveals that IgG-mediated allergic inflammation regulates lipid metabolism.

## INTRODUCTION

Type 2 immunity provides protection against parasites and non-microbial noxious substances and regulates several homeostatic processes^1,2^. However, when it is directed toward innocuous antigens from food, pollen, animal dander, *etc*. it causes allergic diseases^3^. These begin early in life and are often chronic, thus creating an inflammatory ambient that may precede or exacerbate other pathologies^4^. In this regard, allergy has been associated with a higher risk of atherosclerosis^5,6^, which accounts for over 80% of deaths caused by cardiovascular disease, the major cause of mortality worldwide^7^.

Atherosclerosis is a disease characterized by the chronic inflammation of blood vessels^8^. From an immunological perspective, atherosclerosis is predominantly driven by type 1 immunity, which is opposed to the hallmark type 2 identity of allergic disease^9^. Intriguingly, rather than counteracting atherosclerosis, allergy seems to exacerbate this condition^10^. Atherosclerosis begins early in life, typically associated with dyslipidemia, progresses slowly, and can remain silent until clinical manifestations arise in the form of ischemic heart disease and/or stroke^11,12^.

The pathology of allergic disease is heterogeneous and can generally be divided into two phases that are initiated by allergen exposure in a time-dependent manner. Upon allergen exposure, acute allergic reactions take place within minutes, or even seconds, via IgE (classical pathway)^13^ or, although less characterized in humans, via IgG (alternative pathway)^14,15^. Then, late phase inflammation develops hours, or even days, after allergen encounter, and it is mainly orchestrated by CD4 Th2 cells^16^.

Recent studies in human allergic patients point towards metabolic alterations linked to allergic disease^17–20^. Also, a recent transcriptomic study in the blood of healthy donors found an association between triglyceride (TG) levels and canonical genes of type 2 immunity^21^, which underscores the existence of an interplay between allergic disease and lipid metabolism. Yet the relationship of allergic disease and its different phases to metabolic changes and, particularly, to dyslipidemia, the main risk factor of atherosclerosis, remains poorly understood.

Here, we investigated the impact of allergic inflammation in dyslipidemia, and the effect of the diet, in a murine model of allergy and atherosclerosis. Furthermore, we ascertained the contribution of acute and late phase inflammation to allergic dyslipidemia using cell-depleting strategies and passive sensitization models and delved into the role of the classical and alternative anaphylaxis pathways. Moreover, untargeted lipidomics were applied to define the lipid fingerprint of allergic inflammation, and potential genes involved were studied in adipose tissue and liver. Lastly, we evaluated changes in serum TG levels in patients undergoing allergic reactions. Overall, this study reveals the importance of humoral allergic inflammation in lipid metabolism, which may contribute to atherosclerosis.

## RESULTS

### Allergic reactivity causes a diet-independent increase in blood triglycerides

To investigate the clinical association between allergic disease and atherosclerosis, we applied a well-established model of allergy and anaphylaxis^22,23^ in LDL receptor-deficient mice (LDLr KO) (**Fig. 1a**). This mouse strain is prone to atherosclerosis, particularly when fed a high cholesterol (HC) diet^24^. Following sensitization, LDLr KO mice on a HC diet developed high serum levels of specific IgE and IgG1 (**Fig. 1b**), and acute allergic responses upon allergen challenge, measured as hypothermia (**Fig. 1c**). Also, allergic mice exhibited late phase inflammation, characterized by eosinophilia in the peritoneal cavity (site of challenge), and a characteristic Th2-cytokine profile in the supernatants of spleen cell cultures (**Fig. 1d; Fig. S1**).

**Fig. 1.**
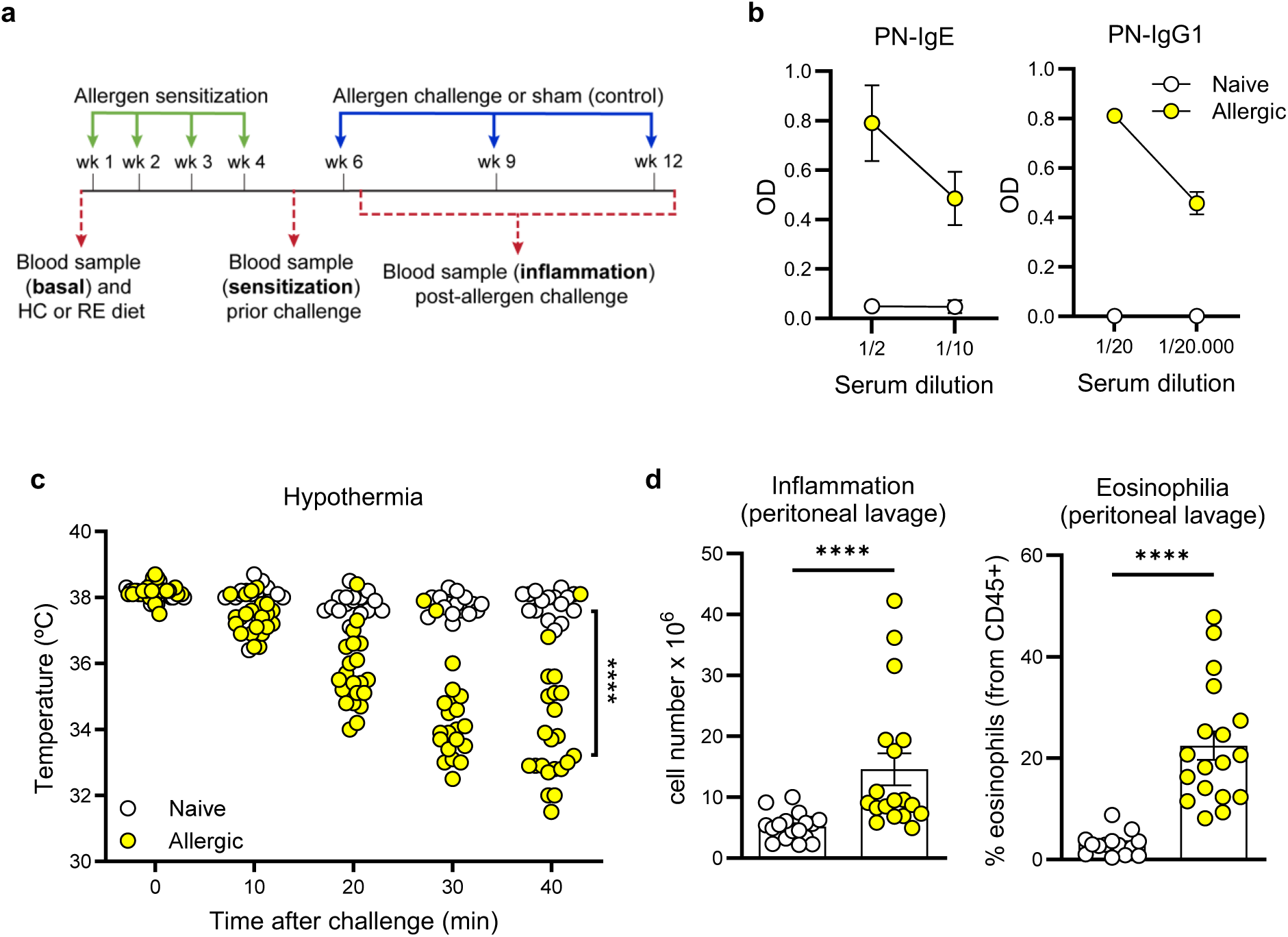
Allergic sensitization and clinical responses in atherosclerosis-prone mice. **a** Protocol for allergen sensitization, challenge, and sample collection in LDLr KO mice fed a high cholesterol (HC) or regular (RE) diet. **b** Serum levels of peanut (PN)-specific IgE and IgG1 after sensitization of HC diet-fed naive (n=14) and allergic (n=15) mice. **c** Acute allergic response following allergen challenge measured as hypothermia in HC diet-fed naive (n=16) and allergic mice (n=22). **d** Late phase inflammation in the peritoneal lavage at 3 days post-allergen challenge quantified in terms of total cell number and eosinophilia in HC diet-fed naive (n=15) or allergic (n=18) mice. Data for b and d are presented as mean ± s.e.m. In c and d, each dot represents an individual mouse. Statistically significant differences in c (at 40 min) and d were calculated with the Mann-Whitney test (****p≤0.0001). Pool of 3 independent experiments. We did not observe any significant differences in the lipid profile of allergic and naive LDLr KO mice on a HC diet except for TGs, which were increased in the allergic group, but only after allergen challenge (Fig. 2a).

Given that the blood lipid profile can inform of cardiovascular risk, we collected fasting blood samples at different time points of allergic pathology including baseline, after sensitization, and post-allergen challenge (**Fig. 1a**).

The fact that elevated TG levels have been recently identified as a risk factor of subclinical atherosclerosis in the PESA study (an observational, longitudinal and prospective study with 4,000 patients), even in patients with normal LDL/HDL^25^, prompted us to investigate these changes further. We questioned whether the systemic TG increase observed in allergic mice following allergen exposure was partly due to the HC diet used to study atherosclerosis or was inherent to allergic inflammation. To address this question, we followed a similar experimental design (**Fig. 1a**), but mice were fed a regular diet (RE) rather than a HC one. Also, we collected blood samples after the first challenge, to assess lipid changes in the incipient stages of atherosclerosis. In agreement with the previous data, we did not observe differences in the lipid profile of allergic and naive mice at different phases of allergic pathology, except for TGs, which were again increased in the allergic group after allergen challenge (**Fig. 2b**). These findings were substantiated in wild type, allergic animals, on a HC diet (**Fig. S2**). Lastly, we inquired about the source of the TG released during allergic inflammation by determining the expression of several genes related to lipid metabolism in the liver and in abdominal fat 2- and 4-days following allergen challenge (**Fig. 2c; Fig. S3).** We found significant changes in *Fasn, Dgat2, Elovl6, Lipin1* and *Srebf1.* Of these, *Lipin1* consistently increased in both adipose tissue and liver, while *Srebf1* decreased in both (**Fig. 2c**). Altogether, these results show that allergic inflammation induces a systemic TG increase in atherosclerosis-prone mice that is diet independent and partly related to an altered expression of *Lipin* and *Srebf1* in liver and adipose tissue.

**Fig. 2.**
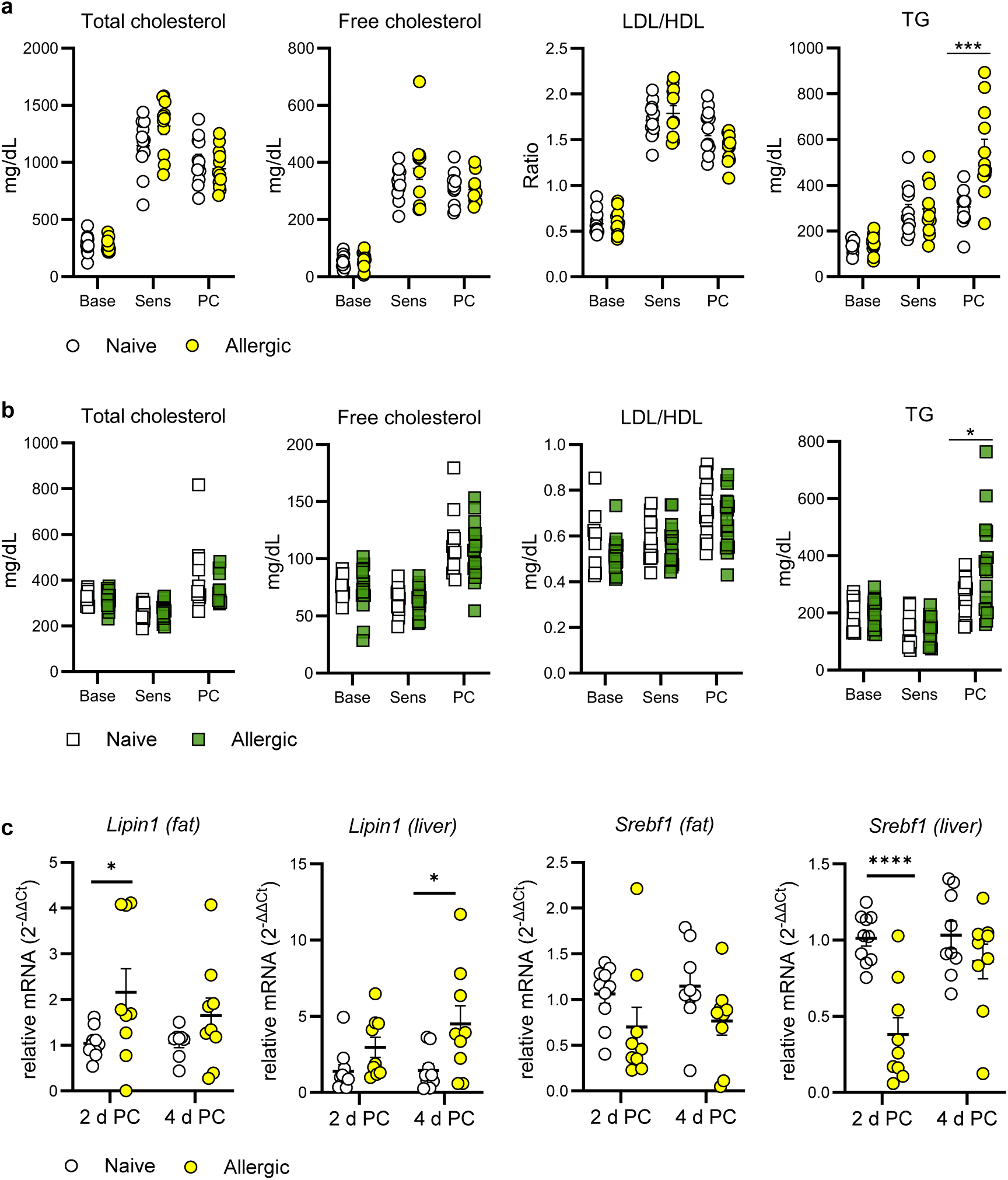
Serum triglyceride (TG) levels increase following allergic inflammation and are associated with changes in gene expression. **a, b** Serum levels of total cholesterol, free cholesterol, LDL/HDL ratio and TGs determined at baseline (Base), after sensitization (Sens) and 3 days post-allergen challenge (PC) in naive and allergic mice fed a high-cholesterol (HC) diet **(a)** or a regular (RE) diet **(b)**. Relative mRNA expression of *Lipin1* and *Srebf1* in abdominal fat and liver of naive and allergic mice on a HC diet at 2- and 4-days PC **(c)**. The data shown in **a** include 12 naive and 12 allergic mice; the data shown in **b** include 15 naive and 17 allergic mice; the data shown in **c** include 10 naive and 9 allergic mice; each dot represents an individual mouse. Data are presented as mean ± s.e.m. Statistically significant differences between groups for each time point were calculated with the Mann-Whitney test or the Student’s t-test according to data normality (*p≤0.05; ***p≤0.001; ****p≤0.0001). Pool of 2-3 independent experiments.

### Allergic inflammation causes a unique lipid signature in circulation

The systemic increase in serum TG levels following an allergic reaction was suggestive of more profound changes in the lipid profile. Thus, we performed a comprehensive, kinetic blood analysis at different times post-allergen (or PBS) exposure in allergic and naive mice under HC or RE diets, via untargeted lipidomics (**Fig. 3a**). The plasma samples, collected at 1-, 3-, and 7-days post-allergen or PBS challenge, were treated for lipid extraction and were analysed using liquid chromatography coupled to mass spectrometry (LC-MS). After curating the data with the respective quality assurance using the blanks, the presence in samples and quality controls (QCs), and according to the coefficient of variance of the QCs (<30%), we obtained 1,374 features including both positive and negative ionization modes.

**Fig. 3.**
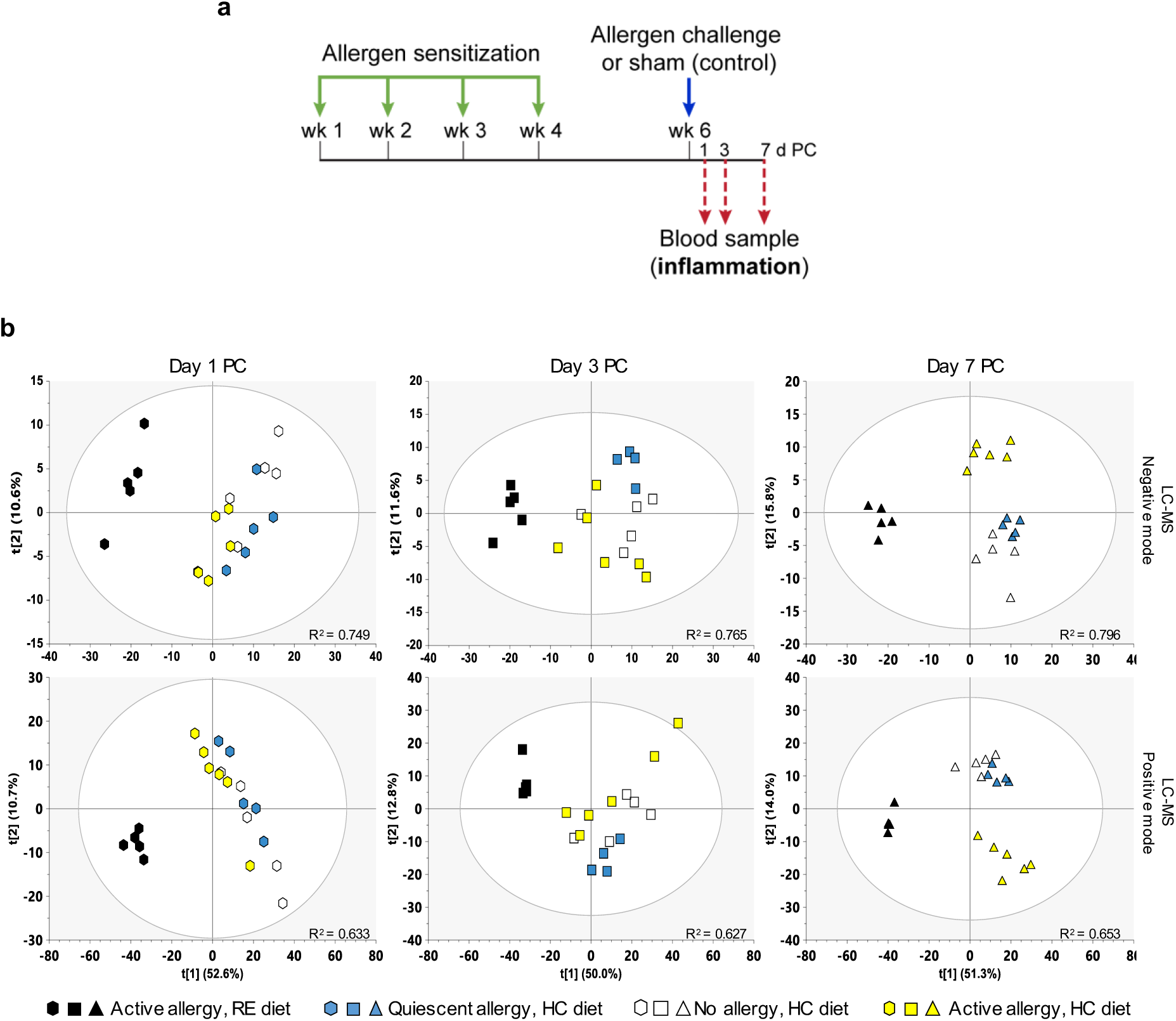
Principal component analysis (PCA) models of the blood lipid profile at different phases of allergic pathology. **a** Protocol for allergen sensitization, challenge, and kinetic of blood sample collection in LDLr KO mice fed a high cholesterol (HC) or regular (RE) diet for lipidomics. **b** Unsupervised PCA models of liquid chromatography coupled to mass spectrometry (LC-MS) in negative and positive ionization modes of plasma samples from allergic or naive mice at different time points (1-, 3-, and 7-days post-allergen challenge, PC) of allergic pathology. The analysis includes allergen-challenged allergic mice on a RE diet (n=5, black color); non-challenged allergic mice on a HC diet (n=5, blue color); allergen-challenged naive mice on a HC diet (n=5, white color); and allergen-challenged allergic mice on a HC diet (n=6, yellow color). All data were unit variance (UV) scaled. The X and Y axis indicate the percentage of variability explained by each component. R^2^ represents the total variability explained by the model to differentiate groups.

Then, we assessed for the major separations between groups based on the overall lipid profile via principal component analysis (PCA) models (**Fig. 3b**). As expected, the group on a RE diet differentiated well from those on a HC diet at all time points. Interestingly, the lipid profile of naive and PBS-challenged allergic mice on a HC diet was comparable and overlapping across the study, which indicates that quiescent allergy, or atopy *per se*, does not alter the blood lipid profile. Intriguingly, upon allergen challenge, the lipid profile of HC-diet-fed allergic mice became increasingly different, as compared to that of naive mice or unchallenged allergic mice on a HC diet. This effect became most obvious 7 days after inducing allergic inflammation, at which point the lipid profile of allergen-exposed mice on a HC diet was clearly distinct from that of the rest of the groups (**Fig. 3b).**

In view of the unique lipid fingerprint exhibited by allergic mice 7 days after allergen exposure, we decided to compare it to the lipid profile of mice with quiescent allergy (sensitized but not exposed to allergen), to better understand the impact of allergic inflammation on the lipid profile. We detected 392 features that were significantly different in this comparison (adjusted p-value, p-false discovery rate (FDR)< 0.05) including those from positive and negative ionization mode. Thus, from these, we identified 171 lipid species **(Fig. S4; Table S1)**. Among them, we detected 25 TG species, all of which were increased during allergic inflammation **(Fig. 4a)**. To characterize these TGs in detail, a closer inspection of the degree of unsaturation was performed revealing subclass-specific differences. TGs with saturated fatty acids were the most prominent group (40%), followed by those containing more than two double bonds (28%), with two double bonds (20%), and one unsaturation (12%). All these TGs were further analyzed using lipid metabolic networks (LINEX) according to the lengths and degree of unsaturation of the fatty acyl chain, and level of significance (**Fig. 4b**). The analysis showed a cluster of TGs with two double bonds in their sum composition that have been associated with cardiovascular disease (TG 54:2; TG 52:2; TG 50:2; TG 48:2)^26,27^. In addition to these, but not clustered together, TGs containing more than 2 double bonds such as TG 56:6, TG 56:4 and TG 53:3 have been also linked to cardiovascular disease^26^ or risk factors of it (*i.e.*, hypertension, diabetes)^28^. Collectively, these results show that allergic inflammation yields a unique lipid signature, characterized by TG changes, that extends beyond the acute phase of the pathology.

**Fig. 4.**
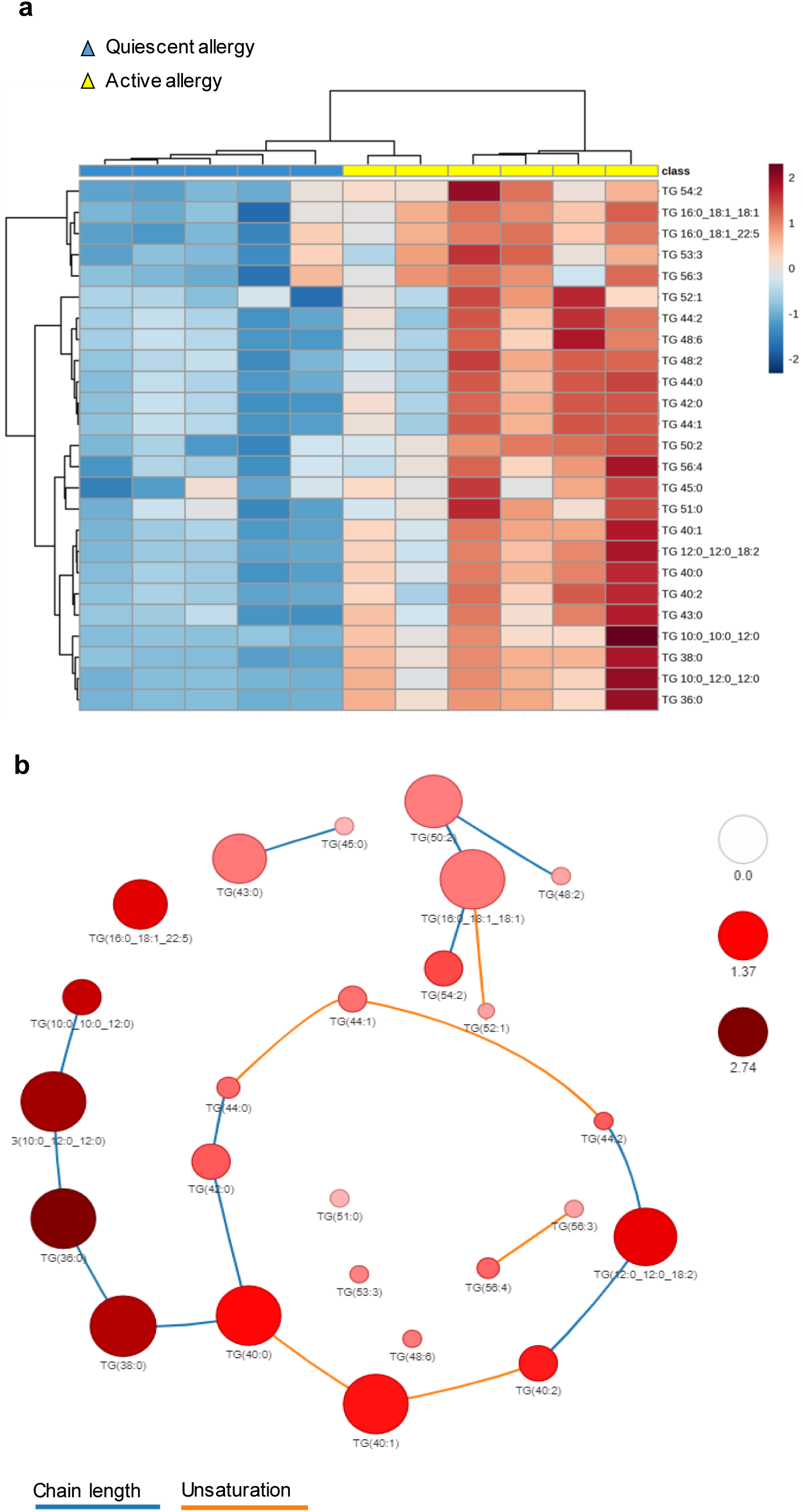
Allergic inflammation induces a unique blood lipid profile of triglycerides (TGs). **a** Heatmap of the 25 TG species differentially altered (p<0.05) between allergic mice with quiescent allergy (non-challenged, blue color, n=5) and active allergy (allergen-challenged, yellow color, n=6), fed a high cholesterol diet, at 7 days post-allergen challenge. The figure shows the clustering results in the form of a dendrogram. Relative lipid abundance is represented along a color gradient from blue (diminished in active allergy) to red (increased in active allergy). **b** Analysis of the lipid metabolic networks of differentially altered TGs in allergic mice with active allergy according to the lengths and degree of unsaturation of the fatty acyl chain, and level of significance with LINEX. The intensity of the color indicates the extent of TG fold change; the sphere size indicates the level of significance of the change according to the -log10 of the p FDR value; the association between TGs according to their length and degree of unsaturation are indicated by a blue and an orange line, respectively. TG 52:2 (TG 16:0_18:1_18:1); TG 56:6 (TG 16:0_18:1_22:5); FDR, false discovery rate.

### Blood changes in triglyceride levels are independent of T-cell driven allergic inflammation

Allergic inflammation is characterized by an acute phase, which happens within minutes, or seconds, following allergen exposure, and by a late phase, that occurs hours or even days after the reaction^16^. The fact that the unique blood lipid signature triggered by allergic inflammation was detected days after allergen exposure made us postulate that late phase inflammation was driving it. CD4 Th2 cells orchestrate late phase inflammation, hence we decided to deplete CD4 T cells prior allergen challenge to block it (**Fig. 5a**). We confirmed by flow cytometry that the antibody regime followed depleted CD4 T cells (**Fig. 5b; Fig. S5**), as reported^22,29^. CD4 T-cell depletion began 2 weeks after sensitization (1 week before allergen challenge) to avoid interfering with the generation of specific IgE/IgG1, both of which contribute to the acute phase. Anti-CD4-depleted and isotype-treated allergic mice showed comparable levels of specific IgE and IgG1 (**Fig. 5c**). Likewise, both groups of allergic mice underwent anaphylaxis following challenge (**Fig. 5d**), thus confirming that CD4 T-cell depletion had neither interfered with sensitization nor with acute allergic responses. The analysis of the serum lipid profile revealed that the TG peak previously observed during allergic inflammation remained intact despite CD4 T-cell depletion (**Fig. 5e**). These data suggest that T-cell-driven late phase inflammation is not involved in the TG changes triggered by allergic inflammation.

**Fig. 5.**
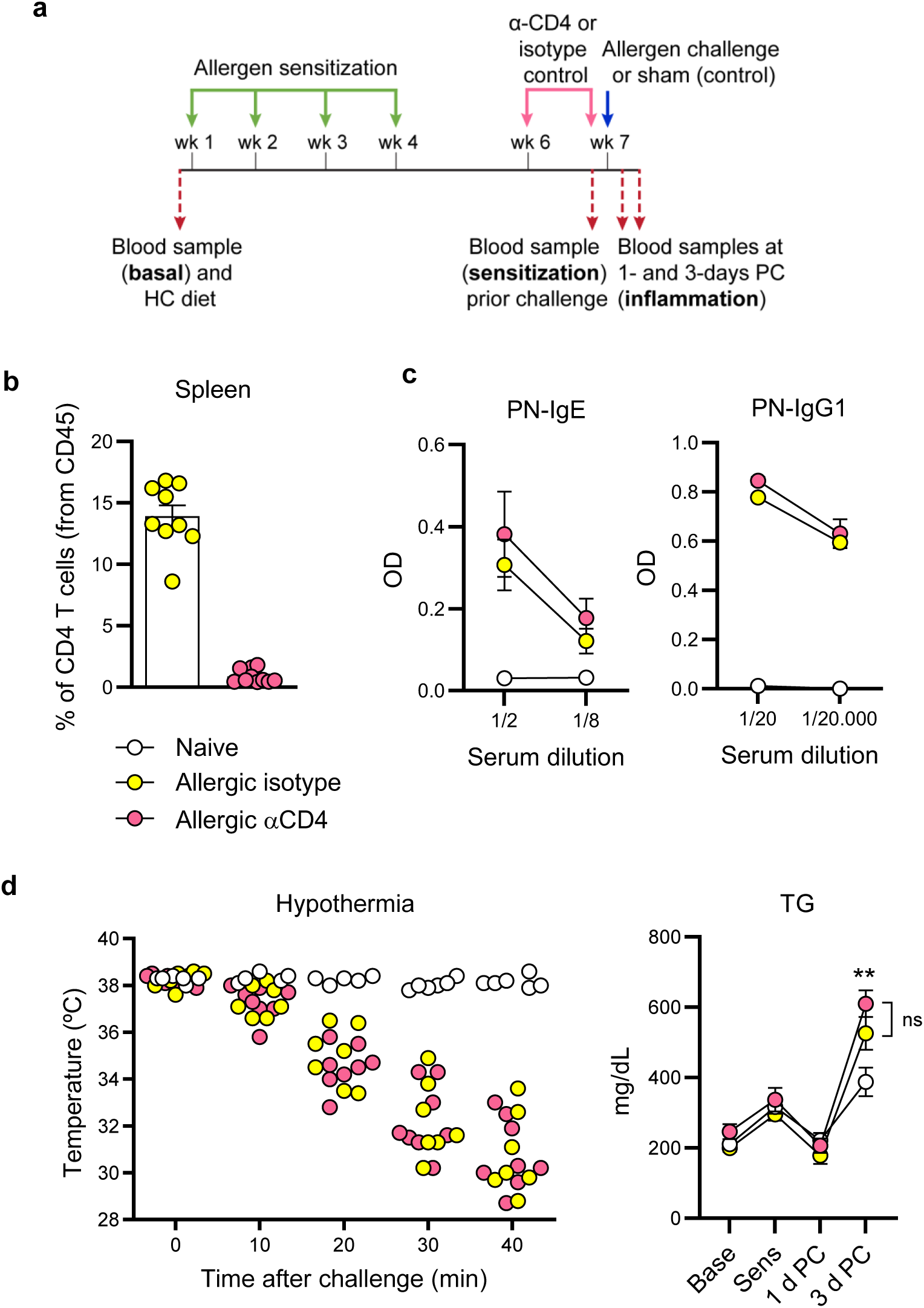
Blood changes in triglyceride (TG) levels during allergic inflammation are independent of CD4 T cells. **a** Protocol for allergen sensitization, CD4 T cell depletion, allergen challenge, and sample collection in LDLr KO mice fed a high cholesterol diet. **b** Flow cytometry analysis of CD4 T cell percentage in the spleen of anti-CD4-depleted and isotype-treated allergic mice. **c** Serum levels of peanut (PN)-specific IgE and IgG1 after sensitization. **d** Acute allergic responses following allergen challenge. **e** Serum TG levels determined at baseline (Base), after sensitization (Sens) and 1- and 3-days post-allergen challenge (PC) in naive (n=8), anti-CD4-depleted-(n=8) and isotype-treated (n=7) allergic mice. Data for b, c and e are presented as mean ± s.e.m. In b and d, each dot represents an individual mouse. Statistically significant differences in e were calculated with the Kruskal-Wallis test (**p≤0.001). Pool of 2 independent experiments.

### Blood changes in triglyceride levels during allergic inflammation are driven by IgG signalling

The seemingly dispensable contribution of late phase allergic inflammation to dyslipidemia raised the possibility that this was triggered by the acute allergic response, which is strictly dependent on IgE and IgG1. To test this hypothesis, we devised a model of passive sensitization, which is based on the transfer of sera from allergic mice into naive recipients^22^. With this approach, only the humoral allergic machinery is transferred to naive mice, thus allowing for intact acute allergic responses and precluding Th2-driven late phase inflammation (**Fig. 6a**). Given that acute allergic responses can operate through the classical (IgE-mediated) and/or the alternative pathway (IgG-mediated), we examined the contribution of each pathway to blood TG levels.

**Fig. 6.**
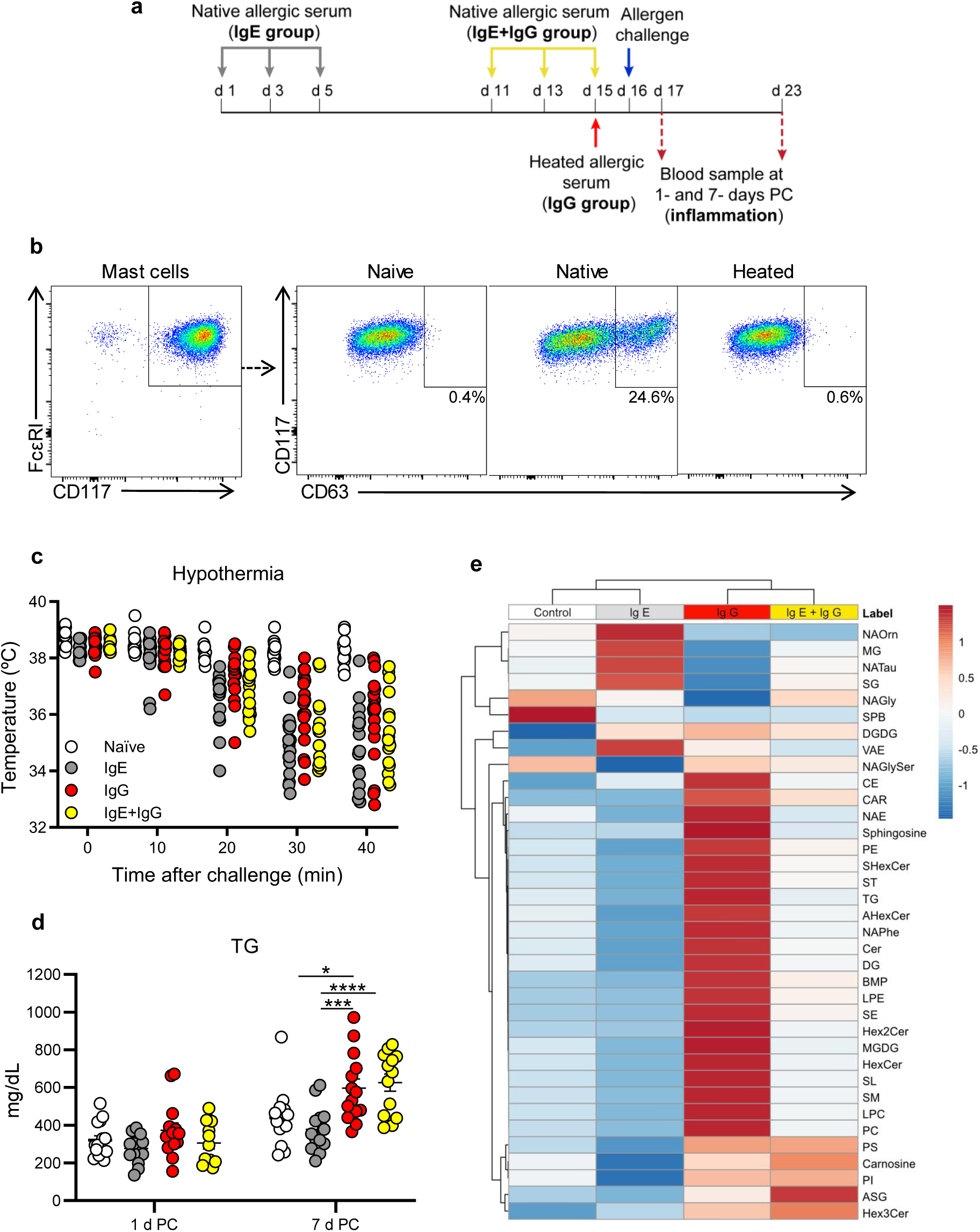
Blood changes in triglyceride (TG) levels and in the lipid profile are driven by the IgG-mediated alternative pathway of anaphylaxis. **a** Protocol for passive sensitization with native and heated sera to induce anaphylaxis via the classical and/or the alternative pathway in LDLr KO mice fed a high cholesterol diet. **b** Activation assay with mast cells sensitized with naive sera, native allergic sera, and heat-treated allergic sera (representative dot plots of 4 independent experiments). **c** Acute allergic responses following allergen challenge of mice passively sensitized with naive sera (n=17) or allergic sera (native or heated) to induce anaphylaxis via the classical (IgE; n=19) or alternative pathway (IgG; n=19), or both (IgE+IgG; n=18). **d** Serum TG levels measured at 1- and 7-days post-allergen challenge (PC) in passively sensitized mice. **e** Heatmap of the lipid species differentially altered (p-value<0.05) at 7 days PC in different groups of passively sensitized mice (a pool of 5 individual samples from each group was analyzed in triplicate). The figure shows the clustering results in the form of a dendrogram. Relative lipid abundance is represented along a color gradient from blue (diminished) to red (increased). Data for d are presented as mean ± s.e.m. In c and d each dot represents an individual mouse from a pool of 3 independent experiments. Statistically significant differences in d were calculated with a two-way-ANOVA followed by Bonferroni’s multiple comparisons test (*p≤0.05; ***p≤0.0005; ****p<0.0001). AHexCer, acylhexosylceramides; ASG, acylsterylglycoside; BMP, bis(monoacylglycero)phosphates; CAR, acyl carnitines; CE, cholesteryl esters; Cer, ceramides; DG, diacylglycerols; DGDG, digalactosyldiacylglycerols; Hex2Cer, dihexosylceramides; Hex3Cer, trihexosylceramides; HexCer, hexosylceramides; LPC, lysoglycerophosphocholines; LPE, lysoglycerophosphoethanolamines; MG, monoacylglycerols; MGDG, mongalactosyldiacylglycerols; NAE, N-acyl ethanolamines; NAGly, N-acyl glycines; NAGlySer, N-acyl glycyl serines; NAOrn, N-acyl ornithines; NAPhe, N-acyl phenylalanines; NATau, N-acyl taurines; PC, glycerophosphocholines; PE, glycerophosphoethanolamines; PI, glycerophosphoinositols; PS, glycerophosphoserines; SE, steryl esters; SG, sterylglycosides; SHexCer, sulfatides; SL, sulfonolipids; SM, sphingomyelins; SPB, sphingoid bases; ST, sterols; VAE, vitamin A fatty acid esters.

Unlike other antibodies, IgE is irreversibly denatured at 57°C due to a heat-labile domain, which is not present in other antibody isotypes^30,31^. We exploited this biochemical property to generate allergic sera without functional IgE. Following the heat-treatment of allergic sera, we observed a decrease in allergen-specific IgE levels as compared to the native allergic sera, while IgG1 levels remain unchanged (**Fig. S6a**). To ascertain if the remnants of denatured IgE were functional, we used a cell activation assay based on bone marrow-derived mast cells^32^. Mast cells sensitized with native allergic sera showed a dose-dependent activation following allergen stimulation by means of CD63 expression detected by flow cytometry (**Fig. 6b; Fig. S6b**). On the other hand, mast cells sensitized with heat-denatured allergic or naive sera did not increase CD63 expression after allergen stimulation, which was comparable to that of unstimulated cells (**Fig. 6b; Fig. S6b**). These data demonstrate that the heat-treatment rendered specific IgE completely ineffective.

We also leveraged on the ability of anaphylactic IgE to remain bound, for months, to mast cells via its high affinity receptor (FcεRI), which contrasts with the rapid clearance of IgG from the circulation in a matter of days^33^. Thus, resting the mice for 2 weeks after passive sensitization with allergic serum allows for blood IgG clearance and restricts anaphylaxis to the classical pathway^22^. Therefore, adjusting the sensitization time prior challenge as well as the type of sera used for it (native and heat-denatured), allowed us to induce anaphylaxis via the classical and/or alternative pathway (**Fig. 6a**). Upon challenge, all the allergic mice developed anaphylaxis to a similar level as determined by hypothermia (**Fig. 6c**), regardless of the sensitization sera used (native or heat-treated) and of the resting period prior challenge (1 day or 2 weeks). In line with the previous experiments, we did not observe changes between groups in the lipid profile following allergen challenge except for TGs (**Fig. 6d, Fig. S6c**). Interestingly, the group of allergic mice that underwent anaphylaxis through the classical pathway (IgE-mediated) did not exhibit a TG peak. However, the groups of mice that experienced anaphylaxis through the alternative pathway (IgG) or a combination of both, showed a significant increase in serum TGs (**Fig. 6d**). Furthermore, we determined the lipid profile of plasma from these mice at 7-days post-allergen challenge via LC-MS analysis, as mentioned above. We identified 1,198 features that were grouped into 36 lipid species for heatmap analysis (**Fig. 6e**). Remarkably, the naive and IgE groups were clustered together, while the IgE+IgG and IgG groups exhibited a profile that was more alike. The IgG group had a distinct lipid profile characterized by high peak intensities in TGs, as well as in lipids belonging to subclasses such as *N*-acyl ethanolamines (NAE), glycerophosphocholines (PC), lysoglycerophosphocholines (LPC), lysoglycerophosphoethanolamines (LPE), sphingosine (SPB), hexosylceramides (HexCer), glycosyldiacylglycerols (MGDG), sulfonolipids (SL), sphingomyelins (SM), ceramides (Cer) and diacylglycerols (DG). In line with this, the univariate analysis showed that 456 lipids changed significantly (p<0.05) when comparing the IgG *vs* control group, while 210 lipids did it in the IgE *vs* control group comparison (**Table S2**). Of note, 645 lipids were differentially expressed when comparing the IgG *vs* IgE group. These differences were attenuated when comparing the IgE+IgG group *vs* the IgE (239 lipids) or IgG (325 lipids) groups, due to their complementary profiles (**Table S2**). This effect waned by 14 days post-allergen challenge (data not shown).

Altogether, these data indicate that allergic reactions are sufficient to induce a blood TG increase and a unique lipid signature that is mostly triggered by the IgG-mediated alternative pathway of anaphylaxis.

### Allergic patients show increased blood triglyceride levels after an allergic reaction

Considering the murine data, we decided to explore whether allergic reactions were able to induce blood TG changes in humans. Many common allergens are environmental and ubiquitous^34,35^, which makes it difficult to control allergen exposure. In addition, subclinical exposures to the allergen might have an immunological impact too complex to be evaluated and controlled. Notwithstanding, we collected blood samples from 59 patients during the onset of an allergic reaction, and another one ≥14 days later (**Fig. 7a**) to measure if allergic inflammation had induced changes in blood TG levels. Most of the patients presented hypersensitivity reactions of grade I and II severity^36^, triggered by drugs or foods (**Table 1**). Given that some anti-inflammatory drugs may alter the blood TG levels^37^, the initial sample was taken prior treatment of the reaction, and the second sample was taken at least 14 days later to allow for drug clearance. We also evaluated in different preclinical models of anaphylaxis that blood TG levels did not change shortly after the allergic reaction (data not shown) and remain unchanged for up to 24 hours after it (**Fig. 5e**).

**Fig. 7.**
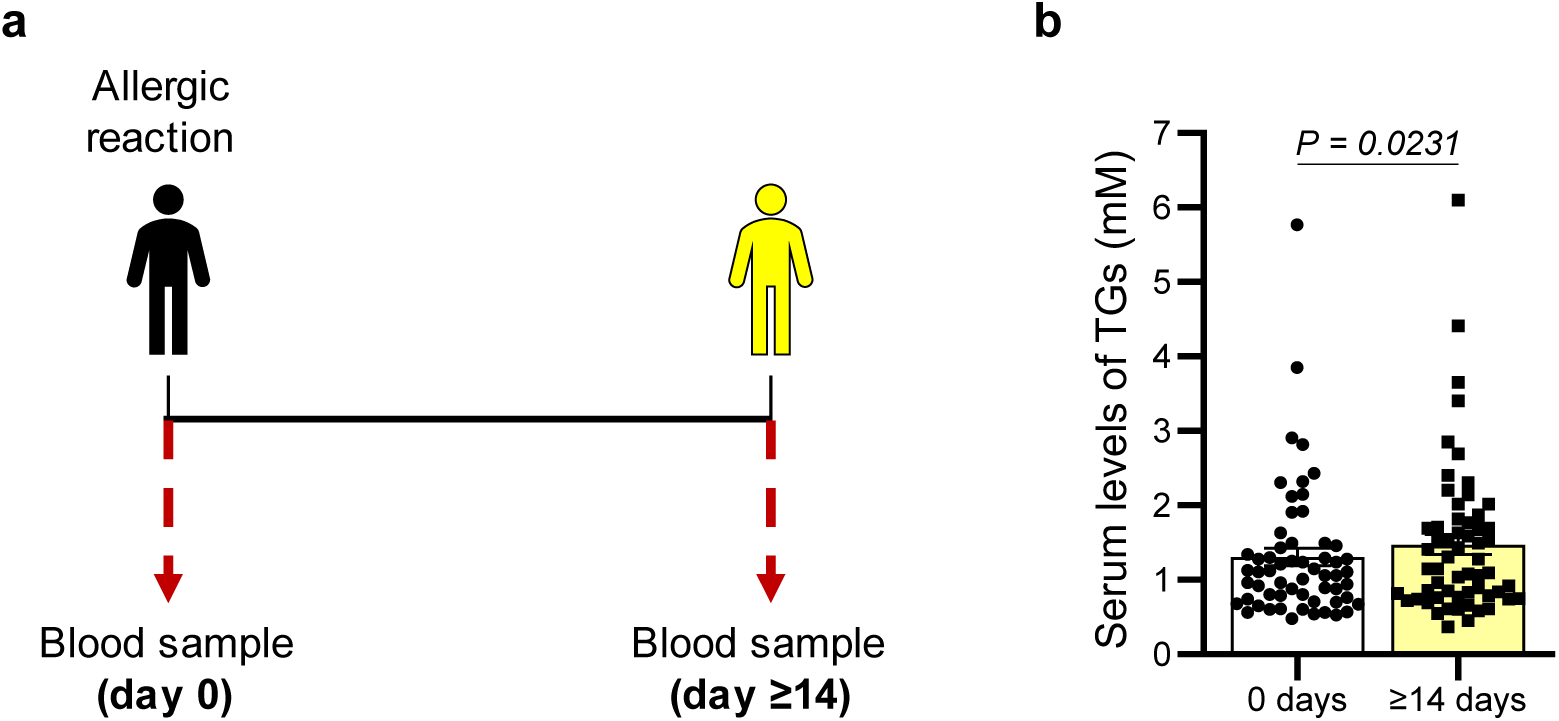
Allergic patients show increased blood triglyceride (TG) levels days after undergoing an allergic reaction. **a** Protocol followed for serum collection from 59 allergic patients. **b** Serum TG levels measured at the onset of the allergic reaction and ≥14 days later. Data presented as mean ± s.e.m. Each dot represents an individual patient. Statistically significant differences were calculated with the Wilcoxon rank-sum test.

**Table 1.**
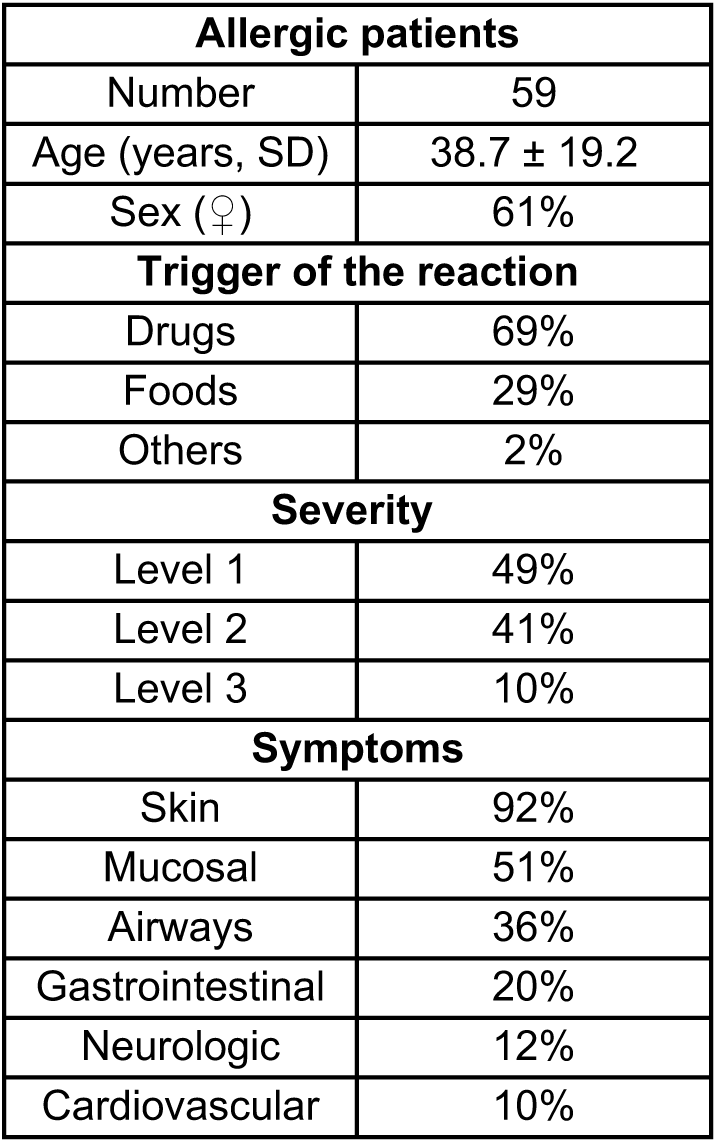
Allergic reactions.

Of note, the serum TG levels of allergic patients increased significantly following an allergic reaction (**Fig. 7b**). These data support our previous findings in mice and confirm that allergic reactions alter blood TG levels in humans as well.

## DISCUSSION

Recent studies in humans indicate that allergic disease may cause metabolic changes. For example, it has been reported that children with food allergy have lower serum levels of sphingolipids and ceramides^17^, and severe respiratory allergy has been associated with higher plasma levels of fatty acids^19^. Along the same line, the metabolomic profiling of serum from food allergic children, at the time of diagnosis and allergy resolution, revealed that the lipid profile is closely related to allergy outgrowth^18^. Also, it has been reported that patients with allergic rhinitis, asthma and atopic dermatitis have increased risk of dyslipidemia^38–40^. While these studies evince the existence of an interplay between allergic disease and lipid metabolism, the mechanisms involved are poorly understood, particularly the contribution of the different phases of allergic inflammation to dyslipidemia.

Using a murine model of allergy and atherosclerosis, we observed that allergic mice exhibited dyslipidemia, which was characterized by increased levels of serum TGs. Intriguingly, this serum TG increase detected in allergic animals was not inherent to allergic disease, but specific to allergic inflammation. Hence, quiescent allergy or atopy in and of itself did not alter serum TG levels as previously reported^41^, but an allergic reaction was required for this to happen. Also, these changes in serum TGs appear to occur regardless of the diet because the blood was collected under fasting conditions in atherosclerosis-prone mice fed HC and RE diets. The gene expression analyses in adipose tissue and liver following allergen challenge revealed changes in lipid metabolism, particularly in *Lipin1* and *Srebf1*, which could be linked to the atherogenic lipid profile induced by allergic inflammation^42,43^.

Several studies in humans have established a long-lasting association between elevated serum TG levels and cardiovascular disease^44–47^, although it is still unclear whether TGs promote disease via direct or indirect mechanisms. A recent study combined large-scale human genetic analysis with experimental evidence. It showed that TG-lowering alleles involved in hepatic production of TG-rich lipoproteins (*TM6SF2* and *PNPLA3*) and peripheral lipolysis (*LPL* and *ANGPTL4*) were associated with lower risk of cardiovascular disease^48^. Also, elevated serum TG levels have been recently identified as a risk factor of subclinical atherosclerosis in the PESA study, even when the LDL/HDL ratio was normal^25^.

A number of studies both in rodents^49,50^ and humans^51,52^ have explored metabolic changes following an allergic reaction, but focusing on the acute phase, to unveil biomarkers or mediators of anaphylaxis. In other words, these studies analyzed metabolic changes in circulation minutes or hours after the allergic reaction was triggered. Our comprehensive, untargeted lipidomic analysis in plasma was performed days (1, 3, and 7) after allergen challenge, well beyond the acute phase reaction was over. Strikingly, these analyses revealed a unique lipid fingerprint in allergic mice following an allergic reaction, particularly at 7 days post allergen exposure, as compared to unchallenged allergic mice. Their profile was characterized by an increase in 25 specific TGs, some of which have been positively associated with risk of cardiovascular disease. For example, the lipidomic analysis of 685 plasma samples of the prospective population-based Bruneck study found that TG 56:6, TG 52:2, TG 50:2, and particularly TG 54:2, were significantly associated with cardiovascular disease risk^26^. Furthermore, using 5,662 serum lipid profiles from PROMIS (a case-control study of first-ever acute myocardial infarction), Harshfield *et al*. identified lipids associated with the genetic polymorphism rs662799 in the *APOA5−APOC3* locus, which is linked to coronary artery disease. Of these, TG 53:3 and TG 56:4 were positively associated with risk factors of cardiovascular disease including hypertension and diabetes, respectively^28^. Also, TG 38:0 has been identified as a serum biomarker of Wilsońs disease in a cohort of 34 patients that included 34 relatives of these patients and 64 healthy individuals as controls. Wilsońs disease is a genetic pathology characterized by copper accumulation and dysregulated lipid metabolism^53^. Although the relationship between Wilsońs disease and atherosclerosis has not been defined, it has been reported that Wilsońs disease patients are at a higher risk of suffering cardiovascular events^54^. On the other hand, Fernandez *et al*. performed a plasma lipidome using 427 samples from the prospective population-based Malmö Diet and Cancer (MDC) study. They determined that TG 48:2 was negatively associated with cardiovascular disease^27^. While further validation of these TG species is required in well-characterized cohorts of allergic patients, it is noteworthy that a substantial number of the TG species identified in the murine model have been associated with risk of cardiovascular disease in humans.

To better understand the interplay between allergic disease and lipid metabolism, we took different approaches to ascertain the contribution of cellular and humoral inflammation to serum TG changes. Given that the TG peak in serum was observed between 3- and 7-days post-allergen exposure, and the lipid profile was unique 7 days after the allergic reaction, we considered that these changes were associated with late phase inflammation, which peaks 3 days after challenge in this model^55^. Although different players such as type 2 innate lymphoid cells can contribute to late phase inflammation^56^, CD4 Th2 cells orchestrate it^16^. Surprisingly, the impairment of CD4-T-cell-driven late-phase inflammation did not affect the increase in serum TGs, thus pointing towards the acute phase response. Indeed, using models of passive sensitization, where there is no type 2 adaptive immune response established, but exclusively the humoral components transferred, we demonstrated that the acute phase alone was sufficient to recapitulate the increase in serum TGs. Furthermore, we delved into the role of the classical (IgE-mediated) and alternative (IgG-mediated) pathways of anaphylaxis to dyslipidemia. Strikingly, the classical pathway of anaphylaxis appeared to be dispensable for the changes in serum TGs, as these were only observed when the alternative pathway or both were active. Also, the alternative pathway was sufficient to induce a unique lipid signature, clearly different from the one produced by the classical pathway, which was similar to that of the control group. The identification of lipid species that are specifically increased days after the onset of an IgG-mediated reaction may be helpful to appraise its actual contribution to human anaphylaxis, which still lacks specific biomarkers^14^.

From a broader perspective, these data display a novel role for humoral responses, in particular those of the IgG isotype insofar as they can regulate the lipid metabolism in response to allergens. However, it is plausible to consider that this mechanism may operate in other diseases, not necessarily type 2 driven, where strong IgG responses are chronically induced^57,58^. For example, autoimmune diseases, such systemic lupus erythematosus and rheumatoid arthritis, are associated with dyslipidemia and high serum TG levels, even in patients naive to corticosteroids^59–61^.

Assessing the precise contribution of acute *vs* late allergic inflammation to serum TG levels in humans is almost an unsurmountable challenge. Even so, we attained to show in a group of 59 patients that an allergic reaction induced an increase in serum TG levels days after the reaction. A recent transcriptomic study in blood of 3,229 individuals from the BIOS (Biobank-based Integrative Omics Study) consortium found an association between TG levels and canonical genes of type 2 immunity^21^. Although information pertaining to allergic disease of these patients was not available, they found that TG levels were associated with a downregulation of the hallmark Th2 cytokine, *IL4,* two IgE-receptor genes (*FCER1A* and *MS4A2*) as well as genes related to allergic mediators or its metabolism (*HDC, HRH4, CPA3, HPGDS, CYP11A1, PTGER3*)^21^. In this context, one may speculate that the increase of TGs observed following an allergic reaction is the output of a regulatory mechanism that prevents immunopathology by reducing the expression of key molecules of the allergic machinery. Yet, the chronic use of this putative mechanism of control may render a detrimental lipid profile in circulation.

### EXPERIMENTAL PROCEDURES

### Mice

Age-, sex-, and strain-matched controls were used in all the experiments. B6;129S7-Ldlrtm1Her/J mice were crossed with B6.SJL-Ptprc^a^Pepc^b^/BoyCrl mice and selected by genotype (LDLr KO) at CNIĆs Animal facility. C57Bl/6 mice were purchased from Charles River. Mice were maintained in biohazard specific pathogen-free conditions on a 12-hour light-dark cycle with a chow diet (RE; LASQCdiet Rod18-A, Altromin) or a HC diet (SSNIFF, S9167-E010) and water *ad libitum*.

All procedures were approved by the Environmental Council of the Community of Madrid (Madrid, Spain) with PROEX references 45.2/20 and 9.7/23.

### Allergy model

Male mice were sensitized with 4 mg of PN butter (∼1 mg of protein; Capitán Mani) and 5 μg of cholera toxin (C8052, Sigma) in 0.5 mL of PBS administered intragastrically, weekly for 4 weeks. Following sensitization, mice were challenged intraperitoneally with 2.5 mg of crude peanut extract (CPE; XPF171D3A25; Stallergens Greer), or 88 μg of CPE per gram of body weight, and core temperature was assessed rectally with a dual thermometer (620-2049, VWR). Serum or plasma samples were collected at different times of allergic pathology under 12-16 hours of fasting and stored at -80°C for further analyses. In some instances, peritoneal lavage was performed with ice-cold 10 mM EDTA in PBS and spleens were processed to assess eosinophilia and CD4 T-cell depletion by flow cytometry^23^, or for cell culture (see below).

### Spleen cell cultures and assessment of cytokine production

Spleens were processed and cultured as reported.^22^ After 5 days of culture, supernatants were stored at -80°C until further analysis. Cytokines in cell-free supernatants were quantified with LEGENDplex™ MU Th1/Th2 Panel (8-plex) using a BD FACSymphony A5 cell analyzer. The cytokine values obtained from supernatants of cells incubated with media were subtracted from those of allergen-stimulated cells.

### Antigen-specific Ig ELISA

ELISAs were performed in a 96-well flat bottom polystyrene plate (3590, Corning). Plates were washed with 0.05% Tween-20 in PBS (washing buffer), and 10% FBS/1% BSA in washing buffer was used as blocking solution. Plates were developed with 50 μL TMB (421101, BioLegend) and stopped using 50 μL 1.5 N H_2_SO_4_. Plates were read at 450 nm using a Thermo Scientific Multiskan FC. No-sample controls were included in all assays. The average optical density of the no-sample control wells was subtracted from all samples prior to plotting.

To determine the <u>serum levels of PN-specific IgE</u>, plates were coated with 50 μL of 2 μg/mL anti-mouse IgE (clone R35-72M; 553413, BD) in PBS, sealed with aluminum foil, and incubated overnight at 4°C. The next day, plates were washed and blocked for 2 hours at room temperature (RT). Plates were washed and incubated with 50 μL of serum diluted to the indicated concentrations in blocking solution overnight at 4°C. A standard curve was included of purified mouse IgE k Isotype control (557079, BD Biosciences) from 120 ng/mL to 0.9 ng/mL diluted in PBS. The next day, plates were washed and incubated with 50 μL of biotinylated-CPE at 150 ng/mL while standards were incubated with 50 μL of blocking solution, for 90 min at RT. Plates were washed and the samples were incubated with 50 μL of streptavidin-HRP diluted in blocking solution and standards with 50 μL of 0.175 μg/mL rat anti-mouse IgE-HRP (1130-05, Southern Biotech) diluted in blocking solution for 1 hour at RT. Lastly, plates were washed, developed, stopped, and read.

To determine the <u>serum levels of PN-specific IgG1</u>, plates were coated with 10 μg/mL CPE in 50 μL of carbonate bicarbonate buffer, sealed with aluminum foil, and incubated overnight at 4°C. The next day, plates were washed and blocked for 2 hours at RT. Plates were washed and incubated with 50 μL of serum diluted to the indicated concentrations in blocking solution overnight at 4°C. The next day, plates were washed and incubated with 50 μL of 0.5 μg/mL goat anti-mouse IgG1-HRP (A10551, Thermo Fisher Scientific) in blocking solution for 2 hours at RT. Ultimately, plates were washed, developed, stopped, and read.

### Antibody treatments and passive sensitization

CD4 T-cell depletion was accomplished via 2 intraperitoneal (i.p.) administrations of 200 µg of anti-CD4 (clone GK1.5; BE0003, BioXCell) one week and one day before challenge^22^. IgG from rat serum (I4131, Sigma-Aldrich) was used as isotype control. Cell depletion was confirmed by flow cytometry. For passive sensitization, a pool of sera collected from allergic or naive mice under fasting conditions was administered to naive LDLr KO mice via i.p. injection. For IgG-passive sensitization, part of the allergic sera pool was heated at 57°C for 4 hours to accomplish IgE denaturation^30,31^. Then, 700 μL of heat-treated sera was transferred to naive recipients the previous day to challenge. The irreversible denaturation and deactivation of IgE was confirmed by ELISA and mast cell activation test^32^. In the rest of the groups (naive, IgE, IgE+IgG), the naive or native allergic sera, respectively, were serially transferred as follows: 100 µL; 200 µL; and 400 µL with an interval of 48 hours between each administration. Then, the mice were challenged either the day after the last administration, to induce IgE- and IgG-mediated anaphylaxis, or 10 days later to allow for IgG clearance and IgE-mediated anaphylaxis^63^.

### Flow cytometry

Staining was conducted in FACS buffer (2.5 mM EDTA, 0.5% BSA, in PBS) in a 96 well V- or U-bottom plate (Corning, 353077) with combinations of the antibodies described in **Table S3** and the dilutions they were used at, based on 50 μL final staining volume. In all assays, cells were incubated with anti-FcγRII/III (clone 93; 101302, BioLegend) before incubation with fluorochrome-conjugated antibodies. Dead cells were excluded by fixable viability dye eFluor780 (65-0865-18, eBioscience) and gated on singlets. On average, a minimum of 50,000 live and singlet cells were analyzed. Fluorescence minus one (FMO) and unstained controls were used for gating. Data were acquired on a LSR II or Fortessa (BD) and analyzed using FlowJo (FlowJo LLC, Ashland, USA).

### RNA extraction and quantitative PCR

Total RNA was extracted from 30 mg liver tissue using NucleoSpin RNA/Protein mini kit (Macherey-Nagel) or from 100 mg abdominal white adipose tissue using the RNeasy Lipid Tissue mini kit (Qiagen) as recommended by the manufacturer. After homogenization of adipose tissue with QIAzol and previous to chloroform addition, samples were centrifuged 5 min at 12,000 g at 4°C to reduce lipid contamination (upper phase) of QIAzol fraction (lower phase). Residual DNA contamination was removed from 5 ug of RNA with the Turbo DNA-free Kit (Thermo Fisher). The retrotranscription was performed from 500 ng of DNAse-treated RNA using the High-Capacity cDNA RT kit (Applied Biosystems). Then, gene expression was measured by real-time quantitative PCR using SYBR green as a reporter (GoTaq(R) qPCR Master Mix, Promega) and mRNA specific primers (**Table S4**). Real-time qPCR analyses were performed with a QuantStudio 5 Real-Time PCR system, 384-well thermal cycler (Thermo Fisher). Expression of each gene of interest was normalized to at least 2 housekeeping genes: β-actin (*Actb*), glyceraldehyde-3-phosphate dehydrogenase (*Gapdh*) and hypoxanthine phosphoribosyltransferase 1 (*Hprt1*). Data are presented as relative fold differences calculated by using the 2-ΔΔCT method with average values of naive mice as a reference.

### Blood lipid profile

Murine serum samples were analyzed with the automated Dimension® RxL Max® clinical chemistry system from Siemens at the Comparative Medicine Unit of CNIC. The assay was performed using a Flex® reagent cartridge. The analysis of the lipid profile included reagents from Siemens (HDL, DF48B; Cholesterol DF27; DF131, LDL; DF55A, Lipase; DF69A, TGs), Spinreact (41035, Free Cholesterol), and Wako (436-91995, 436-91995; NEFA).

### Lipidomic analyses

Data acquisition and analysis was performed at CEMBIO and CBQF. Common reagents include reverse-osmosed ultrapure water from Millipore; LC-MS grade methanol (MeOH), acetonitrile (ACN), isopropanol (IPA) and sodium hydroxide from Fisher Scientific, Honeywell or VWR; HPLC grade methyl tert-butyl ether (MTBE), chloroform, ammonium fluoride (NH4F) (ACS reagent, ≥ 98%) and ammonium formate from Sigma-Aldrich or VWR. Analytical grade ammonia solution (28%, GPR RECTAPUR®) and acetic acid glacial (AnalaR® NORMAPUR®) were obtained from VWR Chemicals.

Lipidomics at <u>CEMBIO</u> (active sensitization model). Lipid extraction was performed as reported^64^. In brief, 50 µL of plasma from 63 mice were thawed on ice for approximately 1 hour and vortexed for 2 min. Subsequently, 800 µL of a previously prepared mixture consisting of MeOH/MTBE/CHCl_3_ (1.33:1:1, v/v/v) and 25 µL light Splash Lipidomics (Mass Spec Standard, 330707, Avanti Polar

Lipids, Inc.) used as the internal standard were added to each sample at RT for deproteinization and lipid extraction. Then, samples were vortex-mixed for 5 min at RT and centrifuged for 15 min at 15°C and 16,100 g. Finally, supernatants were transferred to chromatography vials with inserts and were directly injected into the LC-MS system. In addition, QC samples were set in parallel to the sample preparation. The QC was prepared by pooling equal volumes (30 µL) of each supernatant sample from the study. The QC was split into 5 vials, and these were analyzed throughout the analysis process.

Samples were analyzed using reversed-phased ultra-high liquid chromatography (RP-UHPLC) carried out on an Agilent 1290 Infinity II UHPLC system coupled to an Agilent 6545 quadrupole time-of-flight (QTOF) mass spectrometer equipped with Dual Agilent Jet Stream Electrospray source (Dual AJS ESI). The Agilent 1290 Infinity II Multisampler system, equipped with a multi-wash option, was used to uptake 1 and 2 µL of extracted samples in positive and negative ionization modes, respectively. The multisampler was maintained at 15°C to preserve lipids in a stable environment and avoid precipitation. An Agilent InfinityLab Poroshell 120 ECsingle bondC18 (3.0 × 100 mm, 2.7 µm) (Agilent Technologies) column and a compatible guard column (Agilent InfinityLab Poroshell 120 ECsingle bondC18, 3.0 × 5 mm, 2.7 µm) were used and maintained at 50°C. The chromatography gradient started at 70% of B at 0 – 1 minute, 86% at 3.5 – 10 min, 100% B at 11– 17 min. The starting conditions were recovered by minute 17, followed by a 2-min re-equilibration time; the total running time was 19 min. The mobile phases used for both positive and negative ionization modes consisted of (A) 10 mM ammonium acetate, 0.2 mM NH_4_F in H_2_O/MeOH (9:1, v/v) and (B) 10 mM ammonium acetate, 0.2 mM NH_4_F in ACN/MeOH/IPA (2:3:5, v/v/v). The flow rate was held constant, set at 0.6 mL/min. The multi-wash strategy consisted of a mixture of MeOH/IPA (50:50, v/v) with the wash time set at 15 s, and aqueous phase:organic phase (30:70, v/v) mixture to assist in the starting conditions.

The Agilent 6545 QTOF mass spectrometer equipped with a dual AJS ESI ion source was set with the following parameters: 150 V fragmentor, 65 V skimmer, 3500 V capillary voltage, 750 V octopole radio frequency voltage, 10 L/min nebulizer gas flow, 200°C gas temperature, 50 psi nebulizer gas pressure, 12 L/min sheath gas flow, and 300°C sheath gas temperature. Data were collected in centroid mode in positive and negative ESI modes in separate runs, operated in full scan mode from 50 to 1700 *m/z* with a scan rate of 3 spectra/s. A solution consisting of two reference mass compounds was used throughout the whole analysis: purine (C_5_H_4_N_4_) at *m/z* 121.0509 for the positive and *m/z* 119.0363 for the negative ionization modes; and HP-0921 (C_18_H_18_O_6_N_3_P_3_F_24_) at *m/z* 922.0098 for the positive and *m/z* 980.0163 (HP-0921+acetate) for the negative ionization modes. These masses were continuously infused into the system through an Agilent 1260 Iso Pump at a 1 mL/min (split ratio 1:100) to provide a constant mass correction. Ten iterative-MS/MS runs were performed for both ion modes at the end of the analytical run. They were operated with a MS and MS/MS scan rates of 3 spectra/s, 40 – 1700 *m/z* mass window, a narrow (∼ 1.3 amu) MS/MS isolation width, 3 precursors per cycle, 5,000 counts, and 0.001% of MS/MS threshold. Five iterative-MS/MS runs were set with a collision energy of 20 eV, and the subsequent 5 runs were performed at 40 eV. Reference masses and contaminants detected in blank samples were excluded from the analysis to avoid inclusion in the iterative-MS/MS.

The data collected after the LC-MS analyses in both positive and negative ion modes were reprocessed with the Agilent MassHunter Profinder B.10.0.2 software. The datasets were extracted using the Batch Recursive Feature Extraction (RFE) workflow integrated into the software. This workflow comprises two steps: the Batch Molecular Feature Extraction (MFE) and the Batch Find by Ion Feature extraction (FbI). The MFE algorithm consists in removing unwanted information, including the background noise, and then creating a list of possible components that represent the full range of TOF mass spectral data features, which are the sum of coeluting ions that are related by charge-state envelope, isotope pattern, and/or the presence of different adducts and dimers. Additionally, the MFE is intended to detect coeluting adducts of the same feature, selecting the following adducts: [M+H]^+^, [M+Na]^+^, [M+K]^+^, [M+NH_4_]^+^ and [M+C_2_H_6_N_2_+H]^+^ in LC-MS positive ionization; [M-H]^−^, [M+CH_3_COOH-H]^−^, and [M+Cl]^−^ in LC-MS negative ion mode. The neutral loss (NL) of water is also considered for both ion modes. The algorithm then aligns the molecular features across the study samples using the mass and retention time to build a single spectrum for each compound group. The next step involves FbI, using the median values derived from the MFE process to perform a targeted extraction to improve the reliability of finding and reporting features from complex datasets used for differential analysis.

After data pre-processing, quality assurance (QA) consisted of raw data filtration by keeping all features that were present after blank subtraction, were detected in >50% of QCs and >75% in at least one sample group and had Relative Standard Deviation (RSD) <30% in the QCs. The rest of the signals were excluded from the analyses. Finally, 1,106 and 268 chemical entities were obtained that passed LC-MS quality control in positive and negative ionization, respectively. Missing values were replaced using the k-nearest neighbors (kNN) algorithm^65^ using an in-house script developed in Matlab® (v.R2018b, MathWorks®). The quality of the analyses was tested using PCA models^66^. The lipid annotation process was carried out in three steps: the first one was an initial tentative identification of lipid features, based on the MS1 data, using our online tool CEU Mass Mediator (CMM) (http://ceumass.eps.uspceu.es/mediator/)^67^. This tool for mass-based compound annotation comprises the information available in different databases (KEGG, HMDB, LIPID MAPS, Metlin, MINE and an in-house lipid library). This stage started with the tentative assignment based on (i) accurate mass (maximum mass error tolerance 20 ppm); (ii) retention time; (iii) isotopic pattern distribution; and (iv) adduct formation pattern.

Secondly, to increase the level of confidence annotation, the raw LC-MS/MS data obtained was imported to the Lipid Annotator software (Agilent Technologies Inc.) and the MS-DIAL software^68^, to build a fragmentation-based (MS/MS) library comprising the *m/z* of all the precursors identified as lipids by the software, together with their corresponding RT. The Lipid Annotator method^69^ was set as follows: ion species [M+H]^+^, [M+Na]^+^, and [M+NH_4_]^+^ for positive; and [M-H]^−^, and [M+CH_3_COOH-H]^−^ for negative ionization mode. Then, for both ion modes, the Q-Score was set at ≥ 50; all the lipid classes were selected, the mass deviation was established as ≤ 20 ppm, fragment score threshold was fixed as ≥ 30, and the total score was set at ≥ 60. For the case of MS-DIAL software, analytical parameters were also set as described: ion species [M+H]^+^, [M-H_2_O+H]^+^, [M+Na]^+^, [M+K]^+^, [M+NH_4_]^+^ and [M+C_2_H_6_N_2_+H]^+^ for positive; and [M-H]^−^, [M-H_2_O-H]^−^, [M+Cl]^−^, [M+HCOOH-H]^−^ and [M+CH_3_COOH-H]^−^ for negative mode^64^. Adduct formation with formic acid were observed experimentally in this study even though the mobile phases used for LC are lacking this compound. Formic acid presence was considered due to trace amount contamination levels of formate in acetate salts of LC-MS grade, or because MeOH can be oxidized to formic acid during electrospray ionization. The search was fixed to be performed across a mass range from 50 to 1500 Da (for both MS1 and MS/MS levels), the mass deviation accepted was ≤ 0.01 Da and ≤ 0.025 for MS1 and MS/MS levels respectively, and the identification score cut off was also set at ≥ 60.

Finally, a manual MS/MS spectral interpretation was carried out using the software Agilent MassHunter Qualitative (version 10.0), matching the retention time and MS/MS fragmentation to the available spectral data included in MetFrag and Lipid Maps^70^. Different forms of cations and anions were targeted for the same molecule to ensure its annotations. The interpretation of each obtained spectrum led to the same ID in terms of belonging to the same lipid category and in terms of fatty acids composition, which together give a robust and complete identification. The nomenclature used in this article for the lipid species reported follows the latest update of the shorthand annotation.^71^ Lipid metabolic networks according to their chain length, saturation, and level of significance were performed by Lipid Network Explorer (LINEX) version 2, a web-tool to analyze lipid metabolic networksR^72^.

<u>Lipidomics at CBQF</u> (passive sensitization model). Lipid extraction from 20 plasma samples was performed as reported^73^ with some modifications. Briefly, tubes were added with 1.5 mL of MeOH and 5 mL of MTBE. Samples were homogenized for 1 min followed by 15 min of sonication in ultrasound. This procedure was repeated once again and then the tubes let under rotation (70 rpm) overnight (18 hours). Afterwards, tubes were added with 1.25 mL of HPLC-water, homogenized for 1 minute and centrifuged (1250 g, 10 min, RT). About 3 mL of the upper layer (MTBE containing the extracted lipids) were recovered to a new tube previously weighted. Then, 3 mL of MTBE were added to extraction tubes and these homogenized and centrifuged at the same conditions mentioned above. Upper layer was recovered, and the MTBE extract let to evaporate completely, under nitrogen, for further determination of lipid extract mass.

Plasma lipid extracts were pooled according to their group, dissolved in IPA/ACN (9:1, v/v) at 0.2 mg/mL and analyzed in triplicate on an UHPLC instrument (Elute; Bruker), equipped with an Acquity UPLC BEH C18 (17 µm) pre-column (Waters), an Intensity Solo 2 C18 (100 x 2.1 mm) column (Bruker), and coupled with an UHR–QTOF detector (Impact II; Bruker). The injection method was based on conditions previously reported with some modifications^74^. Thus, mobile phases consisted of ACN/H_2_O (6:4, v/v) (Phase A) and IPA/ACN (9:1, v/v) (Phase B), each one added with 0.1% (v/v) formic acid and 10 mM ammonium formate. Phase B gradient flow was set as follows: 0.0 min: 40%, 2.0 min: 43%, 2.1 min: 50%, 12.0 min: 54%, 12.1 min: 70%, 18.0 min: 99%, 20.0 min: 99%, 20.1 min: 40%, and 22 min: 40%. Flow rate was set at 0.4 mL/min and column temperature at 55°C. The injection volume was 3 µL in positive ionization mode and 5 µL in negative ionization mode. For MS analysis, the following parameters were applied: end plate offset voltage 500 V, capillary voltage 4500 V (positive ionization) or 3000 V (negative ionization), nebulizing gas pressure 35 psi, drying gas flow 8 L/min, drying gas temperature 325°C, quadrupole ion energy 3 eV (positive ionization) or 5 eV (negative ionization), collision energy 10 eV (positive ionization) or 5 eV (negative ionization). Acquisition was performed in an auto MS/MS scan mode over a mass range of m/z 50-1500. For both ionization modes, an external mass calibration was performed with a solution of IPA/H_2_O (1:1, v/v) added with 0.2% (v/v) formic acid and 0.6% (v/v) NaOH 1M, continuously injected at 180 µL/hour. The files resulted from LC-ESI-QTOF analysis were converted into abf format through Reifycs Analysis Base File Converter (Reifycs Inc.). These converted files were used for lipid composition analysis in MS-DIAL 5.1.230517 (Riken). The alignment results were exported and identified compounds were grouped by lipid subclasses, according to LIPID MAPS database classification. Peak intensities of compounds belonging to same lipid class were totalized and used as data for heatmap analysis in ClustVis, a web tool available for visualizing clustering of multivariate data^75^. The MS-DIAL software detected 2,718 different ions among the plasma samples, being annotated 1,236 compounds by the software. From these data were excluded, among the compounds with attributed equal identification, the ones that revealed considerably low signal, leading to a total of 1,198 compounds to assess further statistical analysis. For heatmap analysis, compounds were grouped into 36 subclasses.

### TG measurements in human sera

Human TGs were measured using the Cobas c 501 analyzer (Roche Diagnostic). For this purpose, serum samples (BD Vacutainer tubes) were collected from each patient under two conditions: one during the onset of a hypersensitivity reaction and the other at least 14 days after the reaction. Study population included 118 paired sera from 59 patients presenting hypersensitivity reactions and recruited from three Spanish hospitals (Hospital Universitario Fundación Jiménez Díaz, Hospital Central de la Cruz Roja, Hospital Infantil Universitario Niño Jesús). Acute samples were obtained in the emergency departments when allergen exposure was accidental and at the allergy units if the hypersensitive reaction was observed after a controlled challenge test. Of the 59 patients included, 9 had accidental reactions and 50 had clinically controlled reactions. Each patient’s sex, age, reaction trigger, clinical signs and symptoms were recorded. In addition, the severity of the reaction was determined according to the criteria established by Brown^36^. The study was approved by the Ethics Committee (CEIm FJD, PIC057-19), and authors adhered to the declaration of Helsinki. All patients were included after giving informed consent by the donors or their parents. Inclusion criteria included acceptance to participate in the study and an objective diagnosis of the reaction. Exclusion criteria were the presence of a blood-borne disease, any psychiatric illness or psychological pathology that prevented acceptance for the study or receiving any treatment prior to obtaining the acute phase sample, as these interfere with TG levels.

### Statistics

Independent experiments were performed with multiple biological replicates to ensure reproducibility and statistical power to detect meaningful differences. GraphPad Prism v9 was used for all statistical analyses. Quantitative variables are expressed as means ± standard error of the mean (s.e.m.). Normal distribution was determined by D’Agostino and Pearson test, Shapiro–Wilk test, Kolmogorov–Smirnov test, and Anderson-Darling test. A two-way analysis of variance (ANOVA) or mixed-effects test and one-way ANOVA were used to evaluate differences between more than two groups with a parametric distribution, and Bonferronís multiple comparison test was applied. Data with nonparametric distribution were evaluated with the Kruskal–Wallis test for more than two groups, and the Mann–Whitney or Wilcoxon signed-rank test for two independent groups. A p-value<0.05 was considered significant.

For lipidomic data, multivariate analysis was performed using SIMCA v.16.0 (Sartorius Stedim Data Analytics). The PCA models were used to evaluate data quality and find patterns in the experimental samples. The models were unit variance scaled and evaluated using R^2^ and Q^2^ parameters, which are the classification and prediction capacity, respectively. Complementary, univariate analysis was performed in MATLAB (v.R2018b, MathWorks®) or with the Lipidmaps’ statistical tool to obtain the significance of each feature in the study. The focus was set on the comparison between two groups at day 7. For this statistical analysis, Student’s t-test or Mann-Whitney U test was applied for each metabolite, depending on the normality of the data, which were evaluated by the Kolmogorow-Smirnow-Lilliefors test and the Levene test, respectively. The FDR was performed by using the Benjamini-Hochberg correction for these p-values, and statistical significance was set at 95% level (pFDR< 0.05 for the adjusted p-value)^76^. The MetaboAnalyst online tool (v. 5.0) was used to produce heat maps with hierarchical clustering. Euclidean distance measure and Ward’s clustering method were chosen as the clustering parameters. For lipid maps, LINEX – LipidNetworkExplorer version 2 web-tool was used (https://exbio.wzw.tum.de/linex)^72^. GraphPad Prism v9 was used to represent plots with metabolomics data.

## AUTHOR CONTRIBUTIONS

N.F.-G, R.C.-G. and R.J.-S. performed the *in vivo* experiments, qPCRs, multiplex immunoassays, and analyzed the data. L.M.-S. did ELISAs and analyzed the data. A.J.G.-C., A.G., D.O., L.M.-S, A.L.-F., L.L.-P., and R.N. performed lipidomics, and L.M.R.-A., A.V., D.B., and C.Barbas provided supervision and aided in lipidomics data processing. E.S.-M. and C.L.-S. conducted mast cell cultures and activation assays. M.R.-H. and I.R.-F. helped with experiments. E.N.-B., V.E., and C.Blanco provided samples from allergic patients and/or measured their lipid profile. Y.R.C., B.I., P.M. and F.S.-M. provided expertise in immunology and/or atherosclerosis throughout the project. R.J.-S conceptualized and oversaw the project, designed experiments, and wrote the manuscript. All authors discussed the results, assisted in the preparation of the manuscript, and approved its publication.

## ACKNOWLEDGEMENTS

R.J.-S.’s laboratory has received support from the Severo Ochoa Program (AEI/SEV-2017-0712; Spain), FSE/FEDER through the Instituto de Salud Carlos III (ISCIII; CP20/00043; PI22/00236; Spain), The Nutricia Research Foundation (NRF-2021-13; The Netherlands), New Frontiers in Research Fund (NFRFE-2019-00083; Canada) and SEAIC (BECA20A9; Spain). F.S.-M. reports grants from Ministerio de Ciencia e Innovación (MICIN; PID2020-120412RB-I00; Spain), and Comunidad de Madrid (INTEGRAMUNE, P2022/BMD7209; Spain). V.E. obtained support from the ISCIII (PI21/00158; Spain). N.F.-G. and I.R-F. received a Formación de Profesorado Universitario grant (FPU16/03953 and FPU20/05176, respectively) from Ministerio de Universidades (Spain). N.F.-G. and L.M.-S. are supported by the INVESTIGO Program of the Comunidad de Madrid and Ministerio de Trabajo y Economía Social, Servicio Público de Empleo (SEPE), respectively, which is funded by “Plan de Recuperación, Transformación y Resiliencia” and “NextGenerationEU” of the European Union (09-PIN1-00015.6/2022 and 2022-C23.I01.P03.S0020-0000031). R.C.-G. is supported by Ayudas para contratos Juan de la Cierva-formación 2021 (FJC2021-047282-I) from MICIN (Spain). E.N.B is supported by a Sara Borrell grant (CD23/00125) from ISCIII. P.M. is supported by MICIN-ISCIII-Fondo de Investigación Sanitaria (PI22/01759; PMPTA22/00090-BIOCARDIOTOX) and Comunidad de Madrid (P2022/BMD-7209-INTEGRAMUNE-CM; Spain). The CNIC is supported by the ISCIII, the MICIN, and the Pro CNIC Foundation. CNIC is a Severo Ochoa Center of Excellence (grant CEX2020-001041-S funded by MICIN/AEI/10.13039/501100011033). The R.J.-S.’s lab recognizes Ana Cayuela, Ibon Redondo-Angulo, Pedro L. Majano, Francisco Vega, María Ángeles Vallejo, Melissa Gordon, María Jesús Andrés-Manzano, Elena Prieto-García, and all the members of the R.J.-S. and F.S.-M. labs for technical help and/or scientific input.

