## Supplementary Figures for "Allergic inflammation triggers dyslipidemia via IgG signalling"

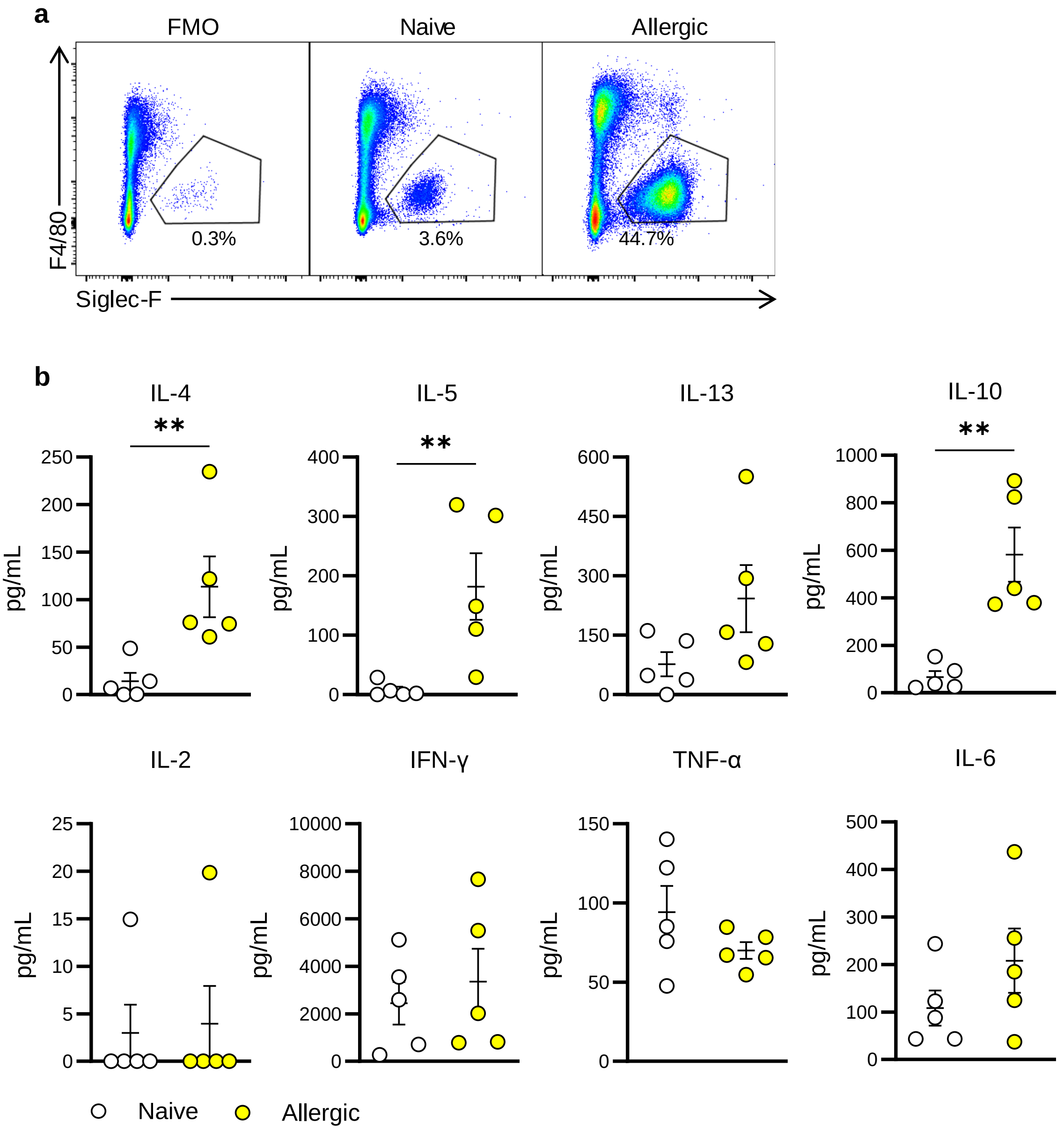


**Fig. S1 Assessment of eosinophilia and cytokine levels by flow cytometry. a** Cytometric identification of eosinophils in the peritoneal lavage of naive and allergic mice at 3 days post-allergen challenge, based on F4/80 and Siglec-F expression gated on live, singlets and CD45+ cells. **b** Quantification of cytokine levels by flow cytometry-based multiplex immunoassay in the supernatants of splenocytes collected from naïve and allergic mice 2 days post-allergen challenge and stimulated with allergen for 5 days. Data for **a** are representative plots of 4 independent experiments; data for **b** are presented as mean ± s.e.m.**,** and each dot represents an individual mouse. Statistically significant differences in **b** were calculated with the Mann-Whitney test (*p≤0.05; **p≤0.01). FMO, fluorescence minus one control.

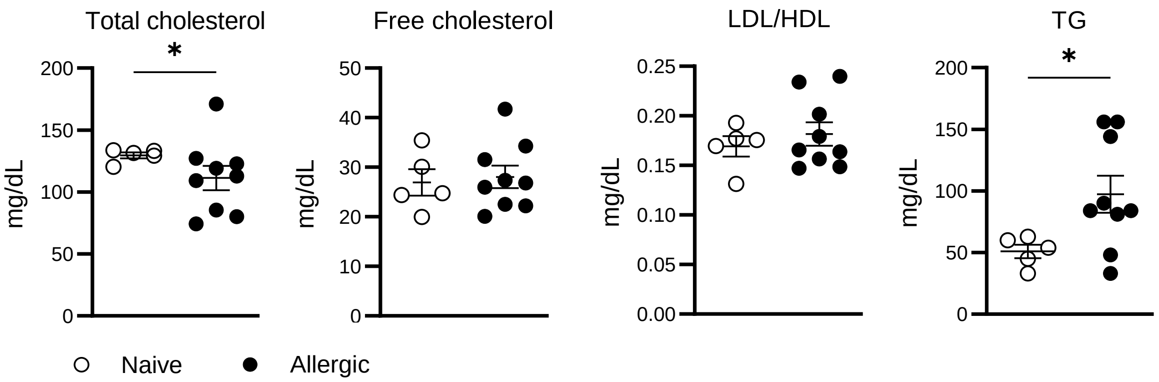


**Fig. S2 Serum triglyceride levels increase following allergic inflammation in wild type mice.** Serum levels of total cholesterol, free cholesterol, LDL/HDL ratio and triglycerides (TG) determined at 3 days post-allergen challenge in naive and allergic wild type mice fed a high-cholesterol diet. The data shown include 5 naïve and 9 allergic mice. Each dot represents an individual mouse. Data are presented as mean ± s.e.m. Statistically significant differences between groups for each time point were calculated with the Mann-Whitney test (*$p \leq 0.05$).

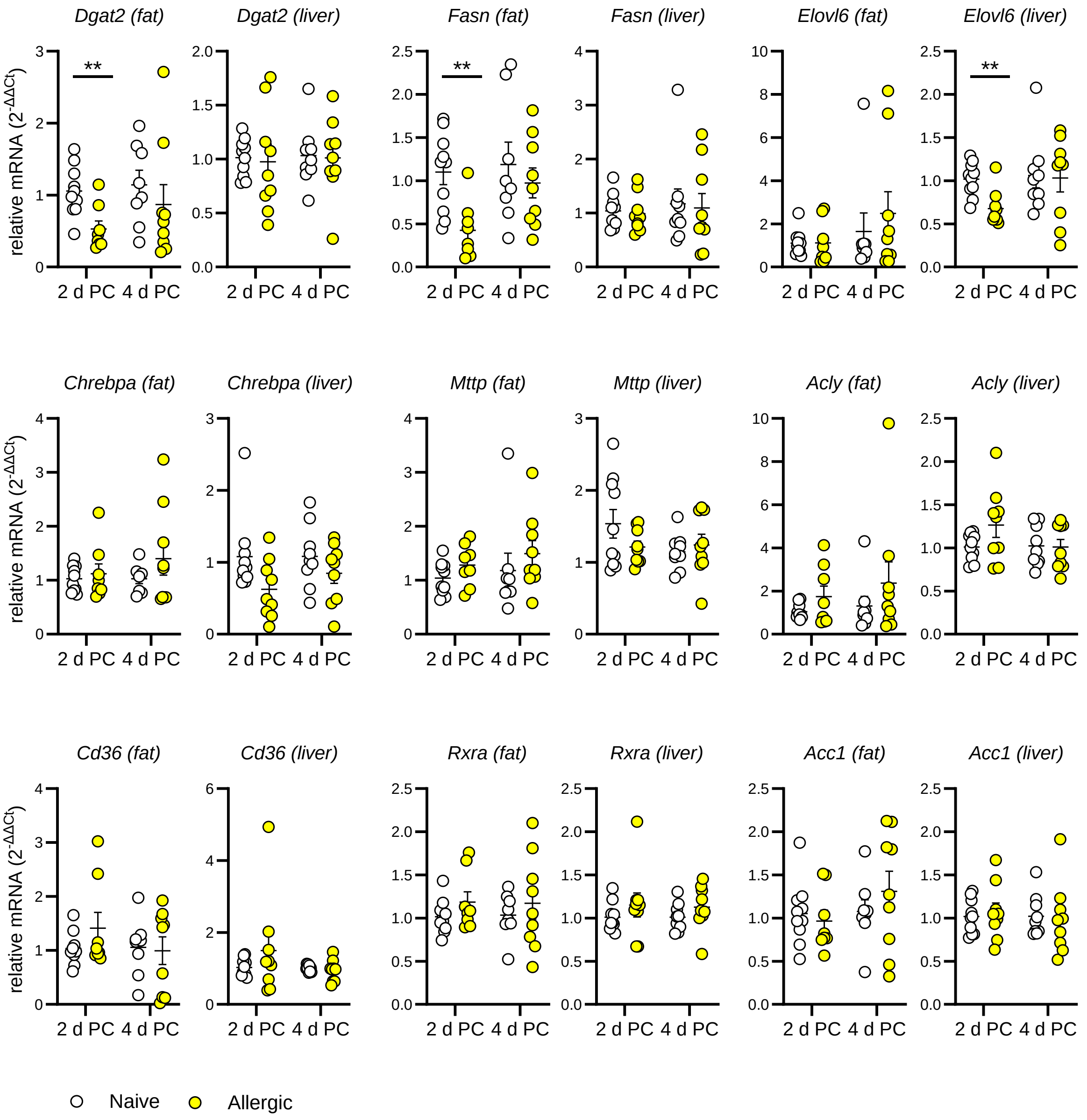


**Fig. S3 Expression of genes associated with lipid metabolism in naive and allergic mice following allergen challenge**. Relative mRNA expression of several genes associated with lipid metabolism in abdominal fat and liver of naive and allergic mice at 2- and 4-days post-allergen challenge (PC). The data shown include 10 naïve and 9 allergic mice; each dot represents an individual mouse. Data are presented as mean ± s.e.m. Statistically significant differences between groups for each time point were calculated with the Mann-Whitney test (*p≤0.05; **p≤0.01). Pool of 2 independent experiments.

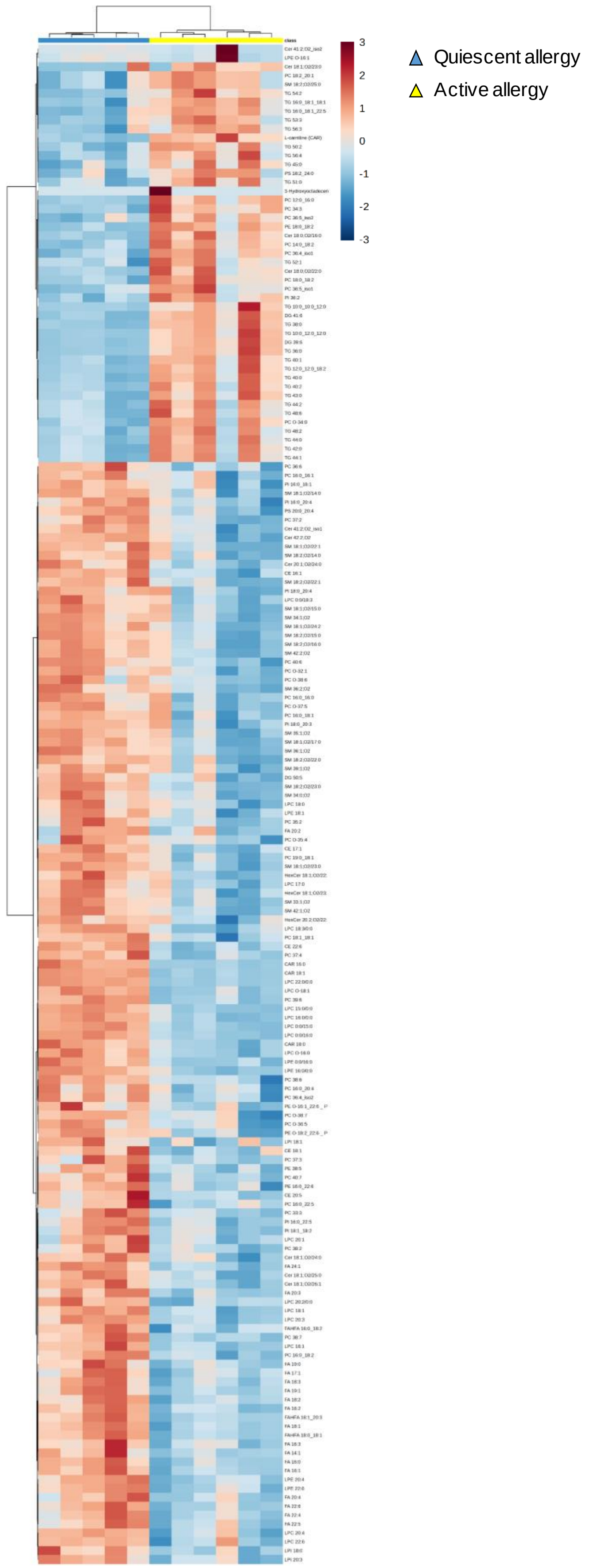


**Fig. S4 Allergic inflammation induces a unique blood lipid profile**. Heatmap of the 171 identified lipid species differentially expressed (p<0.05) between allergic mice with quiescent allergy (non-challenged, blue color, n=5) and active allergy (allergen-challenged, yellow color, n=6) 7 days after challenge. The figure shows the clustering results in the form of a dendrogram. Relative lipid abundance is represented along a gradient color from blue (diminished in active allergy) to red (increased in active allergy). DG; diglycerides; Cer, ceramides; CAR, carnitines; LPE, lysophosphatidylethanolamine; LPC, lysophosphatidylcholine; PC, phosphatidylcholine; CE, cholesteryl ester; PI, phosphoinositide; LPI, lysophosphoinositide; FA, fatty acid; HerCer, hexceramide; SM, sphingomyelin; PS, glycerophosphoserines; PE, glycerophosphoethanolamines; PE-O-, oxidate glycerophosphoethanolamines; PC-O-, oxidate phosphatidylcholine; FAHFA, fatty acid esters of hydroxy fatty acids; TG, triglycerides.

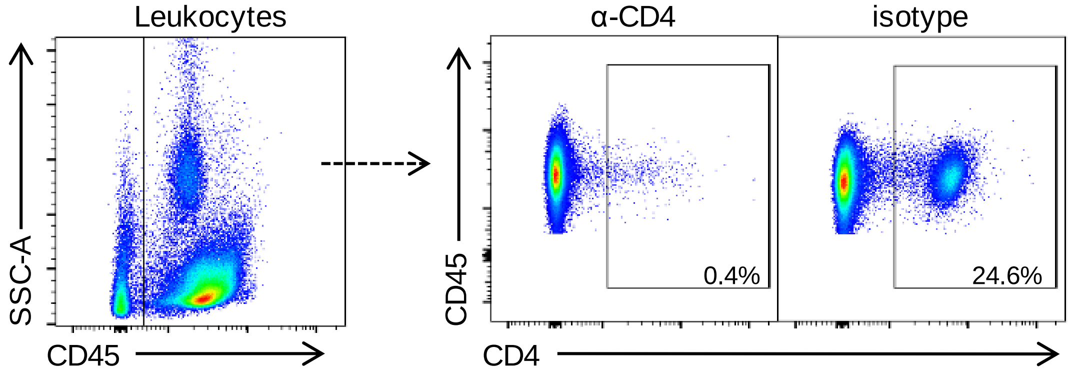


**Fig. S5 Assessment of CD4 T-cell depletion by flow cytometry.** Cytometric identification of CD4 T cells in the spleen of naive and allergic mice 7 days after α-CD4- (clone GK1.5) or isotype- treatment based on CD4 expression (clone RM4-5) gated on live, singlets and CD45+ cells. Representative plots of 3 independent experiments.

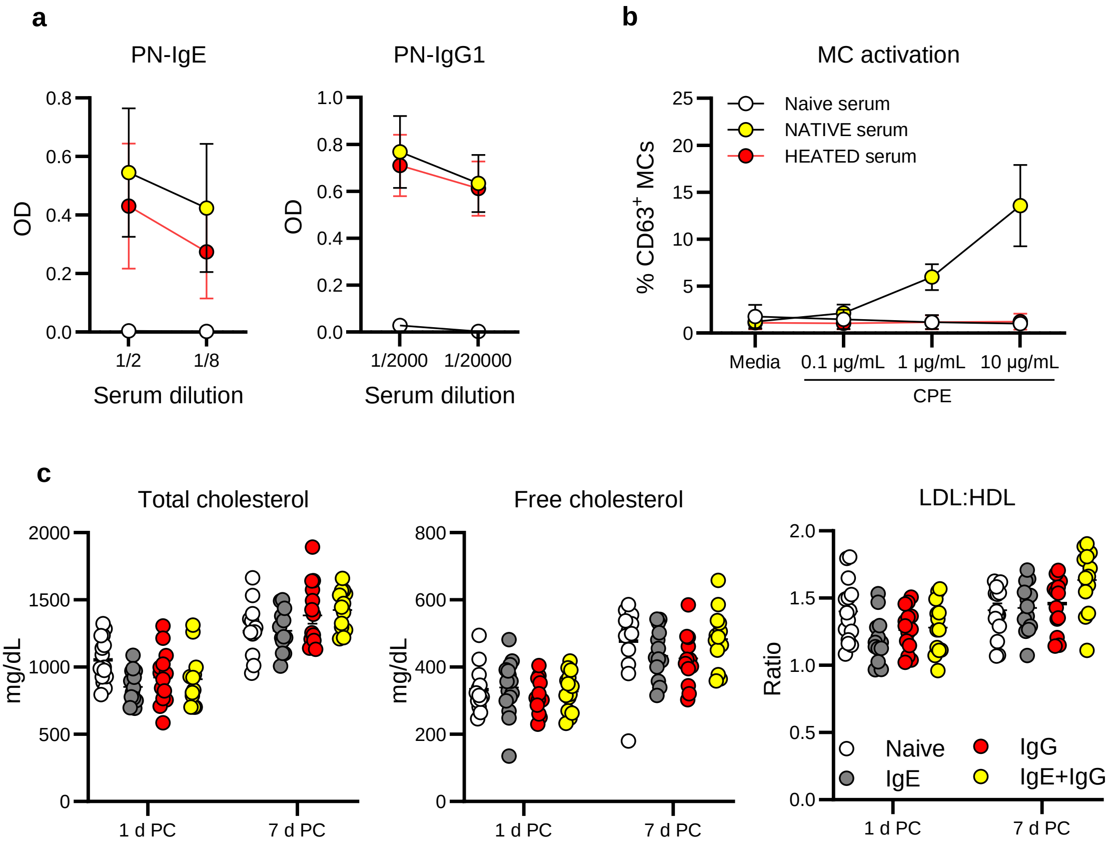


**Fig. S6. Characterization of allergic sera for passive sensitization and measurement of serum lipids in circulation following allergen challenge. a** Levels of peanut (PN)-specific IgE and IgG1 of naive and allergic sera (native or heated) used for passive sensitization. **b** Mast cell activation assays with cells sensitized with naive or allergic sera (native or heated) and challenged with different concentrations of crude peanut extract (CPE) (pooled data from 3 independent experiments). **c** Assessment of serum levels of total cholesterol, free cholesterol, and LDL/HDL ratio at 1- and 7- days post-allergen challenge (PC) in passively sensitized mice with naive (n=17) or allergic sera (native or heated) to induce anaphylaxis via the classical (IgE; n=19), the alternative pathway (IgG; n=19), or both (IgE+IgG; n=18).
