## Supplementary material for "Allergic inflammation triggers dyslipidemia via IgG signalling": 20240502 Supplementary Tables 1-4

| Table S1. Lipids differentially expressed between mice with quiescent allergy and active allergy at 7 days post-allergen challenge. | | | | | | | | |
|---|---|---|---|---|---|---|---|---|
| ESI | Compound Name | Formula | Mass | m/z | Error PPM | Rt (min) | CV_QC (%) | pBH/FDR* |
| (+) | CAR 18:1;O | C25H47NO5 | 441.3544 | 424.3509 | 20 | 1.98 | 3.59 | 0.0262 |
| (+) | CAR 16:0 | C23H45NO4 | 399.3353 | 400.3426 | 1 | 2.33 | 2.19 | 0.0035 |
| (+) | CAR 18:0 | C25H49NO4 | 427.3667 | 428.374 | 2 | 3.21 | 3.50 | 0.0035 |
| (+) | CAR 18:1 | C25H47NO4 | 425.3508 | 426.3581 | 1 | 2.52 | 2.23 | 0.0000 |
| (+) | CE 16:1 | C43H74O2 | 622.568 | 640.6018 | 1 | 14.72 | 1.94 | 0.0262 |
| (+) | CE 17:1 | C44H76O2 | 636.5821 | 654.6159 | 4 | 15.11 | 9.27 | 0.0075 |
| (+) | CE 18:1 | C45H78O2 | 650.6004 | 668.6342 | 0 | 15.48 | 2.03 | 0.0262 |
| (+) | CE 20:5 | C47H74O2 | 670.5688 | 688.6026 | 0 | 13.94 | 8.59 | 0.0262 |
| (+) | CE 22:6 | C49H76O2 | 696.5853 | 714.6191 | 1 | 14.09 | 4.12 | 0.0023 |
| (-) | Cer 18:0;O2/16:0 | C34H69NO3 | 539.5141 | 598.528 | 25 | 9.94 | 3.73 | 0.0106 |
| (-) | Cer 18:0;O2/22:0 | C40H81NO3 | 623.6204 | 682.6343 | 2 | 12.33 | 2.85 | 0.0259 |
| (-) | Cer 18:1;O2/23:0 | C41H81NO3 | 635.6215 | 694.6354 | 0 | 12.36 | 6.20 | 0.0169 |
| (+) | Cer 18:1;O2/24:0 | C42H83NO3 | 649.6371 | 650.6444 | 0 | 12.52 | 8.48 | 0.0233 |
| (-) | Cer 18:1;O2/25:0 | C43H85NO3 | 663.6531 | 722.667 | 0 | 12.61 | 4.09 | 0.0047 |
| (-) | Cer 18:1;O2/26:1 | C44H85NO3 | 675.6405 | 734.6544 | 18 | 12.51 | 4.32 | 0.0030 |
| (-) | Cer 20:1;O2/24:0 | C44H87NO3 | 677.6667 | 736.6806 | 3 | 12.83 | 10.05 | 0.0075 |
| (-) | Cer 41:2;O2_iso1 | C41H79NO3 | 633.606 | 692.6199 | 0 | 12.21 | 3.39 | 0.0108 |
| (-) | Cer 41:2;O2_iso2 | C41H79NO3 | 633.6033 | 692.6172 | 4 | 12.08 | 3.03 | 0.0246 |
| (-) | Cer 42:2;O2 | C42H81NO3 | 647.6225 | 706.6364 | 1 | 12.212 | 3.32 | 0.0156 |
| (+) | DG 39:6 | C42H70O5 | 654.524 | 677.5132 | 3 | 12.26 | 3.80 | 0.0046 |
| (+) | DG 41:6 | C44H74O5 | 682.5538 | 705.543 | 0 | 12.54 | 3.87 | 0.0035 |
| (+) | DG 50:5 | C53H94O5 | 810.7134 | 833.7026 | 4 | 12.17 | 7.82 | 0.0298 |
| (-) | FA 14:1 | C14H26O2 | 226.1935 | 225.1862 | 1 | 2.13 | 2.13 | 0.0119 |
| (-) | FA 16:0 | C16H32O2 | 256.2408 | 255.2335 | 2 | 3.66 | 1.43 | 0.0030 |
| (-) | FA 16:1 | C16H30O2 | 254.2251 | 253.2178 | 2 | 2.98 | 3.71 | 0.0028 |
| (-) | FA 16:2 | C16H28O2 | 252.2087 | 251.2014 | 1 | 2.46 | 5.40 | 0.0035 |
| (-) | FA 16:3 | C16H26O2 | 250.1954 | 249.1881 | 8 | 2.05 | 3.59 | 0.0162 |
| (-) | FA 17:1 | C17H32O2 | 268.2404 | 267.2331 | 1 | 3.41 | 3.08 | 0.0169 |
| (-) | FA 18:1 | C18H34O2 | 282.2566 | 281.2493 | 2 | 3.82 | 1.34 | 0.0023 |
| (-) | FA 18:2 | C18H32O2 | 280.2408 | 279.2335 | 2 | 3.26 | 1.98 | 0.0076 |
| (-) | FA 18:3 | C18H30O2 | 278.2241 | 277.2168 | 2 | 3.662 | 3.43 | 0.0089 |
| (-) | FA 19:0 | C19H38O2 | 298.2871 | 297.2798 | 0 | 4.83 | 3.48 | 0.0136 |
| (-) | FA 19:1 | C19H36O2 | 296.272 | 295.2647 | 2 | 4.21 | 3.30 | 0.0169 |

| | | | | | | | | |
|---|---|---|---|---|---|---|---|---|
| (-) | FA 20:2 | C20H36O2 | 308.2715 | 307.2642 | 0 | 4.1 | 3.88 | 0.0397 |
| (-) | FA 20:3 | C20H34O2 | 306.2563 | 305.249 | 1 | 3.69 | 2.16 | 0.0035 |
| (-) | FA 20:4 | C20H32O2 | 304.2394 | 303.2321 | 3 | 3.16 | 2.28 | 0.0246 |
| (-) | FA 22:4 | C22H36O2 | 332.2727 | 331.2654 | 3 | 3.83 | 3.03 | 0.0125 |
| (-) | FA 22:5 | C22H34O2 | 330.256 | 329.2487 | 0 | 3.36 | 2.92 | 0.0279 |
| (-) | FA 22:6 | C22H32O2 | 328.2406 | 327.2333 | 1 | 2.98 | 1.67 | 0.0246 |
| (-) | FA 24:1 | C24H46O2 | 366.3501 | 365.3428 | 1 | 6.47 | 1.66 | 0.0012 |
| (-) | FAHFA 16:0_18:2 | C34H62O4 | 534.4621 | 533.4548 | 5 | 3.66 | 14.16 | 0.0162 |
| (-) | FAHFA 18:0_18:1 | C36H68O4 | 564.5122 | 563.5049 | 1 | 3.82 | 4.13 | 0.0015 |
| (-) | FAHFA 18:1_20:3 | C38H66O4 | 586.4937 | 585.4864 | 4 | 3.82 | 7.00 | 0.0028 |
| (+) | HexCer 18:1;O2/22:1 | C46H87NO8 | 781.6375 | 782.6448 | 7 | 11.72 | 15.25 | 0.0262 |
| (-) | HexCer 18:1;O2/23:0 | C47H91NO8 | 797.6722 | 856.6861 | 3 | 12.07 | 8.00 | 0.0275 |
| (-) | HexCer 20:2;O2/22:0 | C48H91NO8 | 809.6747 | 868.6886 | 0 | 11.94 | 3.33 | 0.0396 |
| (+) | L-carnitine (CAR) | C7H15NO3 | 161.1055 | 162.1128 | 2 | 0.74 | 1.92 | 0.0262 |
| (-) | LPC 0:0/15:0 | C23H48NO7P | 481.3169 | 540.3308 | 0 | 2.36 | 2.43 | 0.0001 |
| (-) | LPC 0:0/16:0 | C24H50NO7P | 495.3327 | 554.3466 | 1 | 2.36 | 1.96 | 0.0001 |
| (-) | LPC 0:0/18:3 | C26H48NO7P | 517.3207 | 576.3346 | 8 | 2.55 | 4.55 | 0.0136 |
| (-) | LPC 15:0/0:0 | C23H48NO7P | 481.3172 | 540.3311 | 1 | 2.55 | 2.70 | 0.0162 |
| (+) | LPC 16:0/0:0 | C24H50NO7P | 495.3325 | 496.3398 | 0 | 2.56 | 2.04 | 0.0015 |
| (-) | LPC 16:1 | C24H48NO7P | 493.314 | 552.3279 | 6 | 1.92 | 2.82 | 0.0162 |
| (-) | LPC 17:0 | C25H52NO7P | 509.3481 | 568.362 | 0 | 3.22 | 2.63 | 0.0187 |
| (-) | LPC 18:0 | C26H54NO7P | 523.3637 | 582.3776 | 0 | 3.22 | 2.76 | 0.0246 |
| (+) | LPC 18:1 | C26H52NO7P | 521.3484 | 522.3557 | 1 | 2.73 | 1.91 | 0.0035 |
| (-) | LPC 18:3/0:0 | C26H48NO7P | 517.316 | 576.3299 | 2 | 2.68 | 6.41 | 0.0028 |
| (+) | LPC 20:1 | C28H56NO7P | 549.3791 | 550.3864 | 1 | 3.53 | 2.16 | 0.0467 |
| (+) | LPC 20:2/0:0 | C30H52NO7P | 569.345 | 570.3523 | 5 | 2.53 | 6.13 | 0.0262 |
| (-) | LPC 20:3 | C28H52NO7P | 545.3469 | 604.3608 | 2 | 2.72 | 1.13 | 0.0023 |
| (-) | LPC 20:4 | C28H50NO7P | 543.332 | 602.3459 | 1 | 2.12 | 2.10 | 0.0187 |
| (+) | LPC 22:0/0:0 | C30H62NO7P | 579.4266 | 580.4339 | 0 | 4.9 | 5.19 | 0.0379 |
| (-) | LPC 22:6 | C30H50NO7P | 567.3348 | 626.3487 | 4 | 2.04 | 1.51 | 0.0315 |
| (+) | LPC O-16:0 | C24H52NO6P | 481.3534 | 482.3607 | 0 | 2.99 | 2.79 | 0.0030 |
| (+) | LPC O-18:1 | C26H54NO6P | 507.3688 | 508.3761 | 0 | 3.14 | 4.52 | 0.0262 |
| (-) | LPE 0:0/16:0 | C21H44NO7P | 453.2867 | 452.2794 | 3 | 2.46 | 3.47 | 0.0015 |
| (-) | LPE 16:0/0:0 | C21H44NO7P | 453.2861 | 452.2788 | 1 | 2.66 | 2.73 | 0.0162 |
| (+) | LPE 18:1 | C23H46NO7P | 479.3004 | 480.3077 | 2 | 2.85 | 5.08 | 0.0306 |

| | | | | | | | | |
|---|---|---|---|---|---|---|---|---|
| (-) | LPE 20:4 | C25H44NO7P | 501.2853 | 500.278 | 1 | 2.22 | 2.91 | 0.0028 |
| (-) | LPE 22:6 | C27H44NO7P | 525.2851 | 524.2778 | 1 | 2.14 | 1.81 | 0.0039 |
| (-) | LPE O-16:1 | C21H44NO6P | 437.2913 | 436.284 | 1 | 2.84 | 9.25 | 0.0226 |
| (-) | LPI 18:0 | C27H53O12P | 600.3228 | 599.3155 | 8 | 2.56 | 4.29 | 0.0156 |
| (-) | LPI 18:1 | C27H51O12P | 598.3139 | 597.3066 | 3 | 0.69 | 3.43 | 0.0427 |
| (-) | LPI 20:3 | C29H51O12P | 622.3111 | 621.3038 | 1 | 1.95 | 3.76 | 0.0107 |
| (+) | PC 12:0_16:0 | C36H72NO8P | 677.498 | 678.5053 | 2 | 5.9 | 4.74 | 0.0157 |
| (-) | PC 14:0_18:2 | C40H76NO8P | 729.531 | 788.5449 | 0 | 6.36 | 3.17 | 0.0039 |
| (-) | PC 16:0_16:0 | C40H80NO8P | 733.562 | 792.5759 | 0 | 8.89 | 2.88 | 0.0396 |
| (-) | PC 16:0_16:1 | C40H78NO8P | 731.5462 | 790.5601 | 0 | 7.37 | 5.57 | 0.0455 |
| (-) | PC 16:0_18:1 | C42H82NO8P | 759.5761 | 818.59 | 2 | 9.19 | 3.60 | 0.0498 |
| (+) | PC 16:0_18:2 | C42H80NO8P | 757.5633 | 758.5706 | 2 | 7.61 | 2.45 | 0.0061 |
| (-) | PC 16:0_20:4 | C44H81NO8P | 781.5619 | 826.5601 | 0 | 7.59 | 2.95 | 0.0315 |
| (-) | PC 16:0_22:5 | C46H82NO8P | 807.5673 | 852.5655 | 13 | 7.76 | 5.24 | 0.0246 |
| (-) | PC 18:0_18:2 | C44H84NO8P | 785.5934 | 830.5916 | 0 | 9.86 | 2.59 | 0.0107 |
| (-) | PC 18:1_18:1 | C44H84NO8P | 785.5879 | 844.6018 | 7 | 9.36 | 8.89 | 0.0119 |
| (-) | PC 18:2_20:1 | C46H86NO8P | 811.608 | 870.6219 | 1 | 10.61 | 3.16 | 0.0220 |
| (-) | PC 19:0_18:1 | C45H88NO8P | 801.6308 | 860.6447 | 8 | 11.75 | 2.99 | 0.0057 |
| (-) | PC 33:3 | C41H76NO8P | 741.5312 | 800.5451 | 1 | 6.56 | 2.56 | 0.0389 |
| (-) | PC 34:3 | C42H78NO8P | 755.5454 | 814.5593 | 1 | 6.82 | 5.15 | 0.0076 |
| (+) | PC 35:2 | C43H82NO8P | 771.5798 | 772.5871 | 3 | 8.45 | 6.96 | 0.0145 |
| (-) | PC 36:4_iso1 | C44H80NO8P | 781.5583 | 840.5722 | 5 | 6.89 | 2.87 | 0.0075 |
| (+) | PC 36:4_iso2 | C44H80NO8P | 781.5613 | 782.5686 | 1 | 7.61 | 1.81 | 0.0201 |
| (-) | PC 36:5_iso1 | C44H78NO8P | 779.5393 | 838.5532 | 9 | 6.08 | 6.91 | 0.0363 |
| (-) | PC 36:5_iso2 | C44H78NO8P | 779.5555 | 838.5694 | 12 | 6.38 | 3.99 | 0.0218 |
| (+) | PC 36:6 | C44H76NO8P | 777.5316 | 778.5389 | 1 | 6.03 | 3.48 | 0.0262 |
| (+) | PC 37:2 | C45H86NO8P | 799.6082 | 800.6155 | 1 | 10.8 | 3.95 | 0.0254 |
| (+) | PC 37:3 | C45H84NO8P | 797.5923 | 798.5996 | 1 | 8.94 | 4.14 | 0.0250 |
| (+) | PC 37:4 | C45H82NO8P | 795.5773 | 796.5846 | 1 | 8.52 | 6.80 | 0.0046 |
| (+) | PC 38:2 | C46H88NO8P | 813.6263 | 814.6336 | 2 | 11.51 | 5.55 | 0.0293 |
| (+) | PC 38:6 | C46H80NO8P | 805.5635 | 806.5708 | 2 | 7.3 | 1.67 | 0.0262 |
| (-) | PC 38:7 | C46H78NO8P | 803.5467 | 862.5606 | 0 | 6.19 | 4.46 | 0.0023 |
| (+) | PC 39:6 | C47H82NO8P | 819.5715 | 820.5788 | 8 | 7.88 | 7.14 | 0.0006 |
| (+) | PC 40:6 | C48H84NO8P | 833.5932 | 834.6005 | 0 | 8.4 | 18.02 | 0.0190 |
| (+) | PC 40:7 | C48H82NO8P | 831.5785 | 832.5858 | 1 | 7.47 | 3.66 | 0.0180 |

| | | | | | | | | |
|---|---|---|---|---|---|---|---|---|
| (+) | PC O-32:1 | C40H80NO7P | 717.5648 | 718.5721 | 3 | 8.42 | 5.28 | 0.0487 |
| (+) | PC O-34:0 | C42H86NO7P | 747.6254 | 748.6327 | 15 | 12.32 | 8.09 | 0.0262 |
| (-) | PC O-35:4 | C43H80NO7P | 753.5672 | 812.5811 | 0 | 8.74 | 12.14 | 0.0434 |
| (+) | PC O-36:5 | C44H80NO7P | 765.5661 | 766.5734 | 1 | 8.44 | 4.50 | 0.0434 |
| (-) | PC O-37:5 | C45H82NO7P | 779.5731 | 838.587 | 12 | 8.89 | 3.82 | 0.0406 |
| (+) | PC O-38:6 | C46H82NO7P | 791.5821 | 792.5894 | 1 | 8.36 | 2.83 | 0.0318 |
| (+) | PC O-38:7 | C46H80NO7P | 789.5682 | 790.5755 | 1 | 8.05 | 4.49 | 0.0467 |
| (+) | PE 16:0_22:6 | C43H74NO8P | 763.515 | 764.5223 | 0 | 7.63 | 2.35 | 0.0189 |
| (-) | PE 18:0_18:2 | C41H78NO8P | 743.5458 | 742.5385 | 1 | 10.39 | 1.74 | 0.0162 |
| (-) | PE 38:5 | C43H76NO8P | 765.53 | 764.5227 | 1 | 8.16 | 1.85 | 0.0107 |
| (-) | PE O-16:1_22:6 // PE P-16:0_22:6 | C43H74NO7P | 747.5199 | 746.5126 | 1 | 8.48 | 5.80 | 0.0328 |
| (-) | PE O-18:2_22:6 // PE P-18:1_22:6 | C45H76NO7P | 773.5324 | 772.5251 | 5 | 8.71 | 5.26 | 0.0162 |
| (-) | PI 16:0_16:1 | C41H77O13P | 808.5145 | 807.5072 | 5 | 5.84 | 2.51 | 0.0414 |
| (-) | PI 16:0_20:4 | C45H79O13P | 858.5257 | 857.5184 | 0 | 6 | 2.33 | 0.0187 |
| (-) | PI 16:0_22:5 | C47H81O13P | 884.5411 | 883.5338 | 0 | 6.13 | 2.35 | 0.0098 |
| (-) | PI 18:0_20:3 | C47H85O13P | 888.5714 | 887.5641 | 2 | 8.31 | 1.70 | 0.0363 |
| (-) | PI 18:0_20:4 | C47H83O13P | 886.5581 | 885.5508 | 1 | 7.22 | 4.10 | 0.0363 |
| (-) | PI 18:1_18:2 | C45H81O13P | 860.5407 | 859.5334 | 1 | 6.25 | 3.41 | 0.0175 |
| (-) | PI 36:2 | C45H83O13P | 862.5569 | 861.5496 | 0 | 7.37 | 5.06 | 0.0134 |
| (-) | PS 18:2_24:0 | C48H90NO10P | 871.6291 | 870.6218 | 1 | 10.62 | 6.19 | 0.0359 |
| (-) | PS 20:0_20:4 | C46H82NO10P | 839.5691 | 838.5618 | 2 | 6.4 | 4.52 | 0.0076 |
| (-) | SM 18:1;O2/14:0 | C37H75N2O6P | 674.5243 | 733.5382 | 18 | 5.83 | 2.28 | 0.0106 |
| (-) | SM 18:1;O2/15:0 | C38H77N2O6P | 688.5516 | 747.5655 | 0 | 7.08 | 2.51 | 0.0213 |
| (-) | SM 18:1;O2/17:0 | C40H81N2O6P | 716.5834 | 775.5973 | 0 | 8.87 | 4.56 | 0.0107 |
| (-) | SM 18:1;O2/22:1 | C45H89N2O6P | 784.6438 | 843.6577 | 3 | 11.33 | 2.64 | 0.0189 |
| (-) | SM 18:1;O2/23:0 | C46H93N2O6P | 800.6735 | 859.6874 | 4 | 12.039 | 1.64 | 0.0030 |
| (+) | SM 18:1;O2/24:2 | C47H91N2O6P | 810.6623 | 811.6696 | 1 | 11.53 | 2.86 | 0.0074 |
| (-) | SM 18:2;O2/14:0 | C37H73N2O6P | 672.5187 | 731.5326 | 3 | 5.11 | 6.51 | 0.0250 |
| (-) | SM 18:2;O2/15:0 | C38H75N2O6P | 686.5354 | 745.5493 | 1 | 6.05 | 3.39 | 0.0098 |
| (-) | SM 18:2;O2/16:0 | C39H77N2O6P | 700.5511 | 759.565 | 1 | 6.05 | 4.02 | 0.0107 |
| (-) | SM 18:2;O2/22:0 | C45H89N2O6P | 784.6436 | 843.6575 | 3 | 11.51 | 2.36 | 0.0397 |
| (+) | SM 18:2;O2/22:1 | C45H87N2O6P | 782.6286 | 783.6359 | 2 | 9.48 | 6.39 | 0.0128 |
| (+) | SM 18:2;O2/23:0 | C46H91N2O6P | 798.6604 | 799.6677 | 1 | 11.72 | 2.63 | 0.0047 |
| (-) | SM 18:2;O2/25:0 | C48H95N2O6P | 826.6899 | 885.7038 | 3 | 12.03 | 6.24 | 0.0113 |
| (+) | SM 33:1;O2 | C38H77N2O6P | 688.5511 | 689.5584 | 1 | 6.43 | 3.17 | 0.0199 |

| | | | | | | | | |
|---|---|---|---|---|---|---|---|---|
| (+) | SM 34:0;O2 | C39H81N2O6P | 704.5829 | 705.5902 | 0 | 7.74 | 3.49 | 0.0218 |
| (+) | SM 34:1;O2 | C39H79N2O6P | 702.5682 | 703.5755 | 1 | 7.11 | 2.53 | 0.0282 |
| (+) | SM 35:1;O2 | C40H81N2O6P | 716.581 | 717.5883 | 3 | 7.93 | 3.33 | 0.0270 |
| (-) | SM 36:1;O2 | C41H83N2O6P | 730.5977 | 789.6116 | 2 | 8.87 | 3.64 | 0.0107 |
| (+) | SM 36:2;O2 | C41H81N2O6P | 728.5829 | 729.5902 | 0 | 7.39 | 4.49 | 0.0307 |
| (+) | SM 39:1;O2 | C44H89N2O6P | 772.6418 | 773.6491 | 5 | 11.71 | 4.23 | 0.0499 |
| (+) | SM 42:1;O2 | C47H95N2O6P | 814.6922 | 815.6995 | 1 | 12.17 | 3.40 | 0.0128 |
| (+) | SM 42:2;O2 | C47H93N2O6P | 812.6777 | 813.685 | 1 | 11.9 | 2.44 | 0.0157 |
| (+) | TG 10:0_10:0_12:0 | C35H66O6 | 582.4811 | 600.5149 | 8 | 11.58 | 18.39 | 0.0374 |
| (+) | TG 10:0_12:0_12:0 | C37H70O6 | 610.5194 | 669.5798 | 4 | 11.99 | 7.25 | 0.0103 |
| (+) | TG 12:0_12:0_18:2 | C45H82O6 | 718.6116 | 736.6454 | 1 | 12.6 | 4.17 | 0.0145 |
| (+) | TG 16:0_18:1_18:1 | C55H102O6 | 858.7687 | 876.8025 | 1 | 14.54 | 2.47 | 0.0128 |
| (+) | TG 16:0_18:1_22:5 | C59H102O6 | 906.7512 | 965.8116 | 18 | 14.23 | 26.02 | 0.0201 |
| (+) | TG 36:0 | C39H74O6 | 638.5489 | 661.5381 | 1 | 12.25 | 4.34 | 0.0075 |
| (+) | TG 38:0 | C41H78O6 | 666.5805 | 684.6143 | 1 | 12.54 | 4.21 | 0.0045 |
| (+) | TG 40:0 | C43H82O6 | 694.6115 | 712.6453 | 0 | 12.85 | 4.84 | 0.0088 |
| (+) | TG 40:1 | C43H80O6 | 692.5948 | 710.6286 | 1 | 12.56 | 4.99 | 0.0128 |
| (+) | TG 40:2 | C43H78O6 | 690.5796 | 708.6134 | 0 | 12.33 | 5.26 | 0.0298 |
| (+) | TG 42:0 | C45H86O6 | 722.6428 | 740.6766 | 0 | 13.19 | 3.75 | 0.0262 |
| (+) | TG 43:0 | C46H88O6 | 736.6571 | 754.6909 | 1 | 13.38 | 22.66 | 0.0198 |
| (+) | TG 44:0 | C47H90O6 | 750.6634 | 768.6972 | 14 | 13.19 | 7.02 | 0.0367 |
| (+) | TG 44:1 | C47H88O6 | 748.6579 | 766.6917 | 0 | 13.19 | 3.51 | 0.0306 |
| (+) | TG 44:2 | C47H86O6 | 746.6429 | 764.6767 | 1 | 12.9 | 4.48 | 0.0436 |
| (+) | TG 45:0 | C48H92O6 | 764.6886 | 782.7224 | 1 | 13.8 | 16.61 | 0.0446 |
| (+) | TG 48:2 | C51H94O6 | 802.7058 | 820.7396 | 1 | 13.62 | 4.23 | 0.0423 |
| (+) | TG 48:6 | C51H86O6 | 794.6409 | 812.6747 | 2 | 12.67 | 5.49 | 0.0417 |
| (+) | TG 50:2 | C53H98O6 | 830.7306 | 848.7644 | 7 | 14.02 | 3.86 | 0.0180 |
| (+) | TG 51:0 | C54H104O6 | 848.7817 | 866.8155 | 2 | 15.6 | 9.23 | 0.0396 |
| (+) | TG 52:1 | C55H104O6 | 860.7814 | 883.7706 | 2 | 15.19 | 24.98 | 0.0496 |
| (+) | TG 53:3 | C56H102O6 | 870.7755 | 888.8093 | 9 | 14.28 | 8.74 | 0.0467 |
| (+) | TG 54:2 | C57H106O6 | 886.8002 | 904.834 | 1 | 15.15 | 5.91 | 0.0262 |
| (+) | TG 56:3 | C59H108O6 | 912.808 | 930.8418 | 7 | 14.54 | 4.44 | 0.0467 |
| (+) | TG 56:4 | C59H106O6 | 910.7984 | 928.8322 | 1 | 14.67 | 11.74 | 0.0360 |

**Table S2. Lipids differentially expressed between passively-sensitized mice at 7 days post-allergen challenge.**

| Control > IgE | | |
|---|---|---|
| ***Metabolite*** | ***Log2(Fold Change)*** | ***P-value*** |
| NAGlySer 11:0;O(FA 28:6) | -3.845 | 0.000 |
| NAGlySer 11:0;O(FA 28:5) | -3.362 | 0.000 |
| Cer 12:0;O2/37:0;O | -1.792 | 0.004 |
| Cer 19:1;O3/38:1(2OH) | -1.716 | 0.015 |
| Cer 21:0;O3/38:0(2OH) | -1.588 | 0.019 |
| HexCer 16:0;O2/24:3;O | -1.513 | 0.004 |
| TG O-18:0_16:1_16:1 | -1.512 | 0.029 |
| Cer 21:1;O3/38:2(2OH) | -1.507 | 0.012 |
| SE 28:2/18:0 B | -1.491 | 0.013 |
| Cer 12:0;O2/38:0;O A | -1.485 | 0.005 |
| SL 12:1;O/35:1;O | -1.481 | 0.024 |
| Cer 17:1;O3/38:2(2OH) | -1.468 | 0.004 |
| TG O-16:0_16:0_18:2 | -1.458 | 0.002 |
| TG O-52:1\|TG O-18:0_16:0_18:1 | -1.451 | 0.017 |
| Cer 12:0;O2/24:1;O | -1.437 | 0.025 |
| DG 18:0_34:0 | -1.429 | 0.023 |
| TG 44:0\|TG 12:0_14:0_18:0 | -1.386 | 0.005 |
| Cer 13:1;O3/36:4(2OH) | -1.373 | 0.000 |
| DG 18:0_36:0 | -1.348 | 0.005 |
| TG 8:0_8:0_28:2 A | -1.343 | 0.017 |
| CE 20:1 | -1.313 | 0.027 |
| SE 24:1;O4/26:1;O | -1.304 | 0.020 |
| TG 46:0\|TG 12:0_16:0_18:0 | -1.303 | 0.011 |
| SL 12:1;O/36:0;O | -1.301 | 0.002 |
| TG 14:0_16:0_18:0 | -1.297 | 0.014 |
| TG 48:0\|TG 14:0_16:0_18:0 | -1.29 | 0.018 |
| TG O-17:0_18:0_17:1 | -1.289 | 0.007 |
| ST 27:1;O C | -1.277 | 0.016 |
| Cer 12:0;O2/36:0;O | -1.276 | 0.012 |
| TG O-16:1_18:0_20:0 | -1.271 | 0.036 |
| NAGly 20:0;O(FA 28:1) | -1.253 | 0.001 |
| TG 42:0\|TG 12:0_14:0_16:0 | -1.248 | 0.006 |
| TG O-54:2\|TG O-18:0_16:1_20:1 | -1.233 | 0.037 |
| TG 8:0_8:0_26:0 | -1.232 | 0.000 |
| TG 8:0_8:0_38:0 B | -1.227 | 0.003 |
| PI 36:1 | -1.226 | 0.001 |
| TG 14:0_16:0_17:0 | -1.222 | 0.006 |
| TG O-16:1_17:0_17:0 | -1.219 | 0.027 |
| PC 38:4\|PC 18:0_20:4 A | -1.214 | 0.036 |
| TG O-42:0\|TG O-16:0_12:0_14:0 | -1.207 | 0.024 |
| TG O-46:1\|TG O-16:0_12:0_18:1 | -1.203 | 0.024 |
| TG 40:0\|TG 10:0_14:0_16:0 | -1.201 | 0.005 |
| DG 18:0_28:0 | -1.196 | 0.017 |
| Cer 23:0;O3/38:0(2OH) | -1.188 | 0.041 |
| TG 18:0_18:0_20:0 | -1.185 | 0.045 |

| | | |
|---|---|---|
| TG 8:0_8:0_26:1 | -1.182 | 0.010 |
| Cer 13:1;O3/38:0(2OH) | -1.181 | 0.039 |
| TG O-21:1_17:0_18:0 | -1.176 | 0.027 |
| TG O-50:1\|TG O-16:0_16:0_18:1 | -1.171 | 0.021 |
| TG 14:0_19:2_19:2 | -1.162 | 0.010 |
| TG O-20:0_12:0_18:0 | -1.156 | 0.009 |
| Cer 46:2;O2\|Cer 18:1;O2/28:1 | -1.149 | 0.026 |
| TG 13:0_13:0_18:0 | -1.144 | 0.007 |
| Cer 21:1;O3/38:1(2OH) A | -1.144 | 0.020 |
| SL 12:0;O/36:0;O | -1.139 | 0.004 |
| Cer 13:0;O2/30:4;O | -1.139 | 0.040 |
| TG 12:0_18:0_22:0 | -1.136 | 0.015 |
| TG O-46:2\|TG O-16:0_12:0_18:2 | -1.122 | 0.046 |
| TG 38:0\|TG 12:0_12:0_14:0 | -1.121 | 0.023 |
| NAGly 26:0;O(FA 28:2) | -1.106 | 0.023 |
| TG 8:0_8:0_32:0 | -1.101 | 0.007 |
| Cer 25:1;O2/22:3;O | -1.099 | 0.050 |
| ST 27:2;O B | -1.088 | 0.001 |
| TG 14:0_16:0_16:0 | -1.087 | 0.004 |
| TG 40:2\|TG 10:0_12:0_18:2 | -1.086 | 0.011 |
| DG 50:0 | -1.086 | 0.032 |
| TG 10:0_12:0_24:1 | -1.083 | 0.002 |
| SE 24:1;O4/25:1;O B | -1.078 | 0.001 |
| TG 11:0_15:0_19:1 | -1.078 | 0.020 |
| TG 40:1\|TG 10:0_12:0_18:1 | -1.074 | 0.050 |
| TG 8:0_8:0_28:0 | -1.072 | 0.015 |
| TG O-46:0\|TG O-18:0_12:0_16:0 | -1.07 | 0.038 |
| TG 44:1\|TG 12:0_14:0_18:1 | -1.066 | 0.019 |
| TG 13:0_18:0_18:0 | -1.058 | 0.003 |
| TG 14:0_18:1_18:1 | -1.056 | 0.011 |
| TG 13:0_18:0_20:0 | -1.055 | 0.035 |
| TG 46:2\|TG 12:0_16:0_18:2 | -1.05 | 0.017 |
| NAE 22:5 | -1.046 | 0.001 |
| TG 52:0\|TG 16:0_18:0_18:0 | -1.044 | 0.045 |
| Cer 50:2;O2\|Cer 20:0;O2/30:2 | -1.037 | 0.009 |
| TG 50:0\|TG 16:0_16:0_18:0 | -1.034 | 0.013 |
| Cer 13:1;O3/38:1(2OH) | -1.034 | 0.035 |
| DG 42:0 | -1.03 | 0.006 |
| TG 8:0_8:0_28:2 C | -1.03 | 0.024 |
| TG 12:0_16:0_20:4 | -1.025 | 0.009 |
| DG 43:8 | -1.025 | 0.036 |
| TG 8:0_9:0_34:4 | -1.02 | 0.024 |
| TG 8:0_9:0_26:2 | -1.018 | 0.032 |
| TG 14:0_16:0_16:2 | -1.016 | 0.006 |
| Cer 47:2;O2\|Cer 20:1;O2/27:1 | -1.01 | 0.014 |
| TG 44:2\|TG 12:0_14:0_18:2 | -1.009 | 0.027 |
| Cer 18:1;O3/38:0(2OH) | -1.009 | 0.030 |
| TG 8:0_8:0_26:2 | -1.009 | 0.036 |
| DG 50:1 | -1.007 | 0.002 |

| | | |
|---|---|---|
| Cer 12:0;O2/38:0;O B | -1.004 | 0.029 |
| TG 8:0_8:0_24:0 | -1.003 | 0.008 |
| NAOrn 27:1 A | -1.002 | 0.025 |
| TG 42:1\|TG 12:0_12:0_18:1 | -0.999 | 0.010 |
| Cer 13:0;O2/32:4;O | -0.994 | 0.022 |
| TG 10:0_15:0_20:2 | -0.992 | 0.036 |
| NAE 20:2 | -0.989 | 0.001 |
| VAE 27:0 | -0.987 | 0.002 |
| SE 28:1/15:0 | -0.987 | 0.013 |
| TG 14:0_16:0_20:0 | -0.982 | 0.002 |
| TG 16:0_16:0_18:1 | -0.982 | 0.016 |
| Cer 15:3;O2/9:0 | -0.97 | 0.016 |
| CE 17:1 | -0.969 | 0.041 |
| TG 13:1_15:1_19:4 | -0.966 | 0.007 |
| TG O-13:0_13:0_18:0 | -0.963 | 0.033 |
| TG 8:0_8:0_32:4 | -0.961 | 0.007 |
| TG 46:1\|TG 12:0_16:0_18:1 | -0.958 | 0.021 |
| TG 11:0_11:0_21:0 | -0.957 | 0.021 |
| TG 48:2\|TG 14:0_16:0_18:2 | -0.952 | 0.028 |
| Cer 12:1;O3/35:0(2OH) | -0.946 | 0.036 |
| TG 12:0_16:0_18:3 | -0.945 | 0.020 |
| NAOrn 19:0;O B | -0.944 | 0.015 |
| ST 27:2;O E | -0.94 | 0.049 |
| AHexCer (O-12:0)12:2;O2/12:0;O A | -0.939 | 0.029 |
| TG 8:0_8:0_30:0 | -0.929 | 0.004 |
| Cer 12:0;O2/28:1;O | -0.929 | 0.050 |
| Cer 14:0;O3/38:0(2OH) | -0.925 | 0.024 |
| TG 48:1\|TG 14:0_16:0_18:1 | -0.923 | 0.033 |
| TG 8:0_8:0_30:3 | -0.92 | 0.030 |
| Cer 19:1;O3/38:2(2OH) | -0.917 | 0.045 |
| ST 28:1;O B | -0.913 | 0.041 |
| TG 8:0_8:0_36:0 | -0.912 | 0.002 |
| SL 17:0;O/36:0;O A | -0.912 | 0.032 |
| VAE 21:0 | -0.906 | 0.022 |
| TG 54:1\|TG 18:0_18:0_18:1 | -0.905 | 0.033 |
| TG 8:0_8:0_29:1 | -0.903 | 0.016 |
| TG 8:0_18:0_22:2 | -0.9 | 0.016 |
| TG O-50:0\|TG O-18:0_14:0_18:0 | -0.898 | 0.024 |
| CE 18:3 | -0.897 | 0.039 |
| TG O-40:0\|TG O-18:0_10:0_12:0 | -0.886 | 0.018 |
| Cer 12:0;O2/34:0;O | -0.885 | 0.033 |
| HexCer 16:0;O2/19:5 | -0.883 | 0.012 |
| DG 51:10 | -0.883 | 0.038 |
| NAGly 13:0;O(FA 9:0) | -0.881 | 0.011 |
| SM 12:1;O2/28:0 | -0.873 | 0.013 |
| DG 52:1 | -0.868 | 0.026 |
| Cer 12:1;O3/38:3(2OH) | -0.866 | 0.037 |
| SL 16:0;O/36:0;O | -0.862 | 0.039 |
| TG 8:0_8:0_30:1 A | -0.86 | 0.023 |

| | | |
|---|---|---|
| TG 36:0\|TG 10:0_12:0_14:0 | -0.859 | 0.007 |
| Cer 53:3;O2\|Cer 17:3;O2/36:0 | -0.857 | 0.015 |
| Cer 48:3;O2\|Cer 16:2;O2/32:1 | -0.851 | 0.017 |
| Cer 23:1;O3/38:1(2OH) | -0.846 | 0.034 |
| TG O-44:0\|TG O-18:0_12:0_14:0 | -0.845 | 0.038 |
| TG 42:2\|TG 12:0_12:0_18:2 | -0.842 | 0.044 |
| TG 8:0_8:0_30:2 | -0.84 | 0.013 |
| TG O-48:1\|TG O-18:0_12:0_18:1 | -0.837 | 0.036 |
| SM 35:1;O3 | -0.834 | 0.030 |
| TG 17:0_13:1_17:1 | -0.829 | 0.024 |
| TG 8:0_8:0_34:3 | -0.82 | 0.025 |
| TG 8:0_8:0_24:1 | -0.819 | 0.016 |
| SL 18:0;O/36:0;O | -0.819 | 0.040 |
| DG 46:0 | -0.805 | 0.033 |
| SM 37:3;O3 | -0.803 | 0.046 |
| Cer 22:1;O3/38:0(2OH) | -0.8 | 0.022 |
| SL 16:1;O/36:1;O B | -0.788 | 0.002 |
| DG 51:6 | -0.788 | 0.039 |
| TG 48:4\|TG 12:0_18:2_18:2 | -0.784 | 0.036 |
| TG 50:4\|TG 14:0_18:1_18:3 | -0.781 | 0.030 |
| TG 45:0\|TG 14:0_15:0_16:0 | -0.781 | 0.037 |
| DG 44:0 | -0.776 | 0.039 |
| SM 39:3;O3 | -0.774 | 0.032 |
| TG 12:0_18:0_18:2 | -0.771 | 0.008 |
| TG 8:0_8:0_28:1 | -0.762 | 0.020 |
| TG 8:0_8:0_24:2 C | -0.762 | 0.040 |
| DG 52:0 | -0.76 | 0.008 |
| DG 46:1 | -0.76 | 0.011 |
| TG 17:0_15:1_17:1 | -0.756 | 0.008 |
| PE 21:1 | -0.756 | 0.018 |
| TG 17:0_15:1_19:1 | -0.752 | 0.030 |
| DG 54:2 | -0.749 | 0.041 |
| PC 40:6 B | -0.742 | 0.007 |
| DG 54:1 | -0.74 | 0.003 |
| NAGly 26:2 A | -0.739 | 0.028 |
| TG 8:0_8:0_38:0 A | -0.718 | 0.009 |
| TG 48:3\|TG 12:0_18:1_18:2 | -0.715 | 0.043 |
| Cer 16:0;O3/38:0(2OH) | -0.711 | 0.036 |
| TG 8:0_8:0_24:2 B | -0.697 | 0.018 |
| TG 16:0_16:0_19:1 | -0.69 | 0.016 |
| Cer 12:0;O3/8:0(2OH) | -0.685 | 0.006 |
| Cer 12:0;O3/38:0(2OH) | -0.679 | 0.021 |
| Cer 25:1;O3/38:0(2OH) | -0.679 | 0.039 |
| TG 50:6\|TG 12:0_16:0_22:6 | -0.678 | 0.029 |
| TG 8:0_8:0_38:1 | -0.674 | 0.019 |
| TG 8:0_8:0_30:4 | -0.67 | 0.011 |
| TG 18:0_14:1_20:2 | -0.667 | 0.031 |
| SM 12:1;O2/23:0 | -0.66 | 0.046 |
| SM 41:3;O3 | -0.653 | 0.011 |

| | | |
|---|---|---|
| TG 8:0_8:0_36:1 B | -0.652 | 0.040 |
| NAGly 24:2 | -0.651 | 0.000 |
| TG 8:0_8:0_32:6 A | -0.649 | 0.041 |
| SM 12:0;O2/19:0 | -0.644 | 0.016 |
| DG 47:7 | -0.64 | 0.033 |
| SL 13:0;O/36:0;O | -0.636 | 0.002 |
| SM 30:3;O3 | -0.636 | 0.037 |
| TG 8:0_8:0_22:0 B | -0.635 | 0.045 |
| SL 13:2;O/36:6 | -0.633 | 0.016 |
| Cer 13:0;O2/34:3;O | -0.632 | 0.007 |
| LPC 20:3/0:0 | -0.63 | 0.020 |
| TG 8:0_8:0_22:0 C | -0.627 | 0.031 |
| SE 28:1/19:0 | -0.621 | 0.007 |
| TG 54:6\|TG 18:1_18:2_18:3 | -0.609 | 0.047 |
| NAPhe 26:6;O | -0.605 | 0.003 |
| TG 8:0_8:0_37:1 | -0.601 | 0.041 |
| Cer 42:3;O3\|Cer 18:1;O2/24:2;O | -0.592 | 0.041 |
| TG 11:0_18:0_20:1 | -0.588 | 0.042 |
| BMP 13:1_28:7 | -0.586 | 0.032 |

| **IgE > Control** | | |
|---|---|---|
| ***Metabolite*** | ***Log2(Fold Change)*** | ***P-value*** |
| VAE 25:1 B | 1.802 | 0.014 |
| TG 9:0_9:0_20:1 | 1.792 | 0.020 |
| SE 28:2/20:1 | 1.436 | 0.021 |
| PS 12:0 | 1.346 | 0.020 |
| TG 8:0_9:0_38:5 | 0.947 | 0.006 |
| Cer 46:4;O2\|Cer 30:3;O2/16:1 | 0.798 | 0.007 |
| NAGly 13:1 B | 0.79 | 0.005 |
| TG 45:2\|TG 12:0_18:0_15:2 | 0.78 | 0.000 |
| TG 8:0_9:0_30:2 A | 0.77 | 0.003 |
| ST 27:1;O G | 0.751 | 0.000 |

| **Control > IgE + IgG** | | |
|---|---|---|
| ***Metabolite*** | ***Log2(Fold Change)*** | ***P-value*** |
| DG 43:8 | -0.963 | 0.016 |
| SE 28:1/15:0 | -0.85 | 0.022 |
| DG 52:1 | -0.845 | 0.038 |
| Cer 18:1;O3/38:1(2OH) | -0.778 | 0.007 |
| Cer 15:3;O2/9:0 | -0.751 | 0.045 |
| DG 54:1 | -0.663 | 0.016 |

| **IgE + IgG > Control** | | |
|---|---|---|
| ***Metabolite*** | ***Log2(Fold Change)*** | ***P-value*** |
| LPC 12:0 | 3.065 | 0.023 |
| TG 8:0_9:0_22:2 | 2.328 | 0.014 |
| SM 38:4;O3 | 2.065 | 0.037 |
| Cer 46:4;O2\|Cer 30:3;O2/16:1 | 1.488 | 0.010 |
| NATau 20:0;O | 1.457 | 0.002 |

| | | |
|---|---|---|
| TG 8:0_9:0_30:2 A | 1.304 | 0.003 |
| Cer 21:1;O3/38:1(2OH) B | 1.194 | 0.024 |
| NATau 22:3;O | 1.191 | 0.011 |
| TG 8:0_9:0_28:2 | 1.098 | 0.044 |
| SM 36:1;O2 A | 1.086 | 0.050 |
| NAGly 13:1 B | 1.007 | 0.015 |
| TG 43:2\|TG 12:0_16:0_15:2 | 0.945 | 0.006 |
| NATau 17:1;O A | 0.894 | 0.016 |
| Cer 13:2;O2/38:6 | 0.872 | 0.041 |
| SE 28:2/18:1 | 0.822 | 0.047 |
| DG 34:1 | 0.765 | 0.026 |
| Cer 13:0;O2/24:2;O | 0.748 | 0.043 |
| PI 38:4 | 0.686 | 0.021 |
| Cer 13:0;O2/34:3 | 0.649 | 0.013 |
| NAGly 18:1 B | 0.647 | 0.038 |
| Cer 12:1;O2/22:6 | 0.64 | 0.022 |
| Cer 16:3;O2/16:3 | 0.639 | 0.009 |
| Cer 12:0;O2/28:0 | 0.63 | 0.017 |
| HexCer 18:1;O2/22:0 | 0.622 | 0.039 |
| SM 34:0;O3 | 0.621 | 0.034 |
| Cer 12:1;O3/24:5(2OH) B | 0.602 | 0.008 |

| **Control > IgG** | | |
|---|---|---|
| ***Metabolite*** | ***Log2(Fold Change)*** | ***P-value*** |
| NAGlySer 11:0;O(FA 28:6) | -1.572 | 0.007 |
| NAGlySer 11:0;O(FA 28:5) | -0.754 | 0.014 |
| NAGlySer 18:2;O H | -0.634 | 0.027 |
| DG 31:3 | -0.625 | 0.010 |
| NAGly 28:6 E | -0.608 | 0.033 |

| **IgG > Control** | | |
|---|---|---|
| ***Metabolite*** | ***Log2(Fold Change)*** | ***P-value*** |
| PC O-38:5\|PC O-18:1_20:4 | 4.831 | 0.003 |
| SM 38:4;O3 | 4.5 | 0.007 |
| PC 36:4\|PC 16:0_20:4 E | 4.256 | 0.003 |
| SM 34:1;O2\|SM 18:1;O2/16:0 B | 4.247 | 0.005 |
| LPC 12:0 | 4.226 | 0.016 |
| PC 38:6\|PC 16:0_22:6 C | 3.763 | 0.006 |
| SM 36:1;O2 C | 3.633 | 0.000 |
| SE 24:1;O4/19:1;O | 3.535 | 0.046 |
| PC 38:4\|PC 18:0_20:4 B | 3.466 | 0.003 |
| Hex2Cer 16:0;O2/14:0 | 3.414 | 0.003 |
| PC O-36:4\|PC O-16:0_20:4 | 3.259 | 0.000 |
| SM 36:1;O2 A | 3.148 | 0.000 |
| PC 32:0\|PC 16:0_16:0 A | 3.123 | 0.022 |
| PC O-38:5\|PC O-20:5_18:0 | 3.065 | 0.035 |
| PC 36:4\|PC 16:0_20:4 B | 3.007 | 0.013 |
| PC 40:7\|PC 18:1_22:6 | 2.893 | 0.018 |
| PC 34:1\|PC 16:0_18:1 A | 2.834 | 0.022 |

| | | |
|---|---|---|
| Cer 13:0;O2/38:1 | 2.812 | 0.002 |
| SM 34:1;O2\|SM 18:1;O2/16:0 A | 2.785 | 0.009 |
| PC 38:6\|PC 16:0_22:6 A | 2.778 | 0.025 |
| NAGly 22:6;O(FA 22:4) | 2.613 | 0.021 |
| PC 35:2 | 2.243 | 0.002 |
| PC O-15:4_26:7 A | 2.185 | 0.006 |
| PC 40:6 A | 2.183 | 0.010 |
| PC 38:6\|PC 16:0_22:6 B | 2.177 | 0.001 |
| Cer 13:0;O2/32:2 | 2.161 | 0.048 |
| PC 38:6 | 2.15 | 0.031 |
| Cer 13:2;O2/38:6 | 2.147 | 0.003 |
| SE 28:2/18:1 | 2.133 | 0.002 |
| PC 36:2\|PC 18:0_18:2 | 2.09 | 0.012 |
| TG 18:1_20:2_22:4 | 2.073 | 0.001 |
| PC 34:2\|PC 16:0_18:2 A | 2.06 | 0.044 |
| Cer 12:1;O2/40:12;O2 | 2.059 | 0.004 |
| TG 52:2\|TG 16:0_18:1_18:1 | 2.056 | 0.004 |
| SL 13:1;O/32:4;O | 2.047 | 0.008 |
| NAGly 20:4;O(FA 28:6) | 2.042 | 0.026 |
| TG 8:0_9:0_38:5 | 2.009 | 0.000 |
| HexCer 17:0;O2/24:5;O | 1.994 | 0.011 |
| PC O-10:0_28:6 A | 1.99 | 0.013 |
| PC 32:0\|PC 16:0_16:0 B | 1.941 | 0.032 |
| PC 36:4\|PC 16:0_20:4 D | 1.931 | 0.009 |
| PC 40:6 B | 1.924 | 0.012 |
| PC 36:1\|PC 18:0_18:1 B | 1.898 | 0.003 |
| PC O-10:0_28:6 B | 1.881 | 0.007 |
| TG 54:3\|TG 18:0_18:1_18:2 | 1.852 | 0.020 |
| PC 38:4\|PC 18:0_20:4 A | 1.848 | 0.015 |
| TG 8:0_10:0_38:4 | 1.804 | 0.030 |
| PC 38:6 A | 1.802 | 0.012 |
| PC 36:3\|PC 16:0_20:3 A | 1.794 | 0.022 |
| SM 34:1;O2 A | 1.782 | 0.035 |
| TG 54:5\|TG 16:0_18:1_20:4 | 1.78 | 0.008 |
| Cer 13:1;O3/36:5(2OH) | 1.771 | 0.002 |
| TG 50:1\|TG 16:0_16:0_18:1 | 1.763 | 0.006 |
| SL 22:1;O/36:0;O | 1.761 | 0.004 |
| SM 42:2;O2\|SM 18:1;O2/24:1 | 1.76 | 0.008 |
| TG 8:0_9:0_22:2 | 1.751 | 0.039 |
| Hex2Cer 16:1;O2/18:3 | 1.737 | 0.002 |
| SM 12:1;O2/30:1 B | 1.722 | 0.015 |
| SL 20:1;O/36:0 | 1.721 | 0.003 |
| CE 18:3 | 1.721 | 0.006 |
| SM 42:3;O2 | 1.719 | 0.013 |
| BMP 13:1_28:7 | 1.717 | 0.001 |
| PC 40:6\|PC 18:0_22:6 | 1.7 | 0.010 |
| TG 56:6\|TG 18:1_18:2_20:3 | 1.694 | 0.002 |
| CE 22:6 | 1.687 | 0.007 |
| SL 22:1;O/36:2;O | 1.674 | 0.001 |

| | | |
|---|---|---|
| PC 36:3\|PC 18:1_18:2 | 1.669 | 0.005 |
| HexCer 17:0;O2/24:6;O A | 1.666 | 0.027 |
| TG 8:0_14:0_38:6 | 1.663 | 0.007 |
| PC 7:0_28:2 | 1.659 | 0.030 |
| CE 17:1 | 1.647 | 0.004 |
| TG 8:0_10:0_38:8 | 1.641 | 0.017 |
| CE 20:3 | 1.638 | 0.034 |
| TG 8:0_9:0_36:3 B | 1.617 | 0.032 |
| NAGlySer 10:0;O(FA 28:5) | 1.612 | 0.003 |
| SM 36:1;O2 B | 1.594 | 0.011 |
| SL 15:3;O/32:6 A | 1.593 | 0.012 |
| TG 54:4\|TG 18:1_18:1_18:2 A | 1.564 | 0.004 |
| CE 18:0 | 1.564 | 0.043 |
| PC 36:4 | 1.56 | 0.035 |
| TG 8:0_8:0_36:7 | 1.542 | 0.006 |
| PC 36:1 | 1.538 | 0.005 |
| SM 12:1;O2/36:0 | 1.537 | 0.006 |
| TG 8:0_8:0_38:7 | 1.515 | 0.029 |
| TG 8:0_9:0_34:2 | 1.503 | 0.001 |
| ST 27:2;O F | 1.503 | 0.004 |
| HexCer 16:0;O2/28:5;O | 1.501 | 0.043 |
| PC 36:3\|PC 16:0_20:3 B | 1.5 | 0.006 |
| ST 28:1;O B | 1.499 | 0.009 |
| SE 28:2/15:1 | 1.499 | 0.041 |
| NAGly 18:1 B | 1.497 | 0.005 |
| SM 12:1;O2/38:0 | 1.49 | 0.038 |
| TG 52:3\|TG 16:0_18:1_18:2 | 1.489 | 0.014 |
| TG 56:7\|TG 16:0_18:1_22:6 | 1.48 | 0.009 |
| SM 34:1;O2 B | 1.469 | 0.010 |
| PC 38:5\|PC 18:0_20:5 | 1.468 | 0.005 |
| ST 27:2;O E | 1.461 | 0.012 |
| TG 54:5\|TG 18:1_18:2_18:2 | 1.446 | 0.004 |
| CAR 16:0 B | 1.439 | 0.000 |
| Cer 12:1;O2/41:12;O2 A | 1.431 | 0.032 |
| HexCer 16:0;O2/18:1 | 1.428 | 0.019 |
| LPC 20:4 B | 1.421 | 0.001 |
| SL 22:1;O/36:3;O | 1.416 | 0.019 |
| TG 48:0\|TG 14:0_16:0_18:0 | 1.415 | 0.014 |
| LPC 18:1 C | 1.413 | 0.004 |
| TG 52:4\|TG 16:1_18:1_18:2 | 1.409 | 0.009 |
| BMP 8:0_24:1 | 1.407 | 0.000 |
| HexCer 17:0;O2/24:6;O B | 1.406 | 0.004 |
| ST 28:2;O | 1.394 | 0.001 |
| Cer 12:0;O2/22:0;O | 1.391 | 0.033 |
| CE 20:5 | 1.384 | 0.010 |
| SL 13:1;O/34:6 | 1.383 | 0.005 |
| SL 15:3;O/32:6 B | 1.382 | 0.009 |
| TG 20:0_18:1_18:1 | 1.381 | 0.008 |
| SE 28:2/21:3 | 1.371 | 0.006 |

| | | |
|---|---|---|
| HexCer 16:0;O2/19:0 A | 1.369 | 0.008 |
| Cer 41:0;O2\|Cer 18:0;O2/23:0 | 1.369 | 0.046 |
| ST 28:1;O A | 1.365 | 0.018 |
| SL 18:1;O/36:1;O | 1.361 | 0.009 |
| PC 36:2\|PC 18:1_18:1 | 1.358 | 0.008 |
| LPC 16:0 | 1.342 | 0.005 |
| TG 8:0_12:0_38:10 | 1.341 | 0.006 |
| TG 46:1\|TG 12:0_16:0_18:1 | 1.341 | 0.007 |
| SL 18:1;O/36:3;O | 1.337 | 0.007 |
| SL 20:1;O/36:0;O B | 1.33 | 0.006 |
| Cer 12:2;O2/39:11;O2 A | 1.327 | 0.007 |
| TG 50:3\|TG 16:0_16:1_18:2 | 1.325 | 0.013 |
| PC 34:1\|PC 16:0_18:1 B | 1.325 | 0.026 |
| TG 56:8\|TG 16:0_18:2_22:6 | 1.318 | 0.007 |
| Cer 46:4;O2\|Cer 30:3;O2/16:1 | 1.317 | 0.002 |
| TG 8:0_13:0_38:6 | 1.313 | 0.029 |
| CE 20:1 | 1.312 | 0.015 |
| TG 8:0_10:0_38:6 | 1.31 | 0.000 |
| LPC 14:0/0:0 | 1.305 | 0.017 |
| TG 8:0_8:0_32:6 B | 1.304 | 0.008 |
| Cer 34:1;O2\|Cer 18:1;O2/16:0 | 1.304 | 0.027 |
| PC 38:3 | 1.301 | 0.042 |
| SHexCer 12:1;O2/22:0;O B | 1.3 | 0.020 |
| Cer 13:2;O2/38:2 | 1.295 | 0.021 |
| CE 22:5 | 1.288 | 0.007 |
| TG 8:0_11:0_38:3 | 1.284 | 0.010 |
| Cer 12:0;O2/30:0;O | 1.283 | 0.030 |
| TG 50:2\|TG 16:0_16:1_18:1 | 1.273 | 0.017 |
| Cer 17:0;O2/25:0;O | 1.273 | 0.021 |
| TG 8:0_11:0_38:4 | 1.272 | 0.002 |
| TG 48:2\|TG 14:0_16:0_18:2 | 1.272 | 0.011 |
| ST 27:1;O B | 1.27 | 0.010 |
| CAR 26:7 | 1.264 | 0.004 |
| Cer 21:1;O3/38:1(2OH) B | 1.264 | 0.013 |
| SE 24:1;O4/26:0;O | 1.262 | 0.030 |
| SL 15:3;O/34:6 | 1.254 | 0.013 |
| TG 48:1\|TG 14:0_16:0_18:1 | 1.25 | 0.014 |
| TG 42:2\|TG 12:0_12:0_18:2 | 1.241 | 0.014 |
| TG 54:6\|TG 16:0_18:2_20:4 | 1.24 | 0.003 |
| PC 34:2\|PC 16:0_18:2 B | 1.239 | 0.028 |
| TG 45:2\|TG 12:0_18:0_15:2 | 1.234 | 0.022 |
| SL 13:1;O/32:6 | 1.229 | 0.031 |
| Cer 12:1;O2/39:11;O2 B | 1.227 | 0.010 |
| SM 12:1;O2/30:1 A | 1.223 | 0.036 |
| ST 29:1;O A | 1.222 | 0.015 |
| HexCer 41:1;O2\|HexCer 18:1;O2/23:0 | 1.217 | 0.000 |
| TG 16:0_20:0_18:1 | 1.217 | 0.009 |
| TG 48:4\|TG 12:0_18:2_18:2 | 1.216 | 0.012 |
| TG 52:0\|TG 16:0_18:0_18:0 | 1.216 | 0.022 |

| | | |
|---|---|---|
| TG 10:0_15:0_20:2 | 1.213 | 0.012 |
| TG 8:0_12:0_38:3 | 1.213 | 0.026 |
| LPC 18:0 | 1.212 | 0.006 |
| CAR 16:1 | 1.21 | 0.003 |
| HexCer 16:0;O3/18:0(2OH) | 1.21 | 0.003 |
| NAGly 16:2;O(FA 28:5) | 1.209 | 0.006 |
| TG 18:0_18:0_20:0 | 1.209 | 0.046 |
| HexCer 34:1;O3\|HexCer 18:1;O2/16:0;C | 1.206 | 0.035 |
| VAE 26:3 | 1.201 | 0.003 |
| Cer 12:0;O2/22:1;O | 1.198 | 0.023 |
| HexCer 18:1;O2/24:0 | 1.196 | 0.019 |
| ST 27:2;O D | 1.196 | 0.040 |
| TG 8:0_9:0_30:2 A | 1.19 | 0.002 |
| SL 12:0;O/33:0;O | 1.189 | 0.013 |
| TG 8:0_9:0_38:2 | 1.186 | 0.010 |
| TG 50:5\|TG 12:0_18:1_20:4 | 1.184 | 0.007 |
| PC 32:0\|PC 16:0_16:0 C | 1.184 | 0.029 |
| SL 16:1;O/36:3;O | 1.184 | 0.036 |
| AHexCer (O-12:0)12:1;O2/16:3;O | 1.18 | 0.007 |
| Cer 18:0;O2/24:0 | 1.18 | 0.018 |
| TG 56:7\|TG 18:1_18:2_20:4 | 1.178 | 0.007 |
| AHexCer (O-12:0)12:2;O2/12:0;O A | 1.177 | 0.011 |
| TG 8:0_8:0_32:6 A | 1.174 | 0.003 |
| TG 8:0_14:0_38:3 | 1.173 | 0.007 |
| PC O-8:0_26:1 | 1.17 | 0.002 |
| Cer 27:3;O2/11:0 | 1.168 | 0.017 |
| LPC 18:1 A | 1.167 | 0.006 |
| TG 50:6\|TG 12:0_18:2_20:4 | 1.167 | 0.009 |
| Cer 14:0;O2/38:3 | 1.167 | 0.013 |
| HexCer 17:0;O2/22:5;O | 1.167 | 0.024 |
| TG 44:0\|TG 12:0_14:0_18:0 | 1.166 | 0.009 |
| PC 38:5 B | 1.165 | 0.024 |
| NAGly 20:1 | 1.163 | 0.005 |
| Cer 41:1;O2\|Cer 18:1;O2/23:0 B | 1.163 | 0.007 |
| TG 8:0_9:0_34:4 | 1.162 | 0.016 |
| PC 38:4\|PC 18:0_20:4 C | 1.162 | 0.029 |
| SL 15:0;O/36:0;O | 1.159 | 0.033 |
| PC 38:3\|PC 18:0_20:3 | 1.157 | 0.004 |
| TG 50:4\|TG 14:0_18:1_18:3 | 1.157 | 0.011 |
| SM 12:1;O2/34:0 | 1.157 | 0.016 |
| TG 8:0_13:0_38:8 | 1.153 | 0.023 |
| LPC 18:2 A | 1.15 | 0.006 |
| HexCer 16:0;O2/25:1 | 1.143 | 0.018 |
| Cer 12:2;O2/36:10;O2 | 1.141 | 0.006 |
| TG 8:0_9:0_26:2 | 1.141 | 0.022 |
| SM 37:3;O3 | 1.14 | 0.029 |
| Cer 42:1;O3\|Cer 18:1;O2/24:0;O | 1.139 | 0.015 |
| NAGlySer 13:0;O(FA 28:5) | 1.139 | 0.048 |
| HexCer 18:1;O2/22:0 | 1.138 | 0.007 |

| | | |
|---|---|---|
| HexCer 16:0;O2/26:1 | 1.137 | 0.012 |
| DG 49:4 | 1.137 | 0.041 |
| TG 8:0_8:0_26:2 | 1.136 | 0.012 |
| NAGly 8:0;O(FA 22:5) | 1.136 | 0.030 |
| LPC 22:6 | 1.135 | 0.008 |
| TG 18:0_20:1_18:3 | 1.133 | 0.029 |
| TG 8:0_8:0_20:0 B | 1.129 | 0.018 |
| DG 49:9 | 1.128 | 0.042 |
| LPC 18:2 B | 1.127 | 0.041 |
| SL 13:1;O/32:5 | 1.124 | 0.010 |
| TG 40:1\|TG 10:0_12:0_18:1 | 1.122 | 0.037 |
| Cer 13:0;O2/34:3 | 1.121 | 0.018 |
| TG 54:1\|TG 18:0_18:0_18:1 | 1.12 | 0.005 |
| TG 36:0\|TG 10:0_12:0_14:0 | 1.116 | 0.002 |
| TG 42:1\|TG 12:0_12:0_18:1 | 1.116 | 0.007 |
| TG 10:0_10:0_24:1 | 1.107 | 0.020 |
| SE 24:1;O4/25:1;O A | 1.106 | 0.002 |
| TG 40:2\|TG 10:0_12:0_18:2 | 1.105 | 0.011 |
| Cer 42:1;O2\|Cer 18:1;O2/24:0 A | 1.102 | 0.026 |
| HexCer 34:1;O2\|HexCer 18:1;O2/16:0 | 1.1 | 0.008 |
| Cer 12:2;O2/38:11;O2 | 1.097 | 0.006 |
| TG 40:0\|TG 10:0_14:0_16:0 | 1.096 | 0.007 |
| Cer 47:2;O2\|Cer 20:1;O2/27:1 | 1.095 | 0.007 |
| Cer 42:2;O2\|Cer 18:1;O2/24:1 A | 1.093 | 0.006 |
| Cer 46:2;O2\|Cer 18:1;O2/28:1 | 1.093 | 0.039 |
| DG O-37:1\|DG O-23:0_14:1 | 1.089 | 0.000 |
| Cer 13:0;O2/24:2;O | 1.085 | 0.016 |
| SE 24:1;O4/27:0;O | 1.085 | 0.021 |
| HexCer 34:1;O3\|HexCer 18:1;O2/16:0;C | 1.083 | 0.012 |
| TG 44:2\|TG 12:0_14:0_18:2 | 1.081 | 0.020 |
| PC O-15:4_26:7 B | 1.081 | 0.029 |
| TG 8:0_26:1_24:3 | 1.08 | 0.002 |
| TG 16:0_18:0_18:2 | 1.078 | 0.020 |
| PC O-8:0_9:0 | 1.074 | 0.003 |
| TG 38:0\|TG 12:0_12:0_14:0 | 1.072 | 0.018 |
| TG 44:1\|TG 12:0_14:0_18:1 | 1.07 | 0.018 |
| TG 17:0_13:1_17:1 | 1.069 | 0.007 |
| TG 14:0_16:0_18:0 | 1.066 | 0.027 |
| TG 16:0_16:0_18:1 | 1.064 | 0.013 |
| TG 8:0_10:0_38:3 | 1.063 | 0.032 |
| TG 42:0\|TG 12:0_14:0_16:0 | 1.062 | 0.013 |
| TG 14:0_16:0_16:2 | 1.056 | 0.002 |
| Cer 12:0;O3/22:0(2OH) | 1.056 | 0.016 |
| LPC 18:1 B | 1.054 | 0.006 |
| TG 18:1_18:1_18:1 | 1.05 | 0.015 |
| TG 12:0_18:0_22:0 | 1.049 | 0.014 |
| PC 6:0_32:6 | 1.046 | 0.002 |
| NAGly 22:6;O(FA 26:6) | 1.04 | 0.016 |
| LPE 20:4 | 1.039 | 0.011 |

| | | |
|---|---|---|
| SL 12:1;O/24:1 B | 1.037 | 0.043 |
| SL 16:0;O/36:0;O | 1.036 | 0.031 |
| TG 11:0_11:0_21:0 | 1.035 | 0.016 |
| SE 28:2/19:5 | 1.035 | 0.025 |
| Cer 12:2;O2/15:3 | 1.034 | 0.002 |
| TG 8:0_8:0_38:1 | 1.032 | 0.000 |
| TG 18:0_18:0_22:1 | 1.029 | 0.011 |
| TG 13:0_13:0_18:0 | 1.026 | 0.011 |
| TG 16:0_18:1_18:1 | 1.025 | 0.042 |
| SE 24:1;O4/28:0;O A | 1.023 | 0.001 |
| TG 46:2\|TG 12:0_16:0_18:2 | 1.023 | 0.018 |
| SL 17:0;O/36:0;O A | 1.021 | 0.004 |
| LPC 20:4 A | 1.021 | 0.006 |
| TG 14:0_16:0_16:0 | 1.02 | 0.005 |
| TG 8:0_18:0_22:2 | 1.016 | 0.011 |
| TG 48:3\|TG 12:0_18:1_18:2 | 1.013 | 0.016 |
| Cer 42:1;O2\|Cer 18:1;O2/24:0 B | 1.011 | 0.015 |
| Cer 42:2;O2\|Cer 18:1;O2/24:1 B | 1.01 | 0.026 |
| PC 36:4\|PC 16:0_20:4 C | 1.007 | 0.019 |
| TG 8:0_8:0_25:0 | 1.006 | 0.013 |
| TG 8:0_9:0_34:3 A | 1.004 | 0.022 |
| TG 11:0_18:0_20:1 | 0.993 | 0.009 |
| NAGly 22:3 | 0.988 | 0.011 |
| SL 16:1;O/36:1;O B | 0.986 | 0.000 |
| TG 10:0_10:1_8:0;O2 | 0.983 | 0.009 |
| TG 54:6\|TG 18:1_18:2_18:3 | 0.981 | 0.015 |
| TG 18:0_14:1_20:2 | 0.98 | 0.009 |
| TG 8:0_8:0_37:1 | 0.975 | 0.006 |
| TG 8:0_9:0_38:0 | 0.974 | 0.001 |
| Cer 23:1;O3/38:2(2OH) | 0.972 | 0.013 |
| TG 8:0_8:0_22:0 C | 0.971 | 0.007 |
| HexCer 17:0;O2/26:3;O | 0.971 | 0.030 |
| MGDG 9:0_14:1 | 0.97 | 0.000 |
| TG 8:0_8:0_23:0 | 0.97 | 0.026 |
| TG 18:0_20:1_18:2 | 0.968 | 0.007 |
| DG 53:7 | 0.964 | 0.004 |
| SE 28:2/20:1 | 0.964 | 0.029 |
| TG 14:0_18:1_18:1 | 0.962 | 0.014 |
| SE 24:1;O4/25:1;O B | 0.96 | 0.005 |
| Cer 12:0;O3/36:0(2OH) | 0.96 | 0.005 |
| PC 36:1\|PC 18:0_18:1 A | 0.957 | 0.006 |
| TG 17:0_15:1_17:1 | 0.957 | 0.008 |
| Cer 12:2;O2/40:12;O2 | 0.957 | 0.024 |
| PC 38:5 A | 0.956 | 0.013 |
| TG 8:0_11:1_8:0;O2 | 0.952 | 0.007 |
| Cer 53:3;O2\|Cer 17:3;O2/36:0 | 0.943 | 0.012 |
| TG 50:6\|TG 12:0_16:0_22:6 | 0.943 | 0.014 |
| NATau 24:4;O | 0.942 | 0.005 |
| LPE 22:6 | 0.937 | 0.021 |

| | | |
|---|---|---|
| TG 12:0_16:0_18:3 | 0.936 | 0.014 |
| TG 54:3\|TG 16:0_16:0_22:3 | 0.936 | 0.027 |
| PC O-15:4_24:6 | 0.934 | 0.031 |
| Cer 25:1;O3/38:0(2OH) | 0.931 | 0.015 |
| TG 11:0_17:0_19:1 | 0.925 | 0.007 |
| CAR 27:0 | 0.921 | 0.002 |
| LPC 16:1 | 0.918 | 0.018 |
| TG 8:0_8:0_30:4 | 0.916 | 0.001 |
| SL 19:0;O/36:0;O | 0.915 | 0.026 |
| SE 28:2/18:0 A | 0.913 | 0.009 |
| Cer 13:1;O3/36:4(2OH) | 0.912 | 0.047 |
| VAE 27:0 | 0.91 | 0.017 |
| SM 34:0;O3 | 0.908 | 0.007 |
| NAE 7:0 | 0.908 | 0.017 |
| Cer 13:1;O3/26:2(2OH) | 0.908 | 0.025 |
| Cer 41:1;O2\|Cer 18:1;O2/23:0 A | 0.903 | 0.021 |
| SL 13:0;O/36:0;O | 0.89 | 0.005 |
| TG 10:0_12:0_24:1 | 0.888 | 0.005 |
| LPC O-19:3 | 0.888 | 0.008 |
| TG 11:0_15:0_19:1 | 0.887 | 0.020 |
| TG 12:0_16:0_20:4 | 0.886 | 0.011 |
| TG 43:2\|TG 12:0_16:0_15:2 | 0.883 | 0.008 |
| NAPhe 20:4;O | 0.88 | 0.026 |
| NAGly 18:1 A | 0.871 | 0.004 |
| TG 8:0_8:0_29:1 | 0.871 | 0.018 |
| SM 32:7;O2 | 0.866 | 0.000 |
| C17-Sphingosine | 0.863 | 0.044 |
| TG 54:4\|TG 18:1_18:1_18:2 B | 0.86 | 0.002 |
| TG 8:0_9:0_36:6 | 0.86 | 0.048 |
| NAOrn 19:0;O A | 0.855 | 0.016 |
| DG 43:8 | 0.848 | 0.045 |
| PC 36:5\|PC 16:0_20:5 B | 0.848 | 0.045 |
| TG 8:0_8:0_24:0 | 0.847 | 0.007 |
| ST 27:1;O E | 0.847 | 0.009 |
| TG 12:0_14:0_16:0;O | 0.846 | 0.025 |
| TG 14:0_19:2_19:2 | 0.843 | 0.019 |
| Cer 13:2;O2/28:6 | 0.835 | 0.001 |
| ST 27:2;O C | 0.835 | 0.022 |
| Cer 50:2;O2\|Cer 20:0;O2/30:2 | 0.831 | 0.017 |
| LPE 16:0 | 0.829 | 0.016 |
| SM 35:1;O3 | 0.827 | 0.002 |
| TG 8:0_8:0_30:1 B | 0.824 | 0.009 |
| TG 8:0_8:0_34:1 | 0.823 | 0.015 |
| HexCer 16:0;O2/22:0;O | 0.82 | 0.003 |
| Cer 25:1;O2/22:3;O | 0.818 | 0.027 |
| TG 8:0_28:1_19:2 | 0.818 | 0.029 |
| Cer 12:0;O2/29:0 | 0.815 | 0.009 |
| LPC 15:2 | 0.814 | 0.013 |
| TG 8:0_8:0_30:3 | 0.812 | 0.025 |

| | | |
|---|---|---|
| NATau 17:1;O A | 0.809 | 0.017 |
| LPE 18:0 | 0.809 | 0.036 |
| SE 28:2/24:4 | 0.807 | 0.004 |
| TG 14:0_16:0_17:0 | 0.801 | 0.021 |
| NAGlySer 9:0;O(FA 26:4) | 0.801 | 0.030 |
| TG 18:0_18:1_18:1 | 0.8 | 0.008 |
| ST 29:2;O | 0.793 | 0.017 |
| Cer 12:2;O2/41:12;O2 | 0.793 | 0.026 |
| SM 43:4;O2\|SM 17:3;O2/26:1 | 0.793 | 0.042 |
| DG 18:0_34:0 | 0.791 | 0.042 |
| Cer 42:1;O2\|Cer 18:1;O2/24:0 C | 0.79 | 0.033 |
| Cer 13:0;O2/36:5;O | 0.79 | 0.049 |
| Cer 14:2;O3/8:0(2OH) | 0.786 | 0.032 |
| TG 8:0_8:0_28:2 A | 0.786 | 0.045 |
| LPC 20:3/0:0 | 0.785 | 0.017 |
| TG 8:0_8:1_10:0;O2 | 0.785 | 0.031 |
| NAE 15:1 | 0.783 | 0.003 |
| SM 12:1;O2/28:0 | 0.783 | 0.008 |
| NATau 22:3;O | 0.782 | 0.029 |
| NATau 9:0 A | 0.781 | 0.004 |
| TG 8:0_11:0_38:6 | 0.776 | 0.004 |
| DG 55:8 | 0.776 | 0.023 |
| PE O-8:0_15:1 | 0.771 | 0.024 |
| Cer 40:0;O2\|Cer 18:0;O2/22:0 | 0.771 | 0.027 |
| SM 39:3;O3 | 0.767 | 0.035 |
| LPC 15:1 | 0.761 | 0.005 |
| Cer 12:0;O3/33:0(2OH) | 0.753 | 0.048 |
| TG 45:0\|TG 14:0_15:0_16:0 | 0.752 | 0.020 |
| TG 17:0_17:1_19:1 | 0.75 | 0.044 |
| TG 8:0_8:0_34:3 | 0.749 | 0.017 |
| TG 8:0_8:0_21:0 | 0.747 | 0.001 |
| NATau 20:0;O | 0.745 | 0.019 |
| NAPhe 17:4 | 0.744 | 0.005 |
| Cer 12:1;O3/25:1(2OH) | 0.742 | 0.011 |
| TG 8:0_8:0_28:2 C | 0.735 | 0.023 |
| PC O-33:6 | 0.734 | 0.017 |
| SE 28:1/19:0 | 0.731 | 0.006 |
| NAGlySer 11:0;O | 0.731 | 0.008 |
| Cer 12:1;O2/39:11;O2 C | 0.731 | 0.026 |
| TG 8:0_8:0_32:4 | 0.724 | 0.005 |
| TG 17:0_15:1_19:1 | 0.716 | 0.032 |
| DG 15:3_19:4 | 0.71 | 0.040 |
| MGDG O-26:6_7:0 | 0.709 | 0.034 |
| CE 21:3 | 0.707 | 0.002 |
| SM 34:1;O3 | 0.705 | 0.014 |
| CE 21:0 | 0.702 | 0.003 |
| PI 14:0 | 0.702 | 0.033 |
| TG 8:0_8:0_36:1 B | 0.702 | 0.036 |
| Cer 12:1;O3/24:5(2OH) B | 0.697 | 0.011 |

| | | |
|---|---|---|
| DG O-21:5_2:0 | 0.697 | 0.019 |
| TG 8:0_9:0_36:3 A | 0.695 | 0.037 |
| NAE 22:5 | 0.684 | 0.004 |
| TG 16:0_16:0_19:1 | 0.684 | 0.020 |
| TG 8:0_8:0_26:1 | 0.681 | 0.013 |
| SE 28:2/19:0 B | 0.68 | 0.002 |
| Cer 12:2;O2/19:2 | 0.677 | 0.030 |
| MGDG O-17:2_4:0 | 0.675 | 0.009 |
| Cer 40:1;O2\|Cer 18:1;O2/22:0 B | 0.675 | 0.046 |
| Cer 15:3;O2/18:5 | 0.674 | 0.002 |
| Cer 12:2;O2/39:11;O2 B | 0.674 | 0.002 |
| Cer 42:2;O2\|Cer 18:1;O2/24:1 C | 0.669 | 0.035 |
| VAE 21:2 B | 0.667 | 0.009 |
| Cer 15:3;O3/18:5(2OH) | 0.664 | 0.040 |
| ST 27:1;O I | 0.663 | 0.009 |
| DG O-15:1_7:0 | 0.662 | 0.016 |
| NAGly 15:3 A | 0.658 | 0.016 |
| NAGly 26:2 B | 0.654 | 0.003 |
| NAE 20:2 | 0.654 | 0.003 |
| TG 8:0_8:0_30:2 | 0.653 | 0.005 |
| TG 13:1_15:1_19:4 | 0.652 | 0.015 |
| DG O-20:5_4:0 | 0.646 | 0.002 |
| TG 8:0_8:0_24:2 C | 0.643 | 0.017 |
| Cer 17:0;O2/38:2 | 0.64 | 0.011 |
| DG 18:0_36:0 | 0.634 | 0.037 |
| PC 36:4\|PC 18:2_18:2 B | 0.627 | 0.036 |
| Cer 13:0;O2/22:6;O C | 0.622 | 0.029 |
| Cer 12:0;O2/24:1;O | 0.621 | 0.049 |
| TG 8:0_8:0_36:0 | 0.619 | 0.039 |
| Cer 12:0;O2/17:0;O A | 0.615 | 0.040 |
| DG 47:7 | 0.614 | 0.019 |
| SM 12:0;O2/21:0 | 0.611 | 0.002 |
| TG 8:0_8:0_28:1 | 0.61 | 0.021 |
| ST 27:1;O H | 0.61 | 0.024 |
| PE 20:1 | 0.609 | 0.020 |
| Cer 12:2;O2/19:4 | 0.605 | 0.014 |
| DG O-19:5_15:3 | 0.6 | 0.033 |
| DG 52:2 | 0.597 | 0.034 |
| DG O-22:5_3:0 | 0.593 | 0.004 |
| CAR 28:6 D | 0.593 | 0.006 |
| NATau 18:2 B | 0.587 | 0.021 |
| SL 20:3;O/18:3;O | 0.586 | 0.013 |

| **IgE > IgE + IgG** | | |
|---|---|---|
| ***Metabolite*** | ***Log2(Fold Change)*** | ***P-value*** |
| VAE 25:1 B | -1.711 | 0.000 |
| TG 9:0_9:0_20:1 | -1.604 | 0.000 |
| PS 12:0 | -1.425 | 0.005 |
| CE 18:2 | -1.26 | 0.018 |

| | | |
|---|---|---|
| VAE 25:1 A | -0.849 | 0.027 |
| SE 28:2/20:1 | -0.82 | 0.001 |
| ST 27:1;O G | -0.677 | 0.004 |

| **IgE + IgG > IgE** | | |
|---|---|---|
| ***Metabolite*** | ***Log2(Fold Change)*** | ***P-value*** |
| SM 34:1;O2\|SM 18:1;O2/16:0 B | 3.643 | 0.043 |
| NAGlySer 11:0;O(FA 28:5) | 2.943 | 0.002 |
| PC O-39:10 | 2.883 | 0.021 |
| NAGlySer 11:0;O(FA 28:6) | 2.81 | 0.016 |
| PC O-38:5\|PC O-20:5_18:0 | 2.799 | 0.021 |
| SHexCer 12:1;O2/22:0;O A | 2.239 | 0.017 |
| SM 38:4;O3 | 2.133 | 0.006 |
| PI 38:4 | 2.026 | 0.023 |
| PC 40:7\|PC 18:1_22:6 | 2.017 | 0.031 |
| HexCer 16:0;O2/24:3;O | 1.932 | 0.007 |
| Cer 12:0;O2/37:0;O | 1.808 | 0.014 |
| NATau 20:0;O | 1.776 | 0.001 |
| PI 36:1 | 1.711 | 0.018 |
| Cer 12:0;O2/38:0;O A | 1.68 | 0.007 |
| SL 12:1;O/35:1;O | 1.608 | 0.002 |
| PC 34:2\|PC 16:0_18:2 A | 1.597 | 0.041 |
| TG 8:0_9:0_28:2 | 1.557 | 0.009 |
| SM 34:1;O2\|SM 18:1;O2/16:0 A | 1.548 | 0.040 |
| TG O-18:0_16:1_16:1 | 1.54 | 0.026 |
| Cer 13:0;O2/36:4;O | 1.538 | 0.028 |
| CE 20:1 | 1.527 | 0.018 |
| NATau 22:3;O | 1.517 | 0.005 |
| Cer 13:2;O2/38:6 | 1.489 | 0.014 |
| BMP 13:1_28:7 | 1.479 | 0.021 |
| LPC 12:0 | 1.437 | 0.005 |
| SE 28:2/22:0 | 1.428 | 0.024 |
| CE 22:6 | 1.41 | 0.017 |
| ST 27:1;O C | 1.406 | 0.009 |
| ST 28:1;O B | 1.403 | 0.015 |
| TG 48:0\|TG 14:0_16:0_18:0 | 1.387 | 0.026 |
| PC 36:5\|PC 16:0_20:5 A | 1.367 | 0.017 |
| SE 28:2/21:4 | 1.343 | 0.020 |
| HexCer 16:0;O2/19:0 A | 1.34 | 0.036 |
| NAGly 8:0;O(FA 22:5) | 1.326 | 0.014 |
| PC O-10:0_28:6 B | 1.323 | 0.042 |
| Cer 17:1;O3/38:2(2OH) | 1.317 | 0.002 |
| CE 20:5 | 1.31 | 0.015 |
| CE 18:3 | 1.275 | 0.024 |
| TG 44:0\|TG 12:0_14:0_18:0 | 1.273 | 0.012 |
| TG 14:0_16:0_18:0 | 1.272 | 0.009 |
| Cer 12:1;O2/39:11;O2 A | 1.269 | 0.021 |
| CE 20:4 | 1.257 | 0.009 |
| TG 46:0\|TG 12:0_16:0_18:0 | 1.254 | 0.014 |

| | | |
|---|---|---|
| PC 40:6\|PC 18:0_22:6 | 1.249 | 0.038 |
| TG 12:0_18:0_22:0 | 1.244 | 0.019 |
| ST 28:1;O A | 1.237 | 0.011 |
| TG 18:0_18:0_20:0 | 1.237 | 0.044 |
| SE 28:2/15:1 | 1.234 | 0.037 |
| TG O-54:2\|TG O-18:0_16:1_20:1 | 1.225 | 0.017 |
| PC 36:4\|PC 16:0_20:4 D | 1.217 | 0.031 |
| VAE 27:0 | 1.205 | 0.016 |
| SM 12:1;O2/28:0 | 1.201 | 0.023 |
| TG 13:0_18:0_20:0 | 1.199 | 0.009 |
| Cer 13:0;O2/32:2 | 1.189 | 0.000 |
| PC 38:6 A | 1.177 | 0.027 |
| ST 27:1;O B | 1.17 | 0.013 |
| CE 17:1 | 1.169 | 0.041 |
| ST 27:2;O E | 1.162 | 0.034 |
| SL 18:1;O/36:1;O | 1.152 | 0.001 |
| Cer 12:0;O2/28:1;O | 1.148 | 0.028 |
| TG 42:0\|TG 12:0_14:0_16:0 | 1.138 | 0.017 |
| TG 8:0_8:0_24:2 B | 1.137 | 0.004 |
| Cer 12:0;O2/38:0;O B | 1.135 | 0.024 |
| SE 24:1;O4/25:1;O B | 1.13 | 0.014 |
| TG 12:0_16:0_20:4 | 1.13 | 0.034 |
| Cer 15:3;O2/36:6 | 1.129 | 0.022 |
| SL 20:1;O/36:0 | 1.127 | 0.036 |
| TG 8:0_14:0_38:5 | 1.126 | 0.045 |
| Cer 13:1;O3/38:0(2OH) | 1.118 | 0.002 |
| DG 18:0_28:0 | 1.117 | 0.044 |
| SE 28:2/18:0 B | 1.112 | 0.003 |
| VAE 25:0 A | 1.098 | 0.020 |
| AHexCer (O-12:0)12:2;O2/12:0;O A | 1.098 | 0.033 |
| SL 22:1;O/36:0;O | 1.095 | 0.046 |
| Cer 19:1;O3/38:1(2OH) | 1.091 | 0.002 |
| SE 28:2/18:1 | 1.086 | 0.033 |
| TG 14:0_19:2_19:2 | 1.082 | 0.003 |
| TG 13:1_15:1_19:4 | 1.08 | 0.008 |
| SE 24:1;O4/25:1;O A | 1.078 | 0.024 |
| TG 8:0_8:0_28:2 A | 1.078 | 0.039 |
| TG 8:0_8:0_26:1 | 1.077 | 0.035 |
| Cer 21:1;O3/38:1(2OH) B | 1.072 | 0.020 |
| TG 8:0_8:0_26:0 | 1.07 | 0.022 |
| TG 14:0_18:1_18:1 | 1.069 | 0.016 |
| TG 17:0_13:1_17:1 | 1.065 | 0.023 |
| TG 40:0\|TG 10:0_14:0_16:0 | 1.059 | 0.026 |
| SL 13:2;O/36:6 | 1.054 | 0.001 |
| TG 48:1\|TG 14:0_16:0_18:1 | 1.03 | 0.036 |
| TG 11:0_15:0_19:1 | 1.024 | 0.025 |
| SL 15:3;O/34:6 | 1.024 | 0.037 |
| Cer 12:0;O2/26:0;O | 1.022 | 0.018 |
| AHexCer (O-12:0)12:1;O2/16:3;O | 1.02 | 0.012 |

| | | |
|---|---|---|
| SL 12:0;O/36:0;O | 1.017 | 0.014 |
| TG O-16:1_18:0_20:0 | 1.013 | 0.040 |
| SE 28:2/21:2 | 1.013 | 0.040 |
| TG 12:0_16:0_18:3 | 1.011 | 0.006 |
| SL 12:1;O/36:0;O | 1.009 | 0.004 |
| TG 45:0\|TG 14:0_15:0_16:0 | 1.009 | 0.018 |
| VAE 21:0 | 1.008 | 0.009 |
| TG 52:0\|TG 16:0_18:0_18:0 | 1.006 | 0.050 |
| TG 16:0_16:0_18:1 | 1.005 | 0.010 |
| TG 46:1\|TG 12:0_16:0_18:1 | 1.002 | 0.017 |
| TG 8:0_8:0_38:0 B | 0.999 | 0.005 |
| HexCer 16:0;O2/19:5 | 0.998 | 0.023 |
| TG 40:2\|TG 10:0_12:0_18:2 | 0.996 | 0.015 |
| TG 48:2\|TG 14:0_16:0_18:2 | 0.994 | 0.030 |
| Cer 46:2;O2\|Cer 18:1;O2/28:1 | 0.993 | 0.021 |
| PC O-15:4_24:6 | 0.99 | 0.003 |
| TG 16:0_18:1_18:1 | 0.986 | 0.015 |
| TG 14:0_16:0_17:0 | 0.976 | 0.019 |
| TG 8:0_8:0_32:6 A | 0.974 | 0.019 |
| Cer 40:1;O2\|Cer 18:1;O2/22:0 A | 0.973 | 0.026 |
| TG 44:1\|TG 12:0_14:0_18:1 | 0.971 | 0.023 |
| TG 50:1\|TG 16:0_16:0_18:1 | 0.968 | 0.030 |
| SE 28:1/19:0 | 0.967 | 0.032 |
| Cer 12:0;O2/32:0;O | 0.965 | 0.007 |
| SE 28:2/16:1 A | 0.96 | 0.039 |
| TG O-42:0\|TG O-16:0_12:0_14:0 | 0.957 | 0.038 |
| Cer 12:0;O2/28:0 | 0.955 | 0.005 |
| TG 13:0_13:0_18:0 | 0.954 | 0.032 |
| DG 49:9 | 0.954 | 0.045 |
| CAR 20:3 C | 0.952 | 0.022 |
| TG 8:0_8:0_28:2 B | 0.951 | 0.032 |
| TG 54:1\|TG 18:0_18:0_18:1 | 0.95 | 0.023 |
| PC 7:0_28:2 | 0.948 | 0.046 |
| Cer 12:1;O2/22:6 | 0.939 | 0.016 |
| TG 46:2\|TG 12:0_16:0_18:2 | 0.939 | 0.030 |
| LPC 20:3/0:0 | 0.937 | 0.039 |
| HexCer 16:0;O2/26:1 | 0.933 | 0.004 |
| TG 44:2\|TG 12:0_14:0_18:2 | 0.929 | 0.009 |
| Cer 50:2;O2\|Cer 20:0;O2/30:2 | 0.928 | 0.022 |
| TG 8:0_8:0_29:1 | 0.919 | 0.006 |
| TG 14:0_16:0_20:0 | 0.919 | 0.007 |
| Cer 13:0;O2/32:4;O | 0.916 | 0.048 |
| SM 12:1;O2/30:1 A | 0.909 | 0.020 |
| Cer 12:0;O2/34:0;O | 0.906 | 0.032 |
| TG 8:0_8:0_36:0 | 0.903 | 0.017 |
| Cer 42:3;O3\|Cer 18:1;O2/24:2;O | 0.896 | 0.024 |
| TG 8:0_8:0_30:0 | 0.895 | 0.007 |
| TG 50:4\|TG 16:0_16:2_18:2 | 0.893 | 0.024 |
| HexCer 16:0;O2/25:1 | 0.892 | 0.042 |

| | | |
|---|---|---|
| NAGly 20:0;O(FA 28:1) | 0.886 | 0.013 |
| NAGlySer 13:1;O(FA 26:6) | 0.886 | 0.038 |
| PC 36:1 | 0.884 | 0.022 |
| DG 49:4 | 0.882 | 0.004 |
| PC 6:0_30:1 | 0.873 | 0.024 |
| TG 8:0_18:0_22:2 | 0.871 | 0.035 |
| Cer 42:1;O2\|Cer 18:1;O2/24:0 A | 0.871 | 0.037 |
| TG 43:2\|TG 12:0_16:0_15:2 | 0.869 | 0.017 |
| Cer 40:1;O2\|Cer 18:1;O2/22:0 B | 0.863 | 0.043 |
| TG 17:0_15:1_17:1 | 0.858 | 0.018 |
| TG 38:0\|TG 12:0_12:0_14:0 | 0.858 | 0.048 |
| Carnosine | 0.846 | 0.011 |
| ST 27:2;O C | 0.843 | 0.025 |
| ST 29:2;O | 0.835 | 0.041 |
| Cer 12:1;O3/24:5(2OH) B | 0.833 | 0.015 |
| SM 34:0;O3 | 0.832 | 0.004 |
| TG 8:0_9:0_22:2 | 0.83 | 0.029 |
| TG 10:0_12:0_24:1 | 0.829 | 0.043 |
| TG O-52:1\|TG O-18:0_16:0_18:1 | 0.827 | 0.039 |
| NAGly 26:0;O(FA 28:2) | 0.824 | 0.041 |
| DG 47:7 | 0.82 | 0.005 |
| TG 12:0_14:0_16:0;O | 0.819 | 0.017 |
| PC 38:4\|PC 18:0_20:4 C | 0.811 | 0.037 |
| TG O-46:0\|TG O-18:0_12:0_16:0 | 0.811 | 0.037 |
| TG 8:0_8:0_32:0 | 0.81 | 0.020 |
| TG 8:0_8:0_28:0 | 0.81 | 0.032 |
| SL 13:1;O/36:3;O | 0.808 | 0.027 |
| PC 36:4\|PC 16:0_20:4 C | 0.806 | 0.046 |
| TG 13:0_18:0_18:0 | 0.799 | 0.009 |
| SL 20:1;O/36:0;O A | 0.797 | 0.020 |
| PC 36:1\|PC 18:0_18:1 A | 0.791 | 0.028 |
| Cer 12:0;O2/29:1 | 0.784 | 0.046 |
| TG 8:0_8:0_30:2 | 0.78 | 0.019 |
| Cer 13:2;O2/38:2 | 0.779 | 0.049 |
| SM 37:3;O3 | 0.776 | 0.008 |
| SL 15:3;O/32:6 B | 0.768 | 0.043 |
| SL 16:0;O/36:0;O | 0.766 | 0.031 |
| NAGly 13:0;O(FA 9:0) | 0.762 | 0.012 |
| VAE 26:3 | 0.754 | 0.022 |
| Cer 42:2;O2\|Cer 18:1;O2/24:1 B | 0.753 | 0.030 |
| TG 16:0_20:0_18:1 | 0.749 | 0.023 |
| HexCer 16:0;O2/26:4;O | 0.747 | 0.027 |
| SM 35:1;O3 | 0.746 | 0.026 |
| TG 11:0_17:0_19:1 | 0.745 | 0.026 |
| Cer 53:3;O2\|Cer 17:3;O2/36:0 | 0.743 | 0.036 |
| NAOrn 27:1 B | 0.741 | 0.005 |
| Cer 47:2;O2\|Cer 20:1;O2/27:1 | 0.721 | 0.035 |
| TG 12:0_18:0_18:2 | 0.72 | 0.008 |
| DG 15:3_19:4 | 0.719 | 0.025 |

| | | |
|---|---|---|
| SM 34:1;O3 | 0.716 | 0.015 |
| TG 8:0_8:0_22:0 B | 0.71 | 0.004 |
| DG 49:6 | 0.709 | 0.035 |
| Cer 12:2;O2/15:3 | 0.699 | 0.027 |
| Cer 12:1;O3/32:1(2OH) | 0.699 | 0.028 |
| TG 50:4\|TG 14:0_18:2_18:2 | 0.698 | 0.001 |
| HexCer 34:1;O3\|HexCer 18:1;O2/16:0;C | 0.695 | 0.025 |
| PS 21:3 | 0.693 | 0.015 |
| NAE 20:2 | 0.692 | 0.000 |
| NAGly 20:1 | 0.686 | 0.044 |
| CE 21:3 | 0.682 | 0.029 |
| SE 28:2/22:2 | 0.68 | 0.043 |
| TG 8:0_8:0_30:3 | 0.679 | 0.050 |
| DG 50:1 | 0.673 | 0.010 |
| SM 32:0;O3 | 0.668 | 0.015 |
| SM 12:1;O2/23:0 | 0.659 | 0.028 |
| TG 10:0_10:0_24:1 | 0.657 | 0.011 |
| LPC O-15:3 C | 0.657 | 0.015 |
| TG 8:0_8:0_28:1 | 0.655 | 0.012 |
| ST 29:1;O A | 0.652 | 0.049 |
| SM 39:3;O3 | 0.649 | 0.048 |
| SE 28:2/24:4 | 0.646 | 0.035 |
| TG 8:0_8:0_30:4 | 0.645 | 0.022 |
| PC 6:0_32:6 | 0.641 | 0.036 |
| TG 17:0_17:1_19:1 | 0.639 | 0.049 |
| TG 50:6\|TG 12:0_16:0_22:6 | 0.638 | 0.010 |
| PC 38:6 B | 0.638 | 0.026 |
| HexCer 18:1;O2/24:0 | 0.632 | 0.032 |
| Cer 12:0;O2/36:0;O | 0.63 | 0.012 |
| Cer 12:2;O2/38:11;O2 | 0.623 | 0.003 |
| DG 46:0 | 0.623 | 0.046 |
| TG 16:0_16:0_19:1 | 0.619 | 0.043 |
| TG 8:0_8:0_37:1 | 0.615 | 0.038 |
| SM 41:3;O3 | 0.612 | 0.005 |
| DG 40:0 | 0.609 | 0.019 |
| Cer 25:1;O3/38:0(2OH) | 0.609 | 0.044 |
| SM 37:6;O2 | 0.607 | 0.002 |
| Cer 41:1;O2\|Cer 18:1;O2/23:0 B | 0.606 | 0.031 |
| DG 36:2\|DG 18:1_18:1 | 0.601 | 0.018 |
| LPE 16:0 | 0.599 | 0.033 |
| TG 8:0_8:0_36:1 B | 0.593 | 0.033 |
| TG 8:0_8:0_32:6 B | 0.588 | 0.042 |

| **IgE > IgG** | | |
|---|---|---|
| ***Metabolite*** | ***Log2(Fold Change)*** | ***P-value*** |
| PS 12:0 | -1.591 | 0.002 |
| VAE 25:1 B | -1.458 | 0.008 |
| TG 9:0_9:0_20:1 | -1.176 | 0.017 |
| ST 27:1;O G | -0.969 | 0.036 |

| | | |
|---|---|---|
| NATau 16:4 A | -0.898 | 0.024 |
| NAGlySer 18:2;O H | -0.818 | 0.007 |
| NAGly 28:6 E | -0.731 | 0.013 |
| SE 24:1;O4/6:0;O | -0.657 | 0.005 |
| NAOrn 24:5 D | -0.624 | 0.001 |

| **IgG > IgE** | | |
|---|---|---|
| ***Metabolite*** | ***Log2(Fold Change)*** | ***P-value*** |
| SM 34:1;O2\|SM 18:1;O2/16:0 B | 6.521 | 0.002 |
| PC 36:4\|PC 16:0_20:4 E | 4.854 | 0.001 |
| PC 38:6\|PC 16:0_22:6 C | 4.754 | 0.001 |
| PC 32:0\|PC 16:0_16:0 A | 4.692 | 0.043 |
| SE 24:1;O4/19:1;O | 4.677 | 0.000 |
| PC O-38:5\|PC O-18:1_20:4 | 4.573 | 0.003 |
| SM 38:4;O3 | 4.567 | 0.001 |
| PC O-38:5\|PC O-20:5_18:0 | 4.523 | 0.001 |
| PC 38:4\|PC 18:0_20:4 B | 4.477 | 0.000 |
| PC O-36:4\|PC O-16:0_20:4 | 4.445 | 0.003 |
| PC O-39:10 | 4.412 | 0.007 |
| SM 36:1;O2 C | 4.202 | 0.002 |
| SM 36:1;O2 A | 3.941 | 0.021 |
| Cer 13:0;O2/38:1 | 3.782 | 0.025 |
| PC 36:4\|PC 16:0_20:4 B | 3.764 | 0.000 |
| PC 34:1\|PC 16:0_18:1 A | 3.734 | 0.001 |
| PC 40:7\|PC 18:1_22:6 | 3.624 | 0.006 |
| PC 36:0 | 3.37 | 0.000 |
| SHexCer 12:1;O2/22:0;O A | 3.318 | 0.001 |
| Hex2Cer 16:0;O2/14:0 | 3.217 | 0.004 |
| SL 13:1;O/32:4;O | 3.187 | 0.037 |
| SM 34:1;O2\|SM 18:1;O2/16:0 A | 3.138 | 0.004 |
| PC 38:6\|PC 16:0_22:6 A | 3.137 | 0.004 |
| PC 38:4\|PC 18:0_20:4 A | 3.062 | 0.002 |
| PC 34:2\|PC 16:0_18:2 A | 2.955 | 0.003 |
| Cer 12:1;O2/40:12;O2 | 2.937 | 0.001 |
| NAGly 22:6;O(FA 22:4) | 2.817 | 0.000 |
| HexCer 17:0;O2/24:5;O | 2.793 | 0.001 |
| Cer 13:2;O2/38:6 | 2.764 | 0.002 |
| PC 38:6 | 2.762 | 0.005 |
| PC O-10:0_28:6 B | 2.746 | 0.003 |
| CE 18:0 | 2.718 | 0.032 |
| PC 36:4\|PC 18:2_18:2 A | 2.717 | 0.000 |
| TG 48:0\|TG 14:0_16:0_18:0 | 2.705 | 0.001 |
| SM 24:2;O2(FA 14:1) | 2.695 | 0.030 |
| PC 40:6 B | 2.666 | 0.005 |
| CE 20:1 | 2.625 | 0.003 |
| CE 18:3 | 2.618 | 0.001 |
| CE 17:1 | 2.616 | 0.006 |
| NAGlySer 11:0;O(FA 28:5) | 2.608 | 0.001 |
| PC 40:6 A | 2.602 | 0.013 |

| | | |
|---|---|---|
| LPC 12:0 | 2.598 | 0.000 |
| PC 35:2 | 2.593 | 0.002 |
| SL 12:1;O/35:1;O | 2.589 | 0.000 |
| PC 38:6\|PC 16:0_22:6 B | 2.573 | 0.004 |
| TG 44:0\|TG 12:0_14:0_18:0 | 2.553 | 0.000 |
| PC 36:4\|PC 16:0_20:4 D | 2.539 | 0.001 |
| TG 50:1\|TG 16:0_16:0_18:1 | 2.458 | 0.001 |
| Hex2Cer 16:1;O2/18:3 | 2.444 | 0.008 |
| PC O-15:4_26:7 A | 2.438 | 0.001 |
| PC 36:3\|PC 16:0_20:3 A | 2.42 | 0.002 |
| ST 28:1;O B | 2.412 | 0.000 |
| ST 27:2;O E | 2.401 | 0.001 |
| SL 20:1;O/36:0 | 2.401 | 0.004 |
| SE 28:2/18:1 | 2.397 | 0.003 |
| TG 18:0_18:0_20:0 | 2.394 | 0.001 |
| HexCer 16:0;O2/28:5;O | 2.392 | 0.028 |
| CE 22:6 | 2.373 | 0.001 |
| Cer 21:1;O3/38:2(2OH) | 2.37 | 0.002 |
| Cer 12:1;O2/39:11;O2 A | 2.366 | 0.000 |
| TG 14:0_16:0_18:0 | 2.363 | 0.000 |
| TG 42:0\|TG 12:0_14:0_16:0 | 2.311 | 0.001 |
| BMP 13:1_28:7 | 2.303 | 0.001 |
| TG 46:1\|TG 12:0_16:0_18:1 | 2.3 | 0.000 |
| TG 40:0\|TG 10:0_14:0_16:0 | 2.297 | 0.001 |
| ST 28:1;O A | 2.297 | 0.003 |
| PC 38:6 A | 2.292 | 0.001 |
| Cer 19:1;O3/38:1(2OH) | 2.291 | 0.000 |
| Cer 13:1;O3/36:4(2OH) | 2.285 | 0.008 |
| SE 28:2/18:0 B | 2.279 | 0.014 |
| NAGlySer 11:0;O(FA 28:6) | 2.273 | 0.004 |
| HexCer 16:0;O2/19:0 A | 2.261 | 0.003 |
| TG 52:0\|TG 16:0_18:0_18:0 | 2.259 | 0.003 |
| Cer 46:2;O2\|Cer 18:1;O2/28:1 | 2.241 | 0.001 |
| SE 28:2/15:1 | 2.225 | 0.013 |
| TG 48:2\|TG 14:0_16:0_18:2 | 2.224 | 0.001 |
| DG 18:0_34:0 | 2.22 | 0.006 |
| TG 10:0_15:0_20:2 | 2.206 | 0.009 |
| PC 32:0\|PC 16:0_16:0 B | 2.201 | 0.001 |
| TG 40:1\|TG 10:0_12:0_18:1 | 2.196 | 0.001 |
| TG 38:0\|TG 12:0_12:0_14:0 | 2.193 | 0.001 |
| PC 36:5\|PC 16:0_20:5 A | 2.193 | 0.001 |
| TG 40:2\|TG 10:0_12:0_18:2 | 2.191 | 0.000 |
| TG 12:0_18:0_22:0 | 2.185 | 0.003 |
| TG 8:0_9:0_34:4 | 2.182 | 0.000 |
| PC 36:2\|PC 18:0_18:2 | 2.175 | 0.005 |
| TG 48:1\|TG 14:0_16:0_18:1 | 2.173 | 0.003 |
| TG 13:0_13:0_18:0 | 2.169 | 0.000 |
| TG 8:0_9:0_36:6 | 2.167 | 0.018 |
| SM 34:1;O2 A | 2.166 | 0.001 |

| | | |
|---|---|---|
| SL 18:1;O/36:1;O | 2.165 | 0.006 |
| TG 8:0_9:0_26:2 | 2.158 | 0.004 |
| SE 28:2/21:3 | 2.15 | 0.024 |
| TG 8:0_8:0_26:2 | 2.146 | 0.003 |
| TG O-54:2\|TG O-18:0_16:1_20:1 | 2.144 | 0.003 |
| SL 22:1;O/36:0;O | 2.143 | 0.009 |
| TG 44:1\|TG 12:0_14:0_18:1 | 2.136 | 0.000 |
| TG 8:0_8:0_28:2 B | 2.136 | 0.001 |
| Cer 12:0;O2/37:0;O | 2.134 | 0.005 |
| CE 20:5 | 2.13 | 0.002 |
| TG 8:0_8:0_28:2 A | 2.129 | 0.004 |
| TG 42:1\|TG 12:0_12:0_18:1 | 2.116 | 0.001 |
| AHexCer (O-12:0)12:2;O2/12:0;O A | 2.116 | 0.003 |
| PC 36:1\|PC 18:0_18:1 B | 2.109 | 0.002 |
| TG 14:0_16:0_16:0 | 2.106 | 0.000 |
| SE 24:1;O4/26:1;O | 2.105 | 0.002 |
| Cer 47:2;O2\|Cer 20:1;O2/27:1 | 2.104 | 0.002 |
| HexCer 17:0;O2/24:6;O A | 2.103 | 0.000 |
| Cer 13:0;O2/36:4;O | 2.102 | 0.008 |
| PC 40:6\|PC 18:0_22:6 | 2.101 | 0.004 |
| Cer 12:1;O3/21:1(2OH) | 2.097 | 0.000 |
| NAGlySer 13:1;O(FA 26:6) | 2.094 | 0.009 |
| NAGly 20:4;O(FA 28:6) | 2.091 | 0.004 |
| TG 44:2\|TG 12:0_14:0_18:2 | 2.09 | 0.001 |
| TG 42:2\|TG 12:0_12:0_18:2 | 2.083 | 0.001 |
| Cer 13:0;O2/32:2 | 2.078 | 0.010 |
| TG 46:2\|TG 12:0_16:0_18:2 | 2.073 | 0.001 |
| TG 14:0_16:0_16:2 | 2.072 | 0.000 |
| Cer 12:0;O2/38:0;O A | 2.062 | 0.001 |
| Cer 12:0;O2/24:1;O | 2.058 | 0.015 |
| SM 36:1;O2 B | 2.055 | 0.002 |
| HexCer 16:0;O2/24:3;O | 2.051 | 0.002 |
| PC 36:4 | 2.048 | 0.001 |
| TG 16:0_16:0_18:1 | 2.047 | 0.000 |
| SM 42:3;O2 | 2.041 | 0.002 |
| SE 24:1;O4/25:1;O B | 2.038 | 0.001 |
| ST 27:1;O C | 2.027 | 0.009 |
| ST 29:1;O B | 2.026 | 0.014 |
| TG 54:1\|TG 18:0_18:0_18:1 | 2.025 | 0.002 |
| TG 14:0_16:0_17:0 | 2.023 | 0.002 |
| TG 14:0_18:1_18:1 | 2.018 | 0.000 |
| TG O-52:1\|TG O-18:0_16:0_18:1 | 2.018 | 0.001 |
| TG 14:0_19:2_19:2 | 2.005 | 0.001 |
| PC 36:3\|PC 18:1_18:2 | 2.005 | 0.003 |
| SL 15:3;O/32:6 A | 2.005 | 0.004 |
| TG 48:4\|TG 12:0_18:2_18:2 | 2 | 0.002 |
| TG 11:0_11:0_21:0 | 1.991 | 0.000 |
| Cer 21:0;O3/38:0(2OH) | 1.986 | 0.010 |
| SM 42:2;O2\|SM 18:1;O2/24:1 | 1.984 | 0.006 |

| | | |
|---|---|---|
| DG 18:0_36:0 | 1.981 | 0.000 |
| TG 36:0\|TG 10:0_12:0_14:0 | 1.976 | 0.000 |
| TG 10:0_12:0_24:1 | 1.971 | 0.000 |
| NAGly 26:0;O(FA 28:2) | 1.97 | 0.005 |
| TG 11:0_15:0_19:1 | 1.965 | 0.003 |
| PC 7:0_28:2 | 1.952 | 0.015 |
| ST 27:1;O B | 1.95 | 0.000 |
| NAGly 8:0;O(FA 22:5) | 1.944 | 0.008 |
| SM 37:3;O3 | 1.943 | 0.001 |
| TG 16:0_18:1_18:1 | 1.938 | 0.000 |
| TG 50:4\|TG 14:0_18:1_18:3 | 1.938 | 0.001 |
| HexCer 17:0;O2/24:6;O B | 1.935 | 0.001 |
| SL 17:0;O/36:0;O A | 1.932 | 0.008 |
| TG 50:4\|TG 16:0_16:2_18:2 | 1.929 | 0.003 |
| TG 50:2\|TG 16:0_16:1_18:1 | 1.927 | 0.002 |
| Cer 23:1;O3/38:2(2OH) | 1.923 | 0.030 |
| Cer 25:1;O2/22:3;O | 1.918 | 0.007 |
| TG 8:0_18:0_22:2 | 1.916 | 0.002 |
| TG 12:0_16:0_20:4 | 1.911 | 0.006 |
| Cer 13:2;O2/38:2 | 1.908 | 0.000 |
| SL 12:0;O/33:0;O | 1.9 | 0.009 |
| TG 17:0_13:1_17:1 | 1.898 | 0.001 |
| VAE 27:0 | 1.897 | 0.001 |
| SL 16:0;O/36:0;O | 1.897 | 0.002 |
| TG 13:0_18:0_20:0 | 1.893 | 0.000 |
| TG 54:4\|TG 18:1_18:1_18:2 A | 1.887 | 0.004 |
| TG 54:5\|TG 16:0_18:1_20:4 | 1.886 | 0.002 |
| CE 22:5 | 1.883 | 0.001 |
| TG 12:0_16:0_18:3 | 1.881 | 0.001 |
| TG 52:2\|TG 16:0_18:1_18:1 | 1.88 | 0.003 |
| PC O-10:0_28:6 A | 1.877 | 0.005 |
| DG 43:8 | 1.873 | 0.005 |
| Cer 13:1;O3/36:5(2OH) | 1.87 | 0.001 |
| NAGlySer 10:0;O(FA 28:5) | 1.87 | 0.006 |
| Cer 50:2;O2\|Cer 20:0;O2/30:2 | 1.868 | 0.000 |
| TG 54:5\|TG 18:1_18:2_18:2 | 1.868 | 0.002 |
| TG O-17:0_18:0_17:1 | 1.865 | 0.004 |
| TG 8:0_8:0_26:1 | 1.864 | 0.002 |
| TG 8:0_8:0_24:0 | 1.851 | 0.004 |
| PC 38:3 | 1.85 | 0.004 |
| Cer 12:1;O2/41:12;O2 A | 1.849 | 0.008 |
| TG O-16:1_18:0_20:0 | 1.845 | 0.003 |
| TG 18:0_18:0_22:1 | 1.842 | 0.004 |
| TG 52:4\|TG 16:1_18:1_18:2 | 1.838 | 0.001 |
| TG 8:0_8:0_32:6 B | 1.824 | 0.002 |
| TG 46:0\|TG 12:0_16:0_18:0 | 1.823 | 0.001 |
| TG 8:0_8:0_32:6 A | 1.823 | 0.001 |
| PC 36:3\|PC 16:0_20:3 B | 1.811 | 0.005 |
| TG O-42:0\|TG O-16:0_12:0_14:0 | 1.803 | 0.003 |

| | | |
|---|---|---|
| TG O-16:0_16:0_18:2 | 1.802 | 0.001 |
| Cer 53:3;O2\|Cer 17:3;O2/36:0 | 1.799 | 0.001 |
| Cer 17:1;O3/38:2(2OH) | 1.797 | 0.010 |
| SE 24:1;O4/25:1;O A | 1.795 | 0.014 |
| SL 15:3;O/34:6 | 1.794 | 0.005 |
| TG 52:3\|TG 16:0_18:1_18:2 | 1.791 | 0.001 |
| SHexCer 12:1;O2/22:0;O B | 1.788 | 0.007 |
| TG 8:0_12:0_38:0 | 1.784 | 0.023 |
| TG O-50:1\|TG O-16:0_16:0_18:1 | 1.782 | 0.001 |
| TG 8:0_9:0_36:3 B | 1.78 | 0.002 |
| PC 34:1\|PC 16:0_18:1 B | 1.779 | 0.001 |
| TG 8:0_14:0_38:5 | 1.777 | 0.012 |
| TG 8:0_8:0_29:1 | 1.775 | 0.000 |
| SL 16:1;O/36:1;O B | 1.774 | 0.000 |
| SE 24:1;O4/27:0;O | 1.774 | 0.002 |
| SM 12:1;O2/30:1 A | 1.766 | 0.008 |
| DG 49:4 | 1.765 | 0.008 |
| TG 8:0_8:0_28:2 C | 1.765 | 0.013 |
| PC 36:1 | 1.764 | 0.002 |
| VAE 26:3 | 1.763 | 0.001 |
| SL 15:0;O/36:0;O | 1.755 | 0.002 |
| TG O-20:0_12:0_18:0 | 1.755 | 0.006 |
| TG 50:6\|TG 12:0_18:2_20:4 | 1.754 | 0.002 |
| Cer 21:1;O3/38:1(2OH) A | 1.749 | 0.002 |
| SL 22:1;O/36:2;O | 1.749 | 0.005 |
| SM 34:1;O2 B | 1.743 | 0.000 |
| Cer 12:0;O2/22:0;O | 1.742 | 0.043 |
| LPC 16:0 | 1.738 | 0.003 |
| TG 10:0_10:0_24:1 | 1.733 | 0.003 |
| TG 8:0_8:0_30:3 | 1.731 | 0.001 |
| NAE 22:5 | 1.73 | 0.000 |
| TG 48:3\|TG 12:0_18:1_18:2 | 1.728 | 0.002 |
| TG 16:0_20:0_18:1 | 1.727 | 0.001 |
| SE 24:1;O4/26:0;O | 1.726 | 0.009 |
| TG 50:3\|TG 16:0_16:1_18:2 | 1.725 | 0.001 |
| TG 8:0_14:0_38:6 | 1.716 | 0.003 |
| TG 17:0_15:1_17:1 | 1.713 | 0.002 |
| TG 8:0_8:0_38:1 | 1.707 | 0.003 |
| TG 8:0_8:0_36:7 | 1.704 | 0.002 |
| ST 28:2;O | 1.703 | 0.001 |
| Cer 12:0;O2/30:0;O | 1.693 | 0.019 |
| Cer 13:0;O2/34:3 | 1.688 | 0.018 |
| SL 17:0;O/36:0;O B | 1.686 | 0.043 |
| TG 8:0_8:0_32:4 | 1.685 | 0.002 |
| TG 8:0_8:0_38:7 | 1.685 | 0.012 |
| DG 18:0_28:0 | 1.682 | 0.005 |
| Cer 17:1;O3/38:1(2OH) | 1.681 | 0.007 |
| Cer 12:0;O3/36:0(2OH) | 1.677 | 0.038 |
| TG 8:0_8:0_38:0 B | 1.671 | 0.001 |

| | | |
|---|---|---|
| TG 8:0_8:0_23:0 | 1.671 | 0.007 |
| SL 15:3;O/32:6 B | 1.67 | 0.002 |
| PC 34:2\|PC 16:0_18:2 B | 1.669 | 0.001 |
| HexCer 16:0;O2/25:1 | 1.666 | 0.005 |
| TG 8:0_8:0_32:0 | 1.664 | 0.003 |
| TG 8:0_8:0_20:0 B | 1.663 | 0.003 |
| Cer 12:2;O2/39:11;O2 A | 1.661 | 0.001 |
| SM 35:1;O3 | 1.661 | 0.003 |
| HexCer 34:1;O3\|HexCer 18:1;O2/16:0;C | 1.66 | 0.002 |
| SM 12:1;O2/28:0 | 1.656 | 0.001 |
| HexCer 41:1;O2\|HexCer 18:1;O2/23:0 | 1.655 | 0.001 |
| SL 13:1;O/36:3;O | 1.653 | 0.006 |
| HexCer 34:1;O3\|HexCer 18:1;O2/16:0;C | 1.649 | 0.001 |
| NAOrn 27:1 A | 1.649 | 0.003 |
| TG 18:0_14:1_20:2 | 1.647 | 0.001 |
| TG 56:6\|TG 18:1_18:2_20:3 | 1.644 | 0.005 |
| NAE 20:2 | 1.642 | 0.000 |
| SE 28:2/21:4 | 1.638 | 0.019 |
| TG 8:0_8:0_26:0 | 1.63 | 0.000 |
| TG O-21:1_17:0_18:0 | 1.629 | 0.016 |
| Cer 23:0;O3/38:0(2OH) | 1.626 | 0.016 |
| TG 50:6\|TG 12:0_16:0_22:6 | 1.621 | 0.004 |
| Cer 13:1;O3/38:0(2OH) | 1.62 | 0.002 |
| TG O-18:0_16:1_16:1 | 1.62 | 0.040 |
| PC 38:5\|PC 18:0_20:5 | 1.619 | 0.006 |
| TG 13:1_15:1_19:4 | 1.618 | 0.005 |
| Cer 13:0;O2/32:4;O | 1.612 | 0.004 |
| Cer 42:1;O2\|Cer 18:1;O2/24:0 A | 1.611 | 0.001 |
| Cer 25:1;O3/38:0(2OH) | 1.61 | 0.001 |
| TG 18:0_20:1_18:3 | 1.61 | 0.002 |
| ST 29:1;O A | 1.609 | 0.003 |
| TG 8:0_9:0_38:0 | 1.608 | 0.023 |
| HexCer 17:0;O2/22:5;O | 1.605 | 0.001 |
| LPC 18:1 C | 1.601 | 0.006 |
| TG O-52:0\|TG O-18:0_16:0_18:0 | 1.599 | 0.022 |
| TG 8:0_8:0_22:0 C | 1.598 | 0.002 |
| TG O-50:0\|TG O-18:0_14:0_18:0 | 1.596 | 0.003 |
| AHexCer (O-12:0)12:1;O2/16:3;O | 1.596 | 0.009 |
| TG 54:6\|TG 18:1_18:2_18:3 | 1.591 | 0.003 |
| TG 8:0_8:0_30:4 | 1.586 | 0.001 |
| TG 50:5\|TG 12:0_18:1_20:4 | 1.583 | 0.001 |
| TG 11:0_18:0_20:1 | 1.581 | 0.001 |
| SL 20:1;O/36:0;O B | 1.578 | 0.005 |
| TG 8:0_8:0_37:1 | 1.576 | 0.001 |
| Cer 13:0;O2/30:4;O | 1.571 | 0.001 |
| TG 8:0_8:0_34:6 | 1.57 | 0.013 |
| TG 8:0_8:0_34:3 | 1.569 | 0.005 |
| PC 36:2\|PC 18:1_18:1 | 1.566 | 0.001 |
| Cer 19:1;O3/38:2(2OH) | 1.566 | 0.030 |

| | | |
|---|---|---|
| TG 54:3\|TG 18:0_18:1_18:2 | 1.56 | 0.001 |
| TG O-46:2\|TG O-16:0_12:0_18:2 | 1.56 | 0.015 |
| SL 18:1;O/36:3;O | 1.556 | 0.009 |
| TG 56:7\|TG 16:0_18:1_22:6 | 1.553 | 0.008 |
| Cer 41:1;O2\|Cer 18:1;O2/23:0 B | 1.551 | 0.001 |
| LPC 18:1 A | 1.551 | 0.002 |
| TG 8:0_9:0_34:3 A | 1.551 | 0.005 |
| PI 38:4 | 1.551 | 0.036 |
| HexCer 18:1;O2/24:0 | 1.55 | 0.002 |
| SL 18:0;O/36:0;O | 1.55 | 0.012 |
| TG 8:0_13:0_38:8 | 1.548 | 0.042 |
| Cer 14:0;O2/38:3 | 1.547 | 0.007 |
| SM 39:3;O3 | 1.541 | 0.000 |
| TG O-48:0\|TG O-18:0_14:0_16:0 | 1.541 | 0.013 |
| SL 13:1;O/34:6 | 1.54 | 0.002 |
| Cer 42:2;O2\|Cer 18:1;O2/24:1 B | 1.535 | 0.002 |
| TG 56:8\|TG 16:0_18:2_22:6 | 1.534 | 0.005 |
| TG 45:0\|TG 14:0_15:0_16:0 | 1.534 | 0.016 |
| SM 12:1;O2/30:1 B | 1.532 | 0.000 |
| TG 8:0_8:0_36:0 | 1.531 | 0.004 |
| PC O-15:4_26:7 B | 1.53 | 0.002 |
| SL 13:0;O/36:0;O | 1.526 | 0.001 |
| Cer 18:0;O2/24:0 | 1.526 | 0.014 |
| DG 51:10 | 1.526 | 0.015 |
| Cer 15:3;O2/36:6 | 1.523 | 0.014 |
| TG 58:9\|TG 18:1_18:2_22:6 | 1.521 | 0.001 |
| Cer 23:1;O3/38:1(2OH) | 1.521 | 0.005 |
| TG 11:0_17:0_19:1 | 1.519 | 0.003 |
| Cer 15:3;O2/9:0 | 1.515 | 0.006 |
| TG 8:0_10:0_38:2 | 1.515 | 0.014 |
| CAR 26:7 | 1.506 | 0.007 |
| PC 38:3\|PC 18:0_20:3 | 1.505 | 0.002 |
| TG O-46:0\|TG O-18:0_12:0_16:0 | 1.505 | 0.003 |
| Cer 12:0;O2/38:0;O B | 1.505 | 0.009 |
| TG 8:0_10:0_38:4 | 1.504 | 0.003 |
| HexCer 16:0;O2/26:1 | 1.499 | 0.002 |
| TG 8:0_8:0_30:2 | 1.493 | 0.005 |
| Cer 12:2;O2/38:11;O2 | 1.491 | 0.003 |
| ST 27:2;O F | 1.491 | 0.016 |
| TG 18:1_18:1_18:1 | 1.489 | 0.004 |
| LPC 18:0 | 1.485 | 0.003 |
| SE 28:2/22:0 | 1.481 | 0.036 |
| CE 20:3 | 1.48 | 0.013 |
| Cer 21:1;O3/38:0(2OH) | 1.478 | 0.024 |
| ST 27:2;O D | 1.474 | 0.000 |
| TG 17:0_15:1_19:1 | 1.468 | 0.001 |
| LPC 18:2 B | 1.467 | 0.010 |
| SE 28:2/18:0 A | 1.462 | 0.023 |
| PC O-9:0_28:7 | 1.461 | 0.000 |

| | | |
|---|---|---|
| PC O-15:4_24:6 | 1.46 | 0.004 |
| Cer 17:0;O2/25:0;O | 1.453 | 0.001 |
| DG 49:9 | 1.453 | 0.012 |
| SM 12:1;O2/36:0 | 1.45 | 0.001 |
| Cer 40:1;O2\|Cer 18:1;O2/22:0 A | 1.44 | 0.003 |
| TG O-54:1\|TG O-20:0_16:0_18:1 | 1.439 | 0.017 |
| ST 27:2;O C | 1.434 | 0.010 |
| Cer 34:1;O2\|Cer 18:1;O2/16:0 | 1.433 | 0.032 |
| LPC 18:1 B | 1.43 | 0.001 |
| SE 24:1;O4/28:0;O A | 1.43 | 0.009 |
| Cer 13:0;O2/34:3;O | 1.428 | 0.025 |
| LPC 18:2 A | 1.425 | 0.004 |
| TG O-16:1_17:0_17:0 | 1.42 | 0.005 |
| LPC 20:3/0:0 | 1.415 | 0.011 |
| TG 8:0_8:0_28:0 | 1.414 | 0.000 |
| PC 32:0\|PC 16:0_16:0 C | 1.412 | 0.008 |
| TG 56:7\|TG 18:1_18:2_20:4 | 1.406 | 0.001 |
| TG O-48:1\|TG O-18:0_12:0_18:1 | 1.406 | 0.003 |
| TG 8:0_8:0_24:2 C | 1.406 | 0.007 |
| PC 38:4\|PC 18:0_20:4 C | 1.404 | 0.001 |
| PC 36:4\|PC 16:0_20:4 C | 1.391 | 0.021 |
| PI 36:1 | 1.389 | 0.010 |
| Cer 12:2;O2/40:12;O2 | 1.388 | 0.008 |
| BMP 8:0_24:1 | 1.384 | 0.009 |
| TG 8:0_8:0_30:1 A | 1.383 | 0.000 |
| Cer 41:1;O2\|Cer 18:1;O2/23:0 A | 1.383 | 0.013 |
| HexCer 16:0;O3/18:0(2OH) | 1.376 | 0.002 |
| TG 16:0_16:0_19:1 | 1.373 | 0.000 |
| TG 8:0_8:0_28:1 | 1.373 | 0.001 |
| Cer 42:2;O2\|Cer 18:1;O2/24:1 A | 1.369 | 0.002 |
| Cer 48:3;O2\|Cer 16:2;O2/32:1 | 1.365 | 0.002 |
| Cer 12:1;O3/35:0(2OH) | 1.365 | 0.011 |
| CAR 16:0 B | 1.363 | 0.002 |
| NAGly 13:0;O(FA 9:0) | 1.355 | 0.001 |
| TG 18:1_20:2_22:4 | 1.354 | 0.005 |
| TG 8:0_8:0_36:1 B | 1.353 | 0.001 |
| SE 28:1/19:0 | 1.351 | 0.001 |
| TG 8:0_9:0_28:2 | 1.35 | 0.045 |
| TG 8:0_12:0_38:10 | 1.346 | 0.004 |
| NAGly 10:0;O(FA 12:0) | 1.343 | 0.001 |
| PC 6:0_32:6 | 1.343 | 0.007 |
| TG 8:0_10:0_38:8 | 1.342 | 0.002 |
| NAGly 20:1 | 1.338 | 0.003 |
| PC O-8:0_9:0 | 1.335 | 0.002 |
| NAOrn 27:1 B | 1.332 | 0.016 |
| PC O-8:0_26:1 | 1.322 | 0.046 |
| AHexCer (O-12:0)12:2;O2/12:0;O C | 1.318 | 0.043 |
| SL 12:0;O/36:0;O | 1.317 | 0.006 |
| Cer 19:0;O3/38:0(2OH) | 1.311 | 0.030 |

| | | |
|---|---|---|
| TG 8:0_8:0_25:0 | 1.309 | 0.001 |
| TG 8:0_9:0_34:3 B | 1.309 | 0.005 |
| TG 12:0_18:0_18:2 | 1.309 | 0.006 |
| LPC 16:1 | 1.304 | 0.004 |
| TG 8:0_11:0_38:1 | 1.301 | 0.000 |
| SM 12:1;O2/38:0 | 1.301 | 0.009 |
| TG 8:0_11:1_8:0;O2 | 1.3 | 0.003 |
| LPC 20:4 B | 1.291 | 0.007 |
| TG 8:0_26:1_24:3 | 1.29 | 0.000 |
| TG 17:0_17:1_19:1 | 1.289 | 0.002 |
| Cer 12:2;O2/15:3 | 1.289 | 0.004 |
| SL 22:1;O/36:3;O | 1.289 | 0.017 |
| CE 20:4 | 1.283 | 0.009 |
| TG 10:0_10:1_8:0;O2 | 1.281 | 0.002 |
| TG O-46:1\|TG O-16:0_12:0_18:1 | 1.276 | 0.022 |
| PC 36:1\|PC 18:0_18:1 A | 1.274 | 0.002 |
| SL 13:1;O/32:5 | 1.274 | 0.009 |
| Cer 12:1;O3/32:1(2OH) | 1.274 | 0.018 |
| SE 28:2/21:2 | 1.272 | 0.019 |
| TG 8:0_10:0_38:3 | 1.271 | 0.031 |
| TG 54:6\|TG 16:0_18:2_20:4 | 1.266 | 0.001 |
| Cer 42:1;O2\|Cer 18:1;O2/24:0 B | 1.266 | 0.003 |
| Cer 12:0;O3/33:0(2OH) | 1.266 | 0.046 |
| SE 28:2/24:4 | 1.265 | 0.004 |
| DG 51:6 | 1.262 | 0.013 |
| TG O-13:0_13:0_18:0 | 1.258 | 0.012 |
| DG 47:7 | 1.255 | 0.003 |
| SE 28:2/19:0 B | 1.255 | 0.018 |
| LPC 22:6 | 1.254 | 0.005 |
| PC 40:7 | 1.252 | 0.013 |
| CE 21:0 | 1.25 | 0.000 |
| HexCer 16:0;O2/19:5 | 1.247 | 0.000 |
| TG 18:0_18:1_18:1 | 1.246 | 0.003 |
| TG 8:0_8:0_30:1 B | 1.242 | 0.000 |
| TG 8:0_8:0_24:1 | 1.241 | 0.001 |
| Cer 41:0;O3\|Cer 17:0;O2/24:0;O | 1.241 | 0.004 |
| VAE 21:0 | 1.241 | 0.005 |
| PC 38:5 B | 1.238 | 0.002 |
| NAGly 20:0;O(FA 28:1) | 1.238 | 0.004 |
| SM 12:1;O2/23:0 | 1.237 | 0.004 |
| HexCer 16:0;O2/21:2 | 1.236 | 0.029 |
| PC 6:0_30:1 | 1.233 | 0.006 |
| DG 54:2 | 1.232 | 0.004 |
| TG 8:0_9:0_34:2 | 1.226 | 0.002 |
| PI 14:0 | 1.221 | 0.003 |
| SL 12:1;O/24:1 B | 1.22 | 0.004 |
| NAOrn 19:0;O B | 1.22 | 0.007 |
| NAGly 16:2;O(FA 28:5) | 1.218 | 0.006 |
| SL 19:0;O/36:0;O | 1.208 | 0.002 |

| | | |
|---|---|---|
| LPC O-19:3 | 1.207 | 0.001 |
| Cer 42:1;O2\|Cer 18:1;O2/24:0 C | 1.206 | 0.000 |
| LPC 20:4 A | 1.206 | 0.005 |
| SL 14:0;O/36:0;O | 1.203 | 0.023 |
| SM 12:1;O2/34:0 | 1.194 | 0.000 |
| HexCer 34:1;O2\|HexCer 18:1;O2/16:0 | 1.194 | 0.011 |
| HexCer 16:0;O2/24:0;O | 1.188 | 0.004 |
| Cer 12:0;O2/34:0;O | 1.186 | 0.031 |
| HexCer 18:1;O2/22:0 | 1.184 | 0.000 |
| Cer 41:0;O2\|Cer 18:0;O2/23:0 | 1.183 | 0.003 |
| TG 8:0_14:0_38:3 | 1.171 | 0.006 |
| DG 15:3_19:4 | 1.166 | 0.007 |
| LPC O-15:3 C | 1.165 | 0.001 |
| TG 8:0_8:1_10:0;O2 | 1.163 | 0.014 |
| Cer 42:2;O2\|Cer 18:1;O2/24:1 C | 1.163 | 0.017 |
| SE 28:2/19:4 | 1.157 | 0.019 |
| ST 27:2;O A | 1.155 | 0.002 |
| LPC 14:0/0:0 | 1.154 | 0.009 |
| SL 12:1;O/36:0;O | 1.153 | 0.005 |
| Cer 40:1;O2\|Cer 18:1;O2/22:0 B | 1.152 | 0.008 |
| Cer 27:3;O2/11:0 | 1.152 | 0.019 |
| PC 36:5\|PC 16:0_20:5 B | 1.151 | 0.001 |
| Cer 21:1;O3/38:1(2OH) B | 1.142 | 0.017 |
| SL 20:1;O/36:0;O A | 1.14 | 0.020 |
| TG 12:0_14:0_16:0;O | 1.137 | 0.000 |
| DG 50:0 | 1.136 | 0.045 |
| Cer 22:1;O3/38:0(2OH) | 1.132 | 0.008 |
| Cer 12:0;O2/29:1 | 1.131 | 0.022 |
| DG O-37:1\|DG O-23:0_14:1 | 1.13 | 0.000 |
| Cer 14:2;O3/8:0(2OH) | 1.13 | 0.006 |
| SM 34:1;O3 | 1.123 | 0.004 |
| TG 50:4\|TG 14:0_18:2_18:2 | 1.122 | 0.000 |
| SE 28:2/19:5 | 1.122 | 0.035 |
| SM 34:0;O3 | 1.119 | 0.000 |
| DG 50:1 | 1.118 | 0.001 |
| SL 16:1;O/36:1;O A | 1.118 | 0.043 |
| PE 21:1 | 1.115 | 0.000 |
| VAE 25:0 A | 1.115 | 0.020 |
| NAGly 26:2 B | 1.11 | 0.015 |
| NATau 22:3;O | 1.107 | 0.003 |
| Cer 18:1;O3/38:0(2OH) | 1.107 | 0.013 |
| Cer 27:0;O2/38:4 | 1.105 | 0.021 |
| DG 52:0 | 1.103 | 0.002 |
| TG 8:0_11:0_38:4 | 1.099 | 0.021 |
| SM 43:4;O2\|SM 17:3;O2/26:1 | 1.098 | 0.005 |
| PE O-8:0_15:1 | 1.097 | 0.006 |
| LPE 16:0 | 1.097 | 0.007 |
| DG 52:1 | 1.097 | 0.021 |
| Cer 13:2;O2/28:6 | 1.094 | 0.006 |

| | | |
|---|---|---|
| CAR 27:0 | 1.093 | 0.011 |
| Cer 40:0;O2\|Cer 18:0;O2/22:0 | 1.093 | 0.021 |
| DG 48:0 | 1.092 | 0.040 |
| NATau 24:4;O | 1.09 | 0.002 |
| PC 38:6 B | 1.089 | 0.002 |
| LPC 15:1 | 1.086 | 0.012 |
| SM 41:3;O3 | 1.084 | 0.000 |
| CAR 16:1 | 1.082 | 0.011 |
| TG 14:0_16:0_20:0 | 1.079 | 0.020 |
| TG 18:0_20:1_18:2 | 1.078 | 0.011 |
| TG 8:0_9:0_38:2 | 1.076 | 0.003 |
| ST 27:1;O E | 1.071 | 0.000 |
| NATau 20:0;O | 1.064 | 0.005 |
| Cer 12:1;O2/41:12;O2 B | 1.064 | 0.019 |
| MGDG 9:0_14:1 | 1.063 | 0.000 |
| TG 8:0_9:0_38:5 | 1.063 | 0.006 |
| SL 13:1;O/32:6 | 1.062 | 0.010 |
| NAGly 18:1 B | 1.061 | 0.000 |
| Cer 42:3;O3\|Cer 18:1;O2/24:2;O | 1.051 | 0.020 |
| TG 8:0_8:0_24:2 A | 1.051 | 0.030 |
| Cer 12:0;O3/8:0(2OH) | 1.049 | 0.002 |
| PC 38:5 A | 1.048 | 0.001 |
| Cer 42:1;O3\|Cer 18:1;O2/24:0;O | 1.047 | 0.020 |
| ST 27:2;O B | 1.045 | 0.002 |
| Cer 12:2;O2/36:10;O2 | 1.045 | 0.006 |
| Cer 12:1;O2/39:11;O2 B | 1.044 | 0.013 |
| NAPhe 26:6;O | 1.043 | 0.000 |
| Cer 14:0;O3/38:0(2OH) | 1.035 | 0.009 |
| NAE 7:0 | 1.03 | 0.007 |
| TG 8:0_8:0_34:1 | 1.028 | 0.001 |
| NAPhe 20:4;O | 1.026 | 0.033 |
| TG O-40:0\|TG O-18:0_10:0_12:0 | 1.019 | 0.011 |
| Cer 16:0;O3/38:0(2OH) | 1.019 | 0.014 |
| HexCer 17:0;O2/26:3;O | 1.016 | 0.018 |
| CE 24:6 | 1.014 | 0.001 |
| TG 13:0_18:0_18:0 | 1.014 | 0.010 |
| CE 21:3 | 1.013 | 0.000 |
| TG 20:0_18:1_18:1 | 1.013 | 0.012 |
| Cer 13:1;O3/38:1(2OH) | 1.006 | 0.044 |
| Cer 13:0;O2/24:2;O | 1.003 | 0.036 |
| ST 27:1;O I | 1 | 0.001 |
| TG 8:0_8:0_38:6 | 0.993 | 0.022 |
| TG 8:0_12:0_38:3 | 0.99 | 0.008 |
| ST 29:2;O | 0.987 | 0.004 |
| Cer 12:2;O2/19:2 | 0.986 | 0.001 |
| Cer 12:0;O2/36:0;O | 0.985 | 0.002 |
| TG 8:0_8:0_32:1 | 0.985 | 0.032 |
| DG 49:7 | 0.983 | 0.000 |
| Cer 12:1;O3/38:3(2OH) | 0.982 | 0.033 |

| | | |
|---|---|---|
| NAE 20:1 | 0.979 | 0.000 |
| PC O-33:6 | 0.976 | 0.023 |
| NAGlySer 13:0;O(FA 28:5) | 0.976 | 0.025 |
| NAGly 11:0 | 0.971 | 0.037 |
| Cer 15:3;O2/18:5 | 0.97 | 0.000 |
| NAGly 26:2 A | 0.969 | 0.004 |
| Cer 13:0;O2/22:6;O C | 0.966 | 0.008 |
| MGDG O-17:2_4:0 | 0.963 | 0.007 |
| NAGlySer 9:0;O(FA 28:5) | 0.956 | 0.019 |
| TG O-44:0\|TG O-18:0_12:0_14:0 | 0.956 | 0.021 |
| Cer 15:3;O3/18:5(2OH) | 0.955 | 0.000 |
| SM 12:0;O2/21:0 | 0.954 | 0.025 |
| NAE 15:1 | 0.949 | 0.009 |
| PC 32:1 | 0.948 | 0.016 |
| NAGly 15:3 A | 0.944 | 0.015 |
| TG 8:0_9:0_36:3 A | 0.942 | 0.005 |
| TG 8:0_8:0_18:0 | 0.939 | 0.017 |
| LPC 30:3 | 0.936 | 0.001 |
| NAOrn 27:1 C | 0.936 | 0.010 |
| HexCer 16:0;O2/26:4;O | 0.934 | 0.003 |
| TG 8:0_8:0_21:0 | 0.934 | 0.016 |
| NAGly 22:3 | 0.934 | 0.025 |
| DG 42:0 | 0.931 | 0.013 |
| VAE 21:2 B | 0.929 | 0.003 |
| Cer 12:0;O2/17:0;O A | 0.929 | 0.015 |
| SM 32:7;O2 | 0.929 | 0.043 |
| Cer 12:1;O3/24:5(2OH) B | 0.928 | 0.018 |
| PE 20:1 | 0.926 | 0.004 |
| SE 28:2/24:0 | 0.917 | 0.002 |
| LPE 18:0 | 0.917 | 0.031 |
| Cer 12:2;O2/19:4 | 0.911 | 0.005 |
| PC 36:4\|PC 18:2_18:2 B | 0.907 | 0.032 |
| SL 13:2;O/36:6 | 0.9 | 0.039 |
| C17-Sphingosine | 0.899 | 0.039 |
| Cer 12:2;O2/41:12;O2 | 0.896 | 0.028 |
| SE 28:2/22:2 | 0.895 | 0.020 |
| DG O-21:5_2:0 | 0.893 | 0.005 |
| TG 8:0_28:1_19:2 | 0.889 | 0.002 |
| ST 27:2;O H | 0.886 | 0.016 |
| HexCer 16:0;O2/22:0;O | 0.884 | 0.002 |
| TG 8:0_11:0_38:3 | 0.88 | 0.000 |
| LPE 20:4 | 0.878 | 0.035 |
| DG O-19:5_15:3 | 0.871 | 0.010 |
| DG 46:0 | 0.87 | 0.005 |
| Cer 12:1;O2/22:6 | 0.869 | 0.010 |
| SL 13:1;O/30:6 | 0.866 | 0.000 |
| DG 51:8 | 0.866 | 0.001 |
| Cer 12:0;O2/29:1;O | 0.866 | 0.009 |
| DG 37:5 | 0.861 | 0.035 |

| | | |
|---|---|---|
| NAGly 22:6;O(FA 26:6) | 0.86 | 0.009 |
| MGDG O-26:6_7:0 | 0.853 | 0.019 |
| DG 51:7 | 0.845 | 0.023 |
| DG 54:1 | 0.844 | 0.002 |
| NAGly 18:1 A | 0.843 | 0.008 |
| SM 45:0;O2 | 0.84 | 0.022 |
| NAOrn 24:6 | 0.836 | 0.017 |
| TG 54:4\|TG 18:1_18:1_18:2 B | 0.835 | 0.009 |
| TG 8:0_15:4_18:5 | 0.829 | 0.000 |
| TG 50:2\|TG 16:0_16:0_18:2 | 0.818 | 0.001 |
| TG 8:0_8:0_30:0 | 0.811 | 0.044 |
| TG 43:2\|TG 12:0_16:0_15:2 | 0.807 | 0.022 |
| CAR 28:6 D | 0.803 | 0.002 |
| DG 39:6 | 0.795 | 0.002 |
| NAGlySer 9:0;O(FA 26:4) | 0.787 | 0.048 |
| DG O-15:1_7:0 | 0.772 | 0.019 |
| CAR 17:1 | 0.765 | 0.017 |
| NAOrn 19:0;O A | 0.763 | 0.036 |
| TG 8:0_9:0_24:2 | 0.746 | 0.045 |
| Cer 13:0;O2/22:6 | 0.743 | 0.022 |
| LPE 22:6 | 0.726 | 0.022 |
| DG O-20:5_4:0 | 0.721 | 0.001 |
| PS 21:3 | 0.718 | 0.005 |
| TG 8:0_8:0_24:2 B | 0.718 | 0.028 |
| NAPhe 17:4 | 0.716 | 0.005 |
| Carnosine | 0.711 | 0.015 |
| Cer 12:0;O2/29:0 | 0.708 | 0.010 |
| SL 20:3;O/18:3;O | 0.703 | 0.007 |
| NAOrn 14:0 | 0.696 | 0.011 |
| DG O-22:5_3:0 | 0.694 | 0.003 |
| NATau 24:6;O | 0.694 | 0.016 |
| DG 55:8 | 0.686 | 0.040 |
| TG 52:4\|TG 16:0_18:2_18:2 | 0.684 | 0.042 |
| DG 53:7 | 0.681 | 0.003 |
| DG 49:6 | 0.674 | 0.014 |
| SPB 16:1;O3 D | 0.673 | 0.008 |
| DG 17:0 | 0.669 | 0.002 |
| Cer 12:2;O2/18:5 | 0.666 | 0.001 |
| LPC 15:2 | 0.654 | 0.024 |
| NAGlySer 11:0;O | 0.65 | 0.026 |
| ST 27:1;O H | 0.646 | 0.009 |
| TG 50:3\|TG 16:0_16:0_18:3 | 0.646 | 0.031 |
| PC 36:4\|PC 16:0_20:4 A | 0.642 | 0.017 |
| NATau 9:0 A | 0.629 | 0.002 |
| Cer 16:3;O3/32:6(2OH) | 0.629 | 0.003 |
| DG 55:10 | 0.628 | 0.004 |
| ST 27:1;O F | 0.627 | 0.001 |
| SPB 18:1;O3 C | 0.626 | 0.019 |
| Cer 12:1;O2/39:11;O2 C | 0.624 | 0.004 |

| | | |
|---|---|---|
| DG 53:8 | 0.622 | 0.007 |
| SM 37:6;O2 | 0.619 | 0.002 |
| Cer 12:0;O2/32:0;O | 0.608 | 0.019 |
| Cer 12:0;O3/38:0(2OH) | 0.606 | 0.027 |
| DG 51:9 | 0.593 | 0.011 |
| NATau 16:4 C | 0.593 | 0.016 |
| TG 54:3\|TG 16:0_16:0_22:3 | 0.592 | 0.040 |

| IgE + IgG > IgG | | |
|---|---|---|
| ***Metabolite*** | ***Log2(Fold Change)*** | ***P-value*** |
| NATau 20:0;O | -0.712 | 0.026 |
| NAGlySer 18:2;O H | -0.616 | 0.012 |

| IgG > IgE + IgG | | |
|---|---|---|
| ***Metabolite*** | ***Log2(Fold Change)*** | ***P-value*** |
| PC O-38:5\|PC O-18:1_20:4 | 4.314 | 0.006 |
| PC 36:4\|PC 16:0_20:4 E | 2.983 | 0.048 |
| PC 38:4\|PC 18:0_20:4 B | 2.492 | 0.029 |
| SM 38:4;O3 | 2.434 | 0.003 |
| PC 38:6\|PC 16:0_22:6 C | 2.427 | 0.046 |
| Cer 13:0;O2/38:1 | 2.348 | 0.004 |
| SM 36:1;O2 A | 2.061 | 0.008 |
| PC 36:4\|PC 16:0_20:4 B | 1.932 | 0.043 |
| CAR 27:0 | 1.899 | 0.027 |
| DG 43:8 | 1.811 | 0.004 |
| HexCer 16:0;O2/28:5;O | 1.757 | 0.009 |
| HexCer 17:0;O2/24:5;O | 1.708 | 0.028 |
| TG 8:0_9:0_26:2 | 1.661 | 0.005 |
| Cer 23:0;O3/38:0(2OH) | 1.652 | 0.044 |
| SM 34:1;O2\|SM 18:1;O2/16:0 A | 1.59 | 0.032 |
| TG 8:0_8:0_25:0 | 1.57 | 0.000 |
| PC O-15:4_26:7 A | 1.558 | 0.044 |
| PC 32:0\|PC 16:0_16:0 B | 1.494 | 0.026 |
| TG 50:1\|TG 16:0_16:0_18:1 | 1.49 | 0.006 |
| ST 27:2;O F | 1.481 | 0.009 |
| PC 38:6 | 1.48 | 0.035 |
| SM 42:2;O2\|SM 18:1;O2/24:1 | 1.469 | 0.031 |
| CE 17:1 | 1.447 | 0.011 |
| TG 8:0_10:0_38:4 | 1.442 | 0.003 |
| TG 54:3\|TG 18:0_18:1_18:2 | 1.441 | 0.003 |
| TG 18:1_20:2_22:4 | 1.44 | 0.040 |
| HexCer 16:0;O3/18:0(2OH) | 1.439 | 0.002 |
| PC O-10:0_28:6 B | 1.423 | 0.031 |
| PC O-10:0_28:6 A | 1.393 | 0.019 |
| Cer 47:2;O2\|Cer 20:1;O2/27:1 | 1.384 | 0.003 |
| TG 42:2\|TG 12:0_12:0_18:2 | 1.375 | 0.008 |
| SM 12:1;O2/36:0 | 1.354 | 0.019 |
| CE 18:3 | 1.342 | 0.023 |
| TG 38:0\|TG 12:0_12:0_14:0 | 1.335 | 0.011 |

| | | |
|---|---|---|
| TG 52:2\|TG 16:0_18:1_18:1 | 1.333 | 0.004 |
| TG 40:1\|TG 10:0_12:0_18:1 | 1.332 | 0.018 |
| TG 10:0_15:0_20:2 | 1.322 | 0.030 |
| PC 36:4\|PC 16:0_20:4 D | 1.322 | 0.030 |
| TG 48:0\|TG 14:0_16:0_18:0 | 1.318 | 0.029 |
| TG 42:1\|TG 12:0_12:0_18:1 | 1.317 | 0.012 |
| PC 36:2\|PC 18:0_18:2 | 1.313 | 0.030 |
| SE 28:2/18:1 | 1.311 | 0.004 |
| TG 14:0_16:0_16:2 | 1.305 | 0.013 |
| TG 56:6\|TG 18:1_18:2_20:3 | 1.303 | 0.015 |
| TG 46:1\|TG 12:0_16:0_18:1 | 1.298 | 0.007 |
| Cer 15:3;O2/9:0 | 1.296 | 0.015 |
| SM 12:1;O2/30:1 B | 1.282 | 0.022 |
| SL 16:1;O/36:3;O | 1.281 | 0.009 |
| TG 44:0\|TG 12:0_14:0_18:0 | 1.28 | 0.012 |
| Cer 12:0;O2/22:0;O | 1.276 | 0.010 |
| SL 22:1;O/36:3;O | 1.276 | 0.048 |
| Cer 13:2;O2/38:6 | 1.275 | 0.017 |
| TG 8:0_8:0_26:2 | 1.274 | 0.009 |
| SL 20:1;O/36:0 | 1.274 | 0.012 |
| SE 28:2/21:3 | 1.272 | 0.008 |
| PC 36:3\|PC 16:0_20:3 A | 1.27 | 0.045 |
| ST 28:2;O | 1.265 | 0.040 |
| TG 54:5\|TG 16:0_18:1_20:4 | 1.262 | 0.007 |
| TG 8:0_8:0_20:0 B | 1.26 | 0.013 |
| TG 58:9\|TG 18:1_18:2_22:6 | 1.257 | 0.019 |
| TG 52:0\|TG 16:0_18:0_18:0 | 1.254 | 0.020 |
| TG 54:5\|TG 18:1_18:2_18:2 | 1.25 | 0.012 |
| Cer 46:2;O2\|Cer 18:1;O2/28:1 | 1.248 | 0.016 |
| PC 36:1\|PC 18:0_18:1 B | 1.244 | 0.038 |
| TG 36:0\|TG 10:0_12:0_14:0 | 1.242 | 0.012 |
| ST 27:2;O E | 1.239 | 0.023 |
| TG 40:0\|TG 10:0_14:0_16:0 | 1.238 | 0.013 |
| TG 8:0_8:0_32:6 B | 1.236 | 0.007 |
| TG 11:0_11:0_21:0 | 1.234 | 0.024 |
| BMP 8:0_24:1 | 1.234 | 0.049 |
| TG 48:2\|TG 14:0_16:0_18:2 | 1.23 | 0.015 |
| Cer 18:0;O2/24:0 | 1.228 | 0.008 |
| TG 14:0_16:0_16:0 | 1.223 | 0.025 |
| SL 15:3;O/32:6 A | 1.222 | 0.029 |
| TG 13:0_13:0_18:0 | 1.215 | 0.021 |
| Cer 19:0;O3/38:0(2OH) | 1.213 | 0.030 |
| NAGlySer 13:1;O(FA 26:6) | 1.208 | 0.024 |
| SM 12:1;O2/38:0 | 1.205 | 0.031 |
| TG 8:0_8:0_22:0 C | 1.202 | 0.010 |
| Cer 19:1;O3/38:1(2OH) | 1.201 | 0.001 |
| TG 50:3\|TG 16:0_16:1_18:2 | 1.201 | 0.015 |
| TG 40:2\|TG 10:0_12:0_18:2 | 1.194 | 0.008 |
| Cer 14:0;O2/38:3 | 1.191 | 0.007 |

| | | |
|---|---|---|
| TG O-52:1\|TG O-18:0_16:0_18:1 | 1.191 | 0.013 |
| TG 8:0_8:0_28:2 B | 1.185 | 0.012 |
| TG 8:0_10:0_38:2 | 1.184 | 0.009 |
| SM 34:1;O2 B | 1.182 | 0.014 |
| SL 20:1;O/36:0;O B | 1.179 | 0.011 |
| SL 15:0;O/36:0;O | 1.175 | 0.006 |
| TG 8:0_8:0_23:0 | 1.175 | 0.017 |
| TG 42:0\|TG 12:0_14:0_16:0 | 1.173 | 0.015 |
| SM 37:3;O3 | 1.167 | 0.005 |
| CAR 16:0 B | 1.167 | 0.029 |
| TG 44:1\|TG 12:0_14:0_18:1 | 1.166 | 0.013 |
| HexCer 17:0;O2/24:6;O B | 1.163 | 0.032 |
| Cer 41:0;O2\|Cer 18:0;O2/23:0 | 1.162 | 0.012 |
| TG 44:2\|TG 12:0_14:0_18:2 | 1.161 | 0.007 |
| LPC 12:0 | 1.161 | 0.015 |
| PC 36:3\|PC 18:1_18:2 | 1.159 | 0.025 |
| TG 56:8\|TG 16:0_18:2_22:6 | 1.158 | 0.011 |
| TG 54:6\|TG 16:0_18:2_20:4 | 1.157 | 0.015 |
| Cer 13:1;O3/38:2(2OH) | 1.157 | 0.029 |
| NAGly 20:4;O(FA 28:6) | 1.154 | 0.049 |
| SE 24:1;O4/26:1;O | 1.148 | 0.030 |
| Cer 25:1;O2/22:3;O | 1.147 | 0.017 |
| NAGly 26:0;O(FA 28:2) | 1.145 | 0.025 |
| TG 48:1\|TG 14:0_16:0_18:1 | 1.143 | 0.034 |
| DG 18:0_34:0 | 1.142 | 0.040 |
| TG 10:0_12:0_24:1 | 1.141 | 0.017 |
| TG 54:4\|TG 18:1_18:1_18:2 A | 1.138 | 0.022 |
| TG 8:0_10:0_38:8 | 1.134 | 0.001 |
| TG 46:2\|TG 12:0_16:0_18:2 | 1.134 | 0.016 |
| SL 16:0;O/36:0;O | 1.132 | 0.010 |
| Cer 13:2;O2/38:2 | 1.129 | 0.015 |
| TG 8:0_8:0_36:1 A | 1.126 | 0.018 |
| TG 8:0_9:0_36:3 B | 1.122 | 0.032 |
| TG 48:4\|TG 12:0_18:2_18:2 | 1.121 | 0.029 |
| Cer 12:0;O2/22:1;O | 1.118 | 0.036 |
| PC 38:6 A | 1.115 | 0.032 |
| HexCer 34:1;O3\|HexCer 18:1;O2/16:0;C | 1.11 | 0.030 |
| SM 12:1;O2/34:0 | 1.108 | 0.023 |
| CE 20:1 | 1.098 | 0.032 |
| Cer 12:1;O2/39:11;O2 A | 1.097 | 0.040 |
| TG 14:0_16:0_18:0 | 1.091 | 0.015 |
| Cer 12:0;O2/30:0;O | 1.088 | 0.027 |
| DG 39:6 | 1.086 | 0.000 |
| SL 13:0;O/36:0;O | 1.077 | 0.008 |
| DG 18:0_36:0 | 1.077 | 0.040 |
| TG 54:1\|TG 18:0_18:0_18:1 | 1.076 | 0.008 |
| TG 10:0_10:0_24:1 | 1.076 | 0.008 |
| DG 52:1 | 1.074 | 0.002 |
| TG 50:5\|TG 12:0_18:1_20:4 | 1.074 | 0.018 |

| | | |
|---|---|---|
| TG 50:4\|TG 14:0_18:1_18:3 | 1.074 | 0.050 |
| NAGly 16:2;O(FA 28:5) | 1.073 | 0.026 |
| PC O-8:0_26:1 | 1.072 | 0.009 |
| TG 8:0_8:0_34:3 | 1.072 | 0.011 |
| TG 48:3\|TG 12:0_18:1_18:2 | 1.064 | 0.018 |
| ST 28:1;O A | 1.06 | 0.020 |
| Cer 53:3;O2\|Cer 17:3;O2/36:0 | 1.057 | 0.013 |
| TG O-21:1_17:0_18:0 | 1.053 | 0.038 |
| TG 8:0_8:0_30:3 | 1.052 | 0.011 |
| TG 8:0_8:0_28:2 A | 1.051 | 0.027 |
| SL 22:1;O/36:0;O | 1.048 | 0.034 |
| TG 56:7\|TG 18:1_18:2_20:4 | 1.047 | 0.012 |
| TG 14:0_16:0_17:0 | 1.047 | 0.022 |
| TG 8:0_18:0_22:2 | 1.046 | 0.017 |
| NAPhe 20:4;O | 1.046 | 0.038 |
| TG 16:0_16:0_18:1 | 1.041 | 0.009 |
| TG O-50:0\|TG O-18:0_14:0_18:0 | 1.041 | 0.018 |
| TG 50:4\|TG 16:0_16:2_18:2 | 1.036 | 0.006 |
| TG 50:6\|TG 12:0_18:2_20:4 | 1.034 | 0.021 |
| PC 36:3\|PC 16:0_20:3 B | 1.034 | 0.031 |
| SE 24:1;O4/26:0;O | 1.033 | 0.040 |
| PC 32:0\|PC 16:0_16:0 C | 1.025 | 0.037 |
| ST 29:1;O B | 1.024 | 0.027 |
| TG O-48:0\|TG O-18:0_14:0_16:0 | 1.022 | 0.024 |
| AHexCer (O-12:0)12:2;O2/12:0;O A | 1.018 | 0.028 |
| SL 18:1;O/36:1;O | 1.013 | 0.022 |
| TG 18:0_14:1_20:2 | 1.011 | 0.037 |
| VAE 26:3 | 1.009 | 0.005 |
| ST 28:1;O B | 1.009 | 0.031 |
| TG 56:7\|TG 16:0_18:1_22:6 | 1.005 | 0.023 |
| PC 7:0_28:2 | 1.003 | 0.044 |
| Cer 25:1;O3/38:0(2OH) | 1.001 | 0.014 |
| TG 8:0_9:0_36:6 | 0.999 | 0.038 |
| Cer 23:1;O3/38:2(2OH) | 0.996 | 0.017 |
| TG 8:0_8:0_28:2 C | 0.995 | 0.026 |
| TG 52:4\|TG 16:1_18:1_18:2 | 0.988 | 0.042 |
| TG 50:6\|TG 12:0_16:0_22:6 | 0.983 | 0.011 |
| SL 12:1;O/35:1;O | 0.981 | 0.003 |
| TG 16:0_20:0_18:1 | 0.978 | 0.006 |
| TG 54:6\|TG 18:1_18:2_18:3 | 0.975 | 0.022 |
| TG 8:0_11:0_38:6 | 0.973 | 0.013 |
| TG 8:0_8:0_24:2 C | 0.969 | 0.005 |
| TG O-50:1\|TG O-16:0_16:0_18:1 | 0.969 | 0.031 |
| Cer 48:3;O2\|Cer 16:2;O2/32:1 | 0.964 | 0.009 |
| HexCer 34:1;O3\|HexCer 18:1;O2/16:0;C | 0.964 | 0.012 |
| CE 22:5 | 0.964 | 0.046 |
| CE 22:6 | 0.964 | 0.049 |
| TG 8:0_8:0_37:1 | 0.961 | 0.003 |
| ST 29:1;O A | 0.957 | 0.012 |

| | | |
|---|---|---|
| TG 8:0_8:0_38:7 | 0.955 | 0.004 |
| VAE 25:1 A | 0.954 | 0.033 |
| TG 16:0_18:1_18:1 | 0.952 | 0.017 |
| NAE 20:2 | 0.95 | 0.000 |
| SE 28:2/18:0 A | 0.95 | 0.028 |
| TG 14:0_18:1_18:1 | 0.949 | 0.023 |
| TG 8:0_12:0_38:3 | 0.947 | 0.021 |
| TG O-16:0_16:0_18:2 | 0.947 | 0.050 |
| Cer 41:1;O2\|Cer 18:1;O2/23:0 B | 0.946 | 0.009 |
| ST 27:2;O D | 0.944 | 0.022 |
| TG 11:0_15:0_19:1 | 0.942 | 0.019 |
| NAGly 28:0;O(FA 28:2) | 0.942 | 0.033 |
| TG 8:0_8:0_30:4 | 0.941 | 0.000 |
| Cer 50:2;O2\|Cer 20:0;O2/30:2 | 0.94 | 0.019 |
| TG 12:0_18:0_22:0 | 0.94 | 0.048 |
| Cer 23:1;O3/38:1(2OH) | 0.939 | 0.021 |
| TG 8:0_10:0_38:6 | 0.933 | 0.025 |
| TG O-52:0\|TG O-18:0_16:0_18:0 | 0.93 | 0.019 |
| LPC 18:1 C | 0.93 | 0.036 |
| TG 14:0_19:2_19:2 | 0.923 | 0.008 |
| PC 36:2\|PC 18:1_18:1 | 0.922 | 0.030 |
| HexCer 16:0;O2/19:0 A | 0.921 | 0.040 |
| TG O-13:0_13:0_18:0 | 0.92 | 0.034 |
| SL 13:1;O/34:6 | 0.919 | 0.030 |
| HexCer 18:1;O2/24:0 | 0.918 | 0.004 |
| TG O-54:2\|TG O-18:0_16:1_20:1 | 0.918 | 0.021 |
| SM 35:1;O3 | 0.915 | 0.003 |
| LPC 16:0 | 0.914 | 0.033 |
| TG 54:3\|TG 16:0_16:0_22:3 | 0.913 | 0.003 |
| Cer 12:2;O2/36:10;O2 | 0.912 | 0.009 |
| SE 24:1;O4/25:1;O B | 0.908 | 0.018 |
| Cer 17:0;O2/38:2 | 0.907 | 0.022 |
| TG O-20:0_12:0_18:0 | 0.906 | 0.049 |
| SL 15:3;O/32:6 B | 0.902 | 0.028 |
| LPC 18:2 B | 0.895 | 0.047 |
| LPC 18:2 A | 0.894 | 0.022 |
| SM 39:3;O3 | 0.892 | 0.025 |
| TG 8:0_11:0_38:3 | 0.89 | 0.004 |
| TG 8:0_9:0_36:3 A | 0.884 | 0.047 |
| TG 11:0_18:0_20:1 | 0.883 | 0.032 |
| LPC 30:3 | 0.881 | 0.045 |
| PC 36:1 | 0.88 | 0.025 |
| TG 18:0_20:1_18:3 | 0.88 | 0.031 |
| TG 8:0_11:0_38:4 | 0.872 | 0.028 |
| TG 12:0_16:0_18:3 | 0.87 | 0.008 |
| CE 21:0 | 0.869 | 0.004 |
| Cer 12:2;O2/38:11;O2 | 0.868 | 0.005 |
| CAR 16:1 | 0.866 | 0.023 |
| TG 8:0_8:0_21:0 | 0.861 | 0.021 |

| | | |
|---|---|---|
| Cer 17:0;O2/25:0;O | 0.859 | 0.031 |
| SL 13:1;O/32:6 | 0.857 | 0.000 |
| SM 12:1;O2/30:1 A | 0.857 | 0.037 |
| TG 8:0_8:0_29:1 | 0.856 | 0.008 |
| TG 17:0_15:1_17:1 | 0.855 | 0.016 |
| TG 8:0_8:0_32:0 | 0.854 | 0.011 |
| NAGly 18:1 B | 0.85 | 0.005 |
| TG 8:0_8:0_32:6 A | 0.849 | 0.026 |
| Cer 42:2;O2\|Cer 18:1;O2/24:1 A | 0.847 | 0.015 |
| Cer 42:1;O3\|Cer 18:1;O2/24:0;O | 0.846 | 0.035 |
| SL 13:1;O/36:3;O | 0.845 | 0.017 |
| LPC 18:1 A | 0.844 | 0.032 |
| MGDG 9:0_14:1 | 0.84 | 0.020 |
| HexCer 34:1;O2\|HexCer 18:1;O2/16:0 | 0.839 | 0.029 |
| TG 10:0_10:1_8:0;O2 | 0.838 | 0.016 |
| TG 17:0_13:1_17:1 | 0.833 | 0.027 |
| TG 8:0_9:0_38:0 | 0.827 | 0.002 |
| TG 20:0_18:1_18:1 | 0.824 | 0.004 |
| Cer 14:2;O3/8:0(2OH) | 0.822 | 0.012 |
| CE 20:3 | 0.818 | 0.030 |
| PC O-8:0_9:0 | 0.811 | 0.001 |
| TG 18:1_18:1_18:1 | 0.811 | 0.039 |
| Cer 13:2;O2/28:6 | 0.801 | 0.000 |
| PC 38:5 B | 0.787 | 0.030 |
| TG 8:0_8:0_26:1 | 0.787 | 0.045 |
| TG 8:0_26:1_24:3 | 0.784 | 0.041 |
| Cer 42:2;O2\|Cer 18:1;O2/24:1 B | 0.782 | 0.025 |
| ST 27:1;O B | 0.781 | 0.043 |
| TG 12:0_16:0_20:4 | 0.781 | 0.047 |
| TG 11:0_17:0_19:1 | 0.774 | 0.005 |
| SL 15:3;O/34:6 | 0.77 | 0.049 |
| DG 54:1 | 0.767 | 0.010 |
| LPC O-19:3 | 0.767 | 0.047 |
| C17-Sphingosine | 0.765 | 0.034 |
| TG 8:0_8:0_36:1 B | 0.761 | 0.017 |
| TG 16:0_16:0_19:1 | 0.755 | 0.032 |
| TG O-40:0\|TG O-18:0_10:0_12:0 | 0.752 | 0.032 |
| Cer 18:1;O3/38:1(2OH) | 0.751 | 0.001 |
| TG 17:0_15:1_19:1 | 0.742 | 0.046 |
| LPC 18:1 B | 0.731 | 0.048 |
| TG 8:0_11:0_38:1 | 0.73 | 0.003 |
| NAE 20:1 | 0.729 | 0.004 |
| CAR 17:1 | 0.726 | 0.030 |
| TG 8:0_9:0_38:2 | 0.724 | 0.003 |
| LPC 16:1 | 0.724 | 0.037 |
| HexCer 17:0;O2/26:3;O | 0.721 | 0.003 |
| TG 8:0_8:0_28:1 | 0.718 | 0.003 |
| SE 24:1;O4/25:1;O A | 0.717 | 0.019 |
| DG 52:2 | 0.717 | 0.037 |

| | | |
|---|---|---|
| Cer 12:0;O3/8:0(2OH) | 0.716 | 0.003 |
| TG 8:0_8:0_34:1 | 0.713 | 0.001 |
| TG 8:0_8:0_30:2 | 0.713 | 0.028 |
| SL 13:1;O/32:5 | 0.706 | 0.035 |
| LPC 20:4 A | 0.706 | 0.039 |
| NAGlySer 9:0;O(FA 26:4) | 0.705 | 0.022 |
| PC 6:0_32:6 | 0.702 | 0.015 |
| TG 8:0_8:0_18:0 | 0.697 | 0.038 |
| TG 13:0_18:0_20:0 | 0.694 | 0.025 |
| NAGly 22:3 | 0.694 | 0.035 |
| TG 18:0_18:1_18:1 | 0.688 | 0.017 |
| LPE 20:4 | 0.688 | 0.043 |
| Cer 12:1;O2/39:11;O2 B | 0.682 | 0.016 |
| SE 28:2/19:0 B | 0.675 | 0.004 |
| TG 8:0_8:0_38:0 B | 0.672 | 0.022 |
| NAE 15:1 | 0.671 | 0.041 |
| NAOrn 19:0;O A | 0.667 | 0.034 |
| TG 16:0_18:0_18:2 | 0.664 | 0.001 |
| TG 8:0_8:0_30:1 B | 0.663 | 0.012 |
| Cer 12:1;O3/35:0(2OH) | 0.662 | 0.042 |
| Cer 22:1;O3/38:0(2OH) | 0.657 | 0.050 |
| Cer 42:1;O2\|Cer 18:1;O2/24:0 C | 0.654 | 0.003 |
| TG 8:0_14:0_38:5 | 0.651 | 0.014 |
| NAGly 20:1 | 0.651 | 0.050 |
| CE 20:2 | 0.643 | 0.035 |
| ST 27:1;O I | 0.632 | 0.015 |
| Cer 14:0;O3/38:0(2OH) | 0.631 | 0.039 |
| TG 8:0_8:0_36:0 | 0.628 | 0.049 |
| Cer 16:3;O3/32:6(2OH) | 0.625 | 0.010 |
| Cer 12:1;O3/25:1(2OH) | 0.625 | 0.030 |
| Cer 40:0;O2\|Cer 18:0;O2/22:0 | 0.625 | 0.047 |
| TG 54:4\|TG 18:1_18:1_18:2 B | 0.621 | 0.017 |
| SE 28:2/24:4 | 0.619 | 0.006 |
| NAGly 15:3 A | 0.617 | 0.031 |
| HexCer 16:0;O2/22:0;O | 0.612 | 0.033 |
| NAOrn 19:0;O B | 0.612 | 0.044 |
| PC 36:5\|PC 16:0_20:5 B | 0.611 | 0.036 |
| TG 18:0_20:1_18:2 | 0.6 | 0.043 |
| NAPhe 17:4 | 0.594 | 0.009 |
| NAGly 13:0;O(FA 9:0) | 0.593 | 0.025 |
| PE O-8:0_15:1 | 0.592 | 0.048 |
| DG 53:7 | 0.589 | 0.008 |
| Cer 12:2;O2/15:3 | 0.589 | 0.015 |
| TG 12:0_18:0_18:2 | 0.589 | 0.031 |

**Table S3. Antibodies used for flow cytometry analysis.**

| Marker | Clone | Company | Conjugate | Dilution | Catalogue |
|---|---|---|---|---|---|
| CD45 | 30-F11 | BioLegend | PE-Cy7 | 200 | 103114 |
| CD4 | RM4-5 | Biolegend | Pacific Blue | 200 | 100531 |
| F4/80 | BM8 | BioLegend | APC | 200 | 123116 |
| Siglec-F | S17007L | BioLegend | PE | 200 | 155506 |
| CD117 | ACK2 | BioLegend | BV421 | 200 | 135124 |
| FcεRI | MAR-01 | BioLegend | PE-Cy7 | 200 | 134308 |
| CD107a | 1D4B | BioLegend | PerCP/Cy5.5 | 200 | 121626 |
| CD63 | NVG-2 | BioLegend | APC | 200 | 143906 |

**Table S4. Primer sequences of mouse genes used for quantification of mRNAs by real-time PCR.**

| Gene | Forward (5′–3′) | Reverse (3′–5′) |
|---|---|---|
| *Srebf1* | ATCGCAAACAAGCTGACCTG | AGGCCAGATCCAGGTTTGAG |
| *Fasn* | ACTGACTCGGCTACTGACAC | GAGTTGAGCTGGGTTAGGGT |
| *Cd36* | GGCTAAATGAGACTGGGACC | ACATCACCACTCCAATCCCA |
| *Elovl6* | CCGGAAGTTTGCCATGTTCA | AAGTGGGAGTAGCACTGGTC |
| *Dgat2* | CCGATGGGTCCAGAAGAAGT | GTGGTGATGGGCTTGGAGTA |
| *Rxra* | CATGCAGATGGACAAGACGG | GACGCATACACCTTCTCCCT |
| *Mttp* | TCCGTCGAGTTCTCAAGGAG | GTCAAGGCTGTATGTGGACG |
| *Acly* | AGGTCACCATCTTTGTCCGA | GGCCGTCATGTGAGTTTCTG |
| *Acc1* | GTTTGGTCGTGACTGCTCTG | TTGGCAAGTTTCACTGCACA |
| *Chrebpa* | CGACACTCACCCACCTCTTC | TTGTTCAGCCGGATCTTGTC |
| *Lipin1* | CCTTCTATGCTGCTTTTGGGAACC | GTGATCGACCACTTCGCAGAGC |
